# Establishing *Vibrio natriegens* as a high-performance host for acetate-based poly-3-hydroxybutyrate production

**DOI:** 10.1101/2025.06.23.661028

**Authors:** Roland J. Politan, Simona Della Valle, Luke Pineda, Jitendra Joshi, Christian Euler, Gavin Flematti, Georg Fritz

## Abstract

Acetate can be a sustainable and renewable carbon source which holds significant promise for biotechnological production but is underutilized industrially due to limited microbial efficiency. *Vibrio natriegens*, recognized for exceptionally fast growth rates, represents a compelling host for developing efficient acetate-based bioprocesses. In this study, adaptive laboratory evolution significantly enhanced *V. natriegens*’ ability to grow on acetate as the sole carbon source, achieving an 89% increase in growth rate. Genetic and transcriptomic analyses revealed key adaptations improving acetate uptake and metabolism via increased salt tolerance, boosted Pta/AckA pathway activity, and rewired quorum sensing. Further metabolic engineering and bioprocess optimization enabled the evolved strain to reach high cell densities and efficiently convert acetate into the bioplastic poly-3-hydroxybutyrate (PHB), with productivities up to 0.27 g/L/h and PHB accumulation reaching 45.66% of cell biomass. These advances position *V. natriegens* as a highly promising microbial platform for sustainable, scalable, and cost-effective biomanufacturing using acetate as green feedstock.

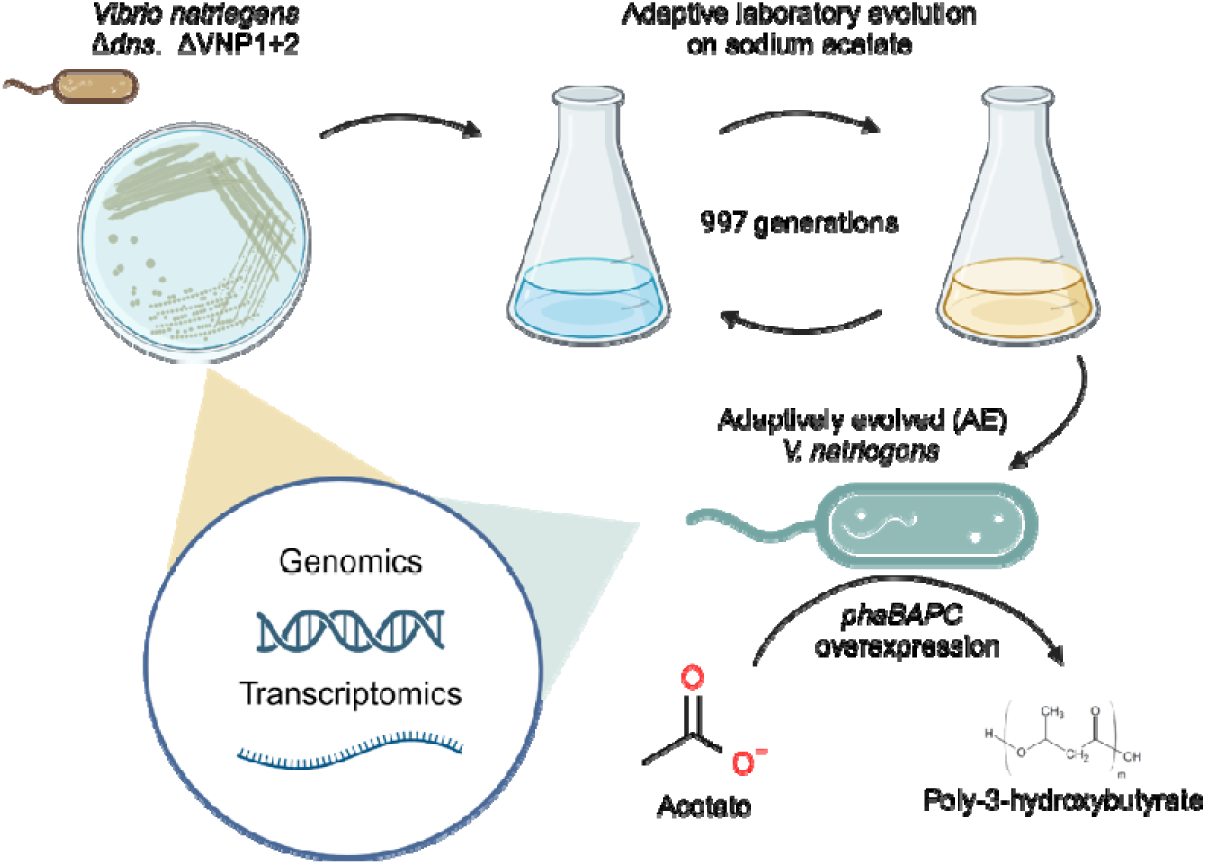

## Introduction

Acetate is increasingly recognized as a sustainable, low-cost carbon source for biomanufacturing. Unlike conventional sugars (e.g. glucose) or glycerol, acetate can be derived from waste streams or C□ gases via processes such as CO□ electrocatalysis, anaerobic digestion, or syngas fermentation^1–3^. Indeed, acetate production can leverage abundant carbon dioxide or methane, yielding a liquid product that is easier to store and handle than gaseous substrates^1^. Economically, acetate is notably cheaper than glucose (on the order of ∼$300–450 versus ∼$500 per ton^1^), and because nearly half of bioprocess cost is often attributed to feedstock, substituting sugar with acetate from non-food biomass or waste offers both cost and sustainability advantages^4,5^. Furthermore, acetate is highly soluble, and its microbial consumption naturally increases culture pH, simplifying pH control through acetate feeding in large-scale fermentations. However, while acetate’s low cost, renewable sourcing, and process-friendly properties make it an attractive feedstock for green bioproduction, the efficient microbial utilization of acetate remains a challenge, and many established bioproduction hosts exhibit low growth rates, limited acetate uptake, or lack the necessary genetic tools for optimization^2,5^.

*Vibrio natriegens* is emerging as a next-generation microbial chassis^6^ with unique attributes that suit acetate-based bioproduction. This marine bacterium boasts an exceptionally fast growth rate – its doubling time can be under 10 minutes, over twice as fast as *Escherichia coli* even in defined medium^7,8^. Correspondingly, *V. natriegens* exhibits very high substrate uptake rates (often >2× those of *E. coli*), which can translate into higher volumetric productivity^9,10^. In addition to its speed, *V. natriegens* is inherently halotolerant, allowing non-sterile or open fermentations, reducing sterilization costs^11^. The bacterium is also non-pathogenic and considered BSL-1, with no known human infections^12^, making it safe for industrial use. Crucially, *V. natriegens* has a broad metabolic capability – it natively grows on glycerol, arabinose, galactose and other substrates – indicating versatility in feedstock utilization^8,10^. A suite of genetic tools and omics resources has rapidly been developed for this organism^8,9,13–26^ making it tractable for engineering. These features – ultrafast growth, high uptake, salt resilience, and genetic accessibility – thus position *V. natriegens* as a superior host compared to traditional bacteria for industrial biomanufacturing.

Engineered *V. natriegens* strains have already demonstrated industrial potential across a broad product spectrum, often outperforming *E. coli* and other industrial strains in volumetric productivity. These include amino acids^8,27^, polyols^28,29,30^, pigments (e.g., melanin^31^, indigoidine^32^, β-carotene and violacein^10^), and organic acids such as succinate^33^ and 3-hydroxypropionate^34^. Notably, productivity gains of 1.5–3× for compounds like 2,3-butanediol^28,29^ and up to 13× for alanine^8^ have been reported. This breadth of successful applications highlights *V. natriegens’* versatility as a microbial platform and suggests that advances in acetate metabolism could generalize to other value-added products.

Among the many products derived from acetyl-CoA, we selected poly-3-hydroxybutyrate (PHB) as a strategic model compound. PHB is a biopolymer, which is naturally accumulated by many microorganisms as an intracellular carbon storage compound. PHB is synthesized through a three-step enzymatic pathway: acetyl-CoA is condensed by PhaA, reduced by PhaB, and polymerized by PhaC, resulting in its accumulation within intracellular granules. As a biodegradable polyester, PHB exhibits mechanical properties comparable to common plastics such as polypropylene and polyethylene^35^, yet it fully degrades under natural conditions^36^. These attributes make PHB a strong candidate for sustainable alternatives in packaging, agriculture, and medical applications currently reliant on petrochemical plastics. However, PHB production is highly sensitive to feedstock costs, which typically account for over 50% of total production expenses^37,38^. Notably, *V. natriegens* has already demonstrated the ability to produce PHB efficiently from crude glycerol under non-sterile conditions, achieving a productivity of 0.052 g/L/h and a final titer of 2.5 g/L^11^. It has also been engineered to produce the copolymer poly(3-hydroxybutyrate-co-lactate) from glucose, combining the transparency of polylactic acid with the impact and heat resistance of PHB^39^. As a proficient acetotroph and natural PHB producer, *V. natriegens* is uniquely poised to deliver on the economic and environmental advantages of using acetate as a feedstock for PHB production.

Despite this potential, a comprehensive understanding of how *V. natriegens* metabolizes acetate remains lacking. Little is known about the enzymes, regulatory nodes, and carbon flux distribution supporting acetate assimilation in this organism, or how its metabolism may be optimized for acetate-based bioproduction. Additionally, the dedicated metabolic models for this condition have not yet been developed. These knowledge gaps hinder rational pathway engineering and process optimization for acetate-based bioproduction.

Here, we address this challenge by applying an integrated approach to establish *V. natriegens* as a robust host for PHB production from acetate. Our strategy combines adaptive laboratory evolution (ALE) to enhance growth and tolerance, transcriptomics and metabolic flux modelling to reveal regulatory and metabolic shifts, targeted genetic engineering to modulate key pathways, and fed-batch bioprocessing to enable sustained, scalable acetate conversion. By illuminating how *V. natriegens* reallocates carbon flux under selective pressure and controlled conditions, we provide novel insights into acetate metabolism and offer a blueprint for developing high-performance microbial platforms for sustainable, acetate-based biomanufacturing.

## Results

### Characterization of wildtype *V. natriegens* growth on acetate

Acetate typically occurs as a byproduct of overflow metabolism through the reversible phosphate acetyltransferase (Pta) and acetate kinase (AckA) pathway, facilitated by the bidirectional acetate permease (ActP)^40^. As the primary carbon source depletes during later stages of growth, cells undergo an “acetate switch” via the catabolite repressor protein (CRP), triggering the upregulation of the high-affinity Acetyl-CoA synthetase (ACS) pathway to convert acetate back into acetyl-CoA in a single enzymatic reaction^40^. At higher environmental concentrations of acetate, the low-affinity Pta/AckA pathway is utilized to synthesize acetyl-CoA from acetate with acetyl phosphate as an intermediate^40^.

#### Acetate metabolism in V. natriegens.

While *V. natriegens* has been demonstrated to grow on acetate as a sole carbon source^8,34^, its associated growth kinetics, metabolic pathways, and concomitant fluxes have not been described. *V. natriegens* DSM 759 is predicted to possess one copy of the *actP* (RS14445, chromosome 1) gene, two copies for each of the *acsA*, *pta*, and *ackA* genes, respectively, referred to as *acsA1* (RS14410)*, pta1* (RS03410)*, ackA1* (RS03415) (chromosome 1) and *acsA2* (RS21410)*, pta2* (RS22305)*, ackA2* (RS21805) (chromosome 2), though only the copies in chromosome 1 are considered core genes^12,14^. Specifically, only the *pta1-ackA1* pair is organized in an operon, while the remaining genes are scattered throughout the genome.

To grow effectively on acetate, carbon flux must be finely balanced between energy production via the TCA cycle and carbon conservation through the glyoxylate shunt. This balance is typically regulated by the IclR repressor, which inhibits the transcription of *aceA* and *aceB*, encoding isocitrate lyase and malate synthase, respectively^40^. Interestingly, *V. natriegens* contains two *iclR* homologues in its genome. The glyoxylate shunt is encoded by a single *aceA-aceB1* operon (RS10725, RS10730; chromosome 1) and an additional *aceB2* copy (RS17175; chromosome 2). These genetic features involving multiple copies of most acetate metabolic genes might contribute to *V. natriegens’* natural efficiency for rapid acetate utilization.

To predict how acetate can be optimally metabolized through the different pathways encoded on the *V. natriegens* genome, we conducted flux balance analysis using the iLC858 genome-scale metabolic model (GSMM) (**Fig. 1A**) ^15^. The model steady-state flux distribution predicted that the Pta/AckA pathway is favored over the ACS pathway for acetate assimilation, likely due to the former being an energetically more efficient route (converting ATP to ADP) than the latter (converting ATP to AMP). Subsequently, acetyl-CoA is metabolized through the TCA cycle, where the model predicts the flux to split almost equally between the TCA cycle (45.8%) and the glyoxylate cycle (54.2%). Most of the flux appears to be recycled in the TCA cycle, with only 14.9% of the flux predicted to exit from oxaloacetate to phosphoenolpyruvate (PEP).

**Figure 1.**
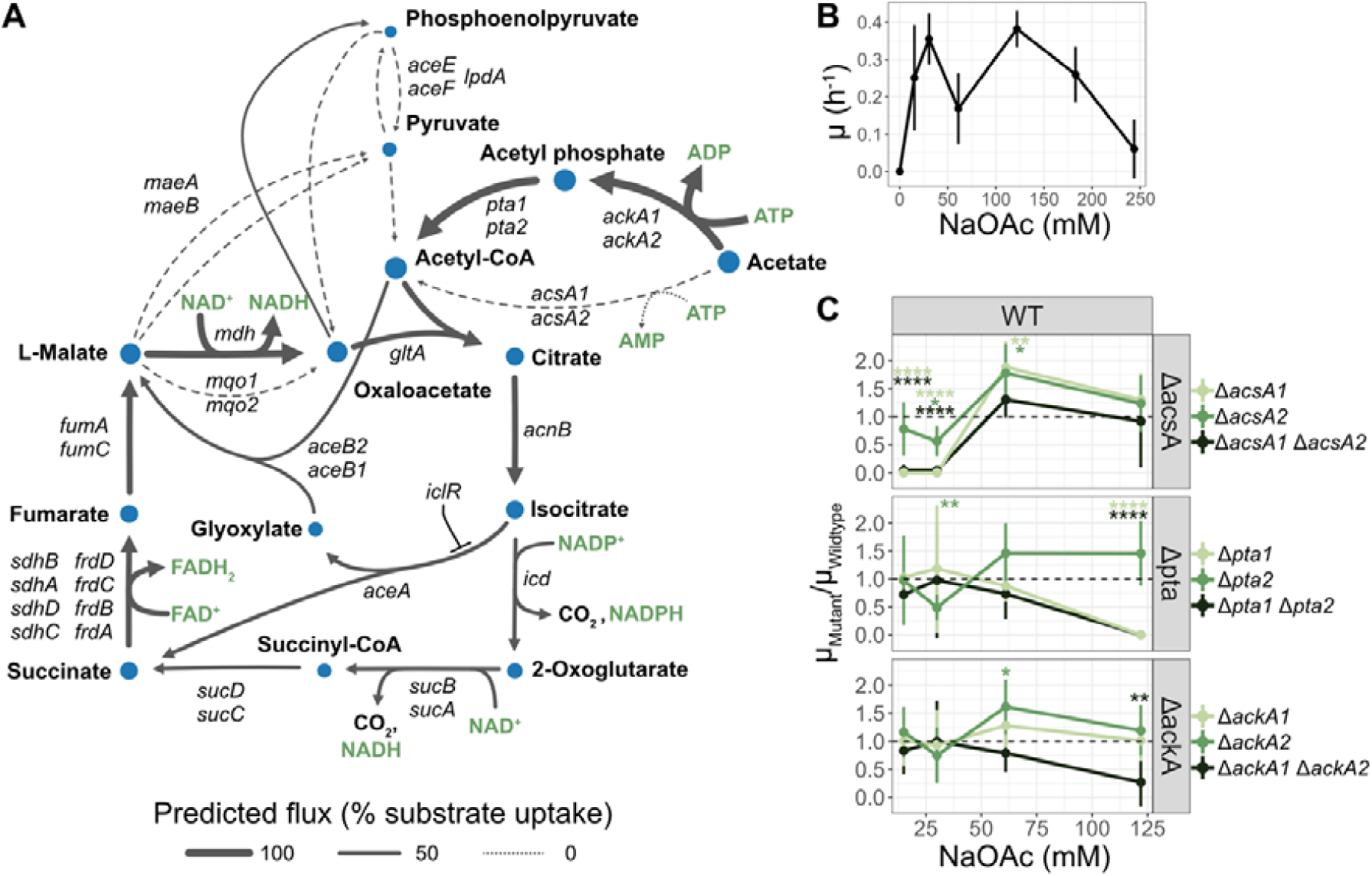
Elucidation of the acetate metabolism pathway in *V. natriegens* through genome scale metabolic modeling and knockout studies. Locus tags are provided in **Table S1**. A) Prediction of acetate flux through central carbon metabolism using GSMM iLC858. B) Growth rate of wildtype *V. natriegens* on different concentrations of NaOAc in MOPS2 media at pH 7.2 (*n* = 6). C) Growth kinetic study of *V. natriegens* mutants related to acetyl-CoA production from acetate (*n* = 6). The growth rate of the mutants was normalized to the wildtype strain. Data and error bars represent mean and standard deviations from independent biological replicates. Statistical significance for C) was performed with a one-sample t-test. (*p< 0.05, **p< 0.01, ***p<0.001, ****p<0.0001)

### Growth kinetics and tolerance towards acetate

Next, we investigated the growth kinetics of *V. natriegens* across different concentrations of neutralized sodium acetate (NaOAc) in MOPS-based minimal media supplemented with 2 g/L (342 mM) NaCl (MOPS2 media, *see* Methods). As anticipated, *V. natriegens* successfully utilized NaOAc as the sole carbon source without the need for complex nitrogen sources (**Fig. 1B**). The maximal growth rates were observed at 31 and 122 mM NaOAc, with rates of 0.38 ± 0.08 h[¹ and 0.39 ± 0.06 h[¹. Notably, *V. natriegens* tolerated up to 244 mM NaOAc, although growth was markedly slower at this concentration, with a rate of 0.06 ± 0.08 h[¹. Intriguingly, a significant reduction in growth rate was observed at an intermediate NaOAc concentration of 61 mM, suggesting a potentially suboptimal transition between the different acetate assimilation pathways in the wild-type strain.

To further elucidate the role of and interplay between the ACS and Pta/AckA pathways, we performed single and double knockouts in *acsA1/acsA2*, *pta1/pta2*, and *ackA1/ackA2*, respectively, and measured the growth rates of the resulting strains at different NaOAc concentrations (**Fig. 1C**). Interestingly, a *V. natriegens* Δ*acsA1* strain was unable to grow on low acetate concentrations (≤ 31 mM), whereas a Δ*pta1* strain failed to grow at high acetate concentrations (≥ 122 mM). In contrast, both the Δ*acsA2* and the Δ*pta2* mutants did not display highly significant changes in growth compared to the wildtype, and the respective double mutants (Δ*acsA1*Δ*acsA2* and Δ*pta1*Δ*pta2)* phenocopied the single mutants of Δ*acsA1* and Δ*pta1*, respectively. This suggests that *acsA1* is essential for acetate metabolism at low concentrations, while *pta1* plays a crucial role at higher acetate levels, whereas *acsA2* and *pta2* are dispensable. In contrast, at high NaOAc concentrations (122 mM), the knockout of either *ackA1* or *ackA2* alone did not affect the growth rate significantly, while a double *ackA* knockout displayed a markedly reduced growth rate, suggesting functional redundancy between these genes. This suggests that the ACS pathway may compensate for the loss of *ackA* in the presence of *pta*. As anticipated, all mutants, including the double knockouts, were capable of growth at 61 mM NaOAc, confirming the concurrent activity of both pathways at this concentration. Intriguingly, both *acsA1* and *acsA2* single mutants exhibited approximately a 75% increase in growth rate at 61 mM NaOAc, restoring growth rates to the peak levels observed for 31 and 122 mM NaOAc (**Fig. 1B**). However, this significant increase was absent in the double *acsA* knockout mutant, suggesting a complex regulatory interplay between the ACS and Pta/AckA pathways in *V. natriegens’* acetate metabolism.

This is a UWA Official document authorised or intended for public access.

### Adaptive laboratory evolution for increased acetate utilization

To enhance *V. natriegens’* growth on and utilization of acetate, we performed adaptive laboratory evolution (ALE) across three independent DSM 759 Δ*dns* ΔVNP1+2 (“WT”) cell lines for approximately 1,000 generations with 122 mM NaOAc as the sole carbon source. This concentration was selected to enable daily 1:1,000 dilutions of the evolving cell culture into fresh media and to promote acetate assimilation via the Pta/AckA pathway, predicted to be the more energetically favorable metabolic route. Following the ALE, the adaptively evolved (AE) cultures were plated on MOPS2 minimal media with 122 mM NaOAc as the sole carbon source and two of the largest colonies from each independent cell line were selected for further characterization.

#### Evolved strains display higher growth rate and tolerance to NaOAc

To measure the growth rates of the AE strains, we grew them in a microplate reader under varying concentrations of NaOAc. The growth rates of the AE strains did not differ significantly from the WT strain at concentrations up to 31 mM NaOAc (0.38 ± 0.08 h^-1^) (**Fig. 2A** and **Fig. S1**). However, the growth rates of AE2 and AE3 are significantly higher at elevated NaOAc concentrations, achieving 0.72 ± 0.06 h^-1^ and 0.67 ± 0.15 h^-1^, respectively, at 122 mM NaOAc. This is approximately double the growth rate of the WT strain at the same concentration, which is also the concentration at which the AE strains evolved. Given that the ACS pathway is predominantly used at lower acetate concentrations and the Pta/AckA pathway at higher concentrations, these results suggest that the AE strains may have adapted to utilize higher acetate concentrations via the Pta/AckA pathway. Furthermore, while the WT strain and AE1 were unable to grow at 244 mM NaOAc, AE3 displayed a growth rate of 0.45 ± 0.24 h^-1^, followed by AE2 at 0.16 ± 0.12 h^-1^. This indicates that the AE2 and AE3 strains have developed higher tolerance to either sodium, acetate or both.

**Figure 2.**
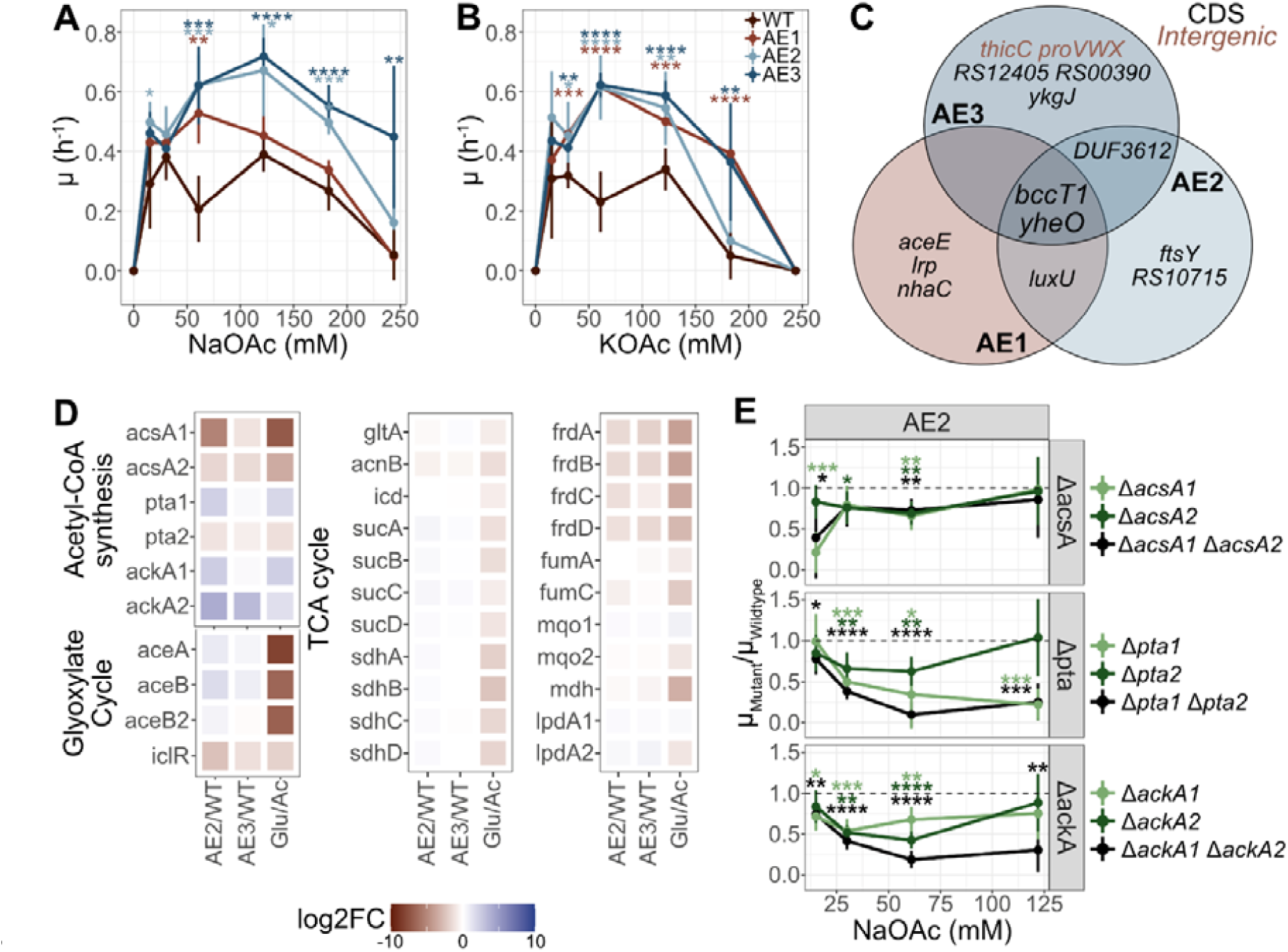
Characterization of the adaptively evolved *V. natriegens* strains. A-B) Growth rate of *V. natriegens* before and after adaptive evolution during steady state growth by looking at the average slope of a linear regression model between plate reader OD_600_ of 0.025-0.1 (n = 6) with the respective standard deviation A) on NaOAc, B) on KOAc (n = 6). C) Venn diagram of the commonly mutated genes during the evolution process. D) Transcriptomic response of WT grown on 28 mM glucose compared to 61 mM NaOAc and the fastest growing AE strains compared to WT during growth on 61 mM NaOAc (n = 3). E) Verification of the preferred acetyl-CoA synthesis pathway after adaptive evolution through a knock-out study. Data and error bars represent mean and standard deviations from independent biological replicates (n = 6). Statistical analysis in A) and B) were performed with unpaired two-sided t-test and in E) with one-sided t-test for *acsA, pta, and ackA* mutants relative to the parental AE2 strain. (*p< 0.05, **p< 0.01, ***p<0.001, ****p<0.0001)

To test whether the observed increase in growth rate and tolerance towards NaOAc was caused by a mere increase in sodium tolerance, we repeated the experiment with potassium acetate (KOAc) (**Fig. 2B**, kinetics in **Fig. S1**). As observed with NaOAc, both the wild-type and AE strains showed peak growth rates at intermediate KOAc concentrations (61–122 mM), reaching maximal rates comparable to those on NaOAc. The only exception is strain AE1, which achieved similar growth rates than AE2 and AE3 when grown on KOAc, while it was markedly slower when grown on NaOAc (cf. **Fig. 2A**). These results further support that the AE strains have indeed evolved to utilize and/or tolerate acetate as a carbon source, and that AE2 and AE3 have developed a higher tolerance towards elevated sodium concentrations compared to AE1.

### Genomic adaptations of evolved strains

To uncover the genetic basis of the phenotypic variations in AE strains, we sequenced two colonies from each independent lineage and re-sequenced the ancestral WT strain to avoid false positives. This analysis revealed mutations in 12 coding sequences and 2 intergenic regions. (**Fig. 2C, Table S2**). We investigated some of the mutations that are observed in more than one independent cell line, indicating a potential adaptive advantage or functional relevance across different evolutionary trajectories. Particularly, every lineage consistently targeted RS06115, encoding a BCCT family bidirectional transporter for compatible solutes^41–43^, and *yheO* (RS14910, see **Fig. S2A, B**), a transcriptional regulator within the *tusDCB* operon that encodes a sulfur transferase complex involved in tRNA modification^44^. Two independent cell lines had a frameshift mutation in one of the quorum sensing genes, *luxU* (RS03340) and a gene encoding a DUF3612 domain-containing protein (RS02055, see **Fig. S2C, D**), which likely functions as a transcriptional regulator based on its homology to MerR. These mutations underscore the potential significance of these genes in the adaptive responses of these strains.

#### Transcriptomics reveals evolutionary adaptations in AE strains

As multiple transcriptional regulators were mutated, we employed a transcriptomics approach in two of the fastest-growing AE strains to elucidate their phenotype. As a reference we analyzed the transcriptome of the WT during fast growth on 5 g/L (28 mM) glucose compared to slow growth at 5 g/L (61 mM) NaOAc (Glu/Ac), as well as the transcriptome profiles of AE2 and AE3 in comparison to WT (AE2/WT, AE3/WT) during steady-state growth at 61 mM NaOAc. Global transcriptomic analysis revealed significant metabolic shifts in the AE strains relative to WT (**Fig. 3A, Table S3**), with AE2 specifically showing high transcriptomic similarity to the WT during growth on glucose as a sole carbon source (**Fig. 3B**), with 154 commonly upregulated and 510 downregulated genes (**Fig. 3C**). Pathways such as fatty acid degradation, microbial metabolism in diverse environments, and valine, leucine, and isoleucine degradation were commonly suppressed between AE2, AE3 (during growth on acetate) and WT (during growth on glucose) (**Fig. 3D-F**). In contrast, pathways including ribosome biogenesis, aminoacyl-tRNA biosynthesis, the one-carbon pool by folate, and arginine biosynthesis were commonly activated (**Fig. 3D-F**). These changes suggest an adaptation in AE2 and AE3 towards faster growth and increased protein synthesis, optimized resource utilization, and a focused metabolic response to harsh environmental conditions, similar to the adaptation of the WT during growth on glucose.

**Figure 3.**
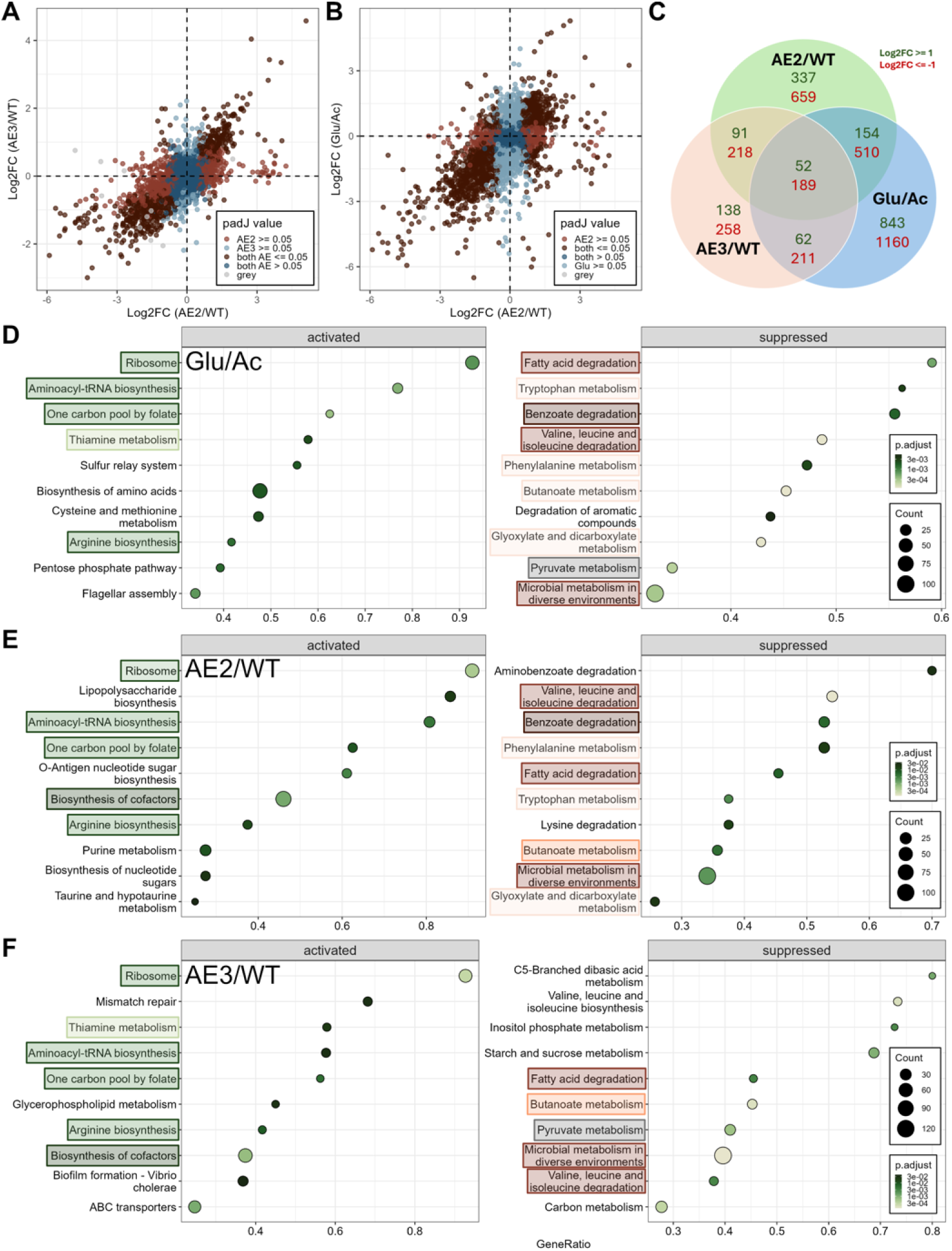
Global transcriptomic analysis performed in this experiment. A) Scatterplot of differentially expressed gene (DEG) in AE2/WT compared to AE3/WT growing in 61 mM NaOAc MOPS2. B) Scatterplot of DEG in AE2/WT (5 g/L or 61 mM NaOAc) compared to Glu/Ac (WT growing on 5 g/L or 28 mM glucose compared to 5 g/L NaOAc). C) Number of commonly regulated genes across the different conditions. D-F) KEGG enrichment analysis of the DEG of D) Glu/Ac, E) AE2/WT, F) AE3/WT. Wald test was used to obtain the p-values.

To better understand how adaptive evolution enhanced growth on NaOAc, we analyzed transcriptomic changes in the acetyl-CoA synthesis pathway (**Table S3**). In the WT strain, acetate metabolism appears to favor the ACS pathway, as indicated by strong upregulation of *acsA1* and *acsA2* (log2FC = 7.1 and 3.5) and downregulation of the Pta/AckA pathway genes *pta1*, *ackA1*, and *ackA2* (log2FC = –1.9, –2.2, and –1.5, respectively) when grown on acetate versus glucose. In contrast, the evolved strains AE2 and AE3 showed a consistent downregulation of *acsA1*, *acsA2*, and *pta2* during growth on acetate compared to WT (log2FC: AE2 = –5.2, –1.7, –1.1; AE3 = –1.2, –1.5, –0.7). Both strains exhibited strong upregulation of the monocistronic *ackA2* gene (log2FC = 4.0 in AE2; 3.4 in AE3), while only AE2 showed upregulation of the *pta1-ackA1* operon (log2FC = 2.2 and 2.4, respectively), indicating a more complete activation of the Pta/AckA pathway for acetate assimilation. These transcriptional changes suggest that AE2, in particular, evolved to preferentially utilize this more energy-efficient route, in line with predictions from the genome-scale metabolic model (**Fig. 1A**).

To verify this observation, we performed targeted knockouts of the *acsA* and *pta-ackA* genes in AE2 (**Fig. 2E**), analogous to the approach used for the WT strain in the previous section. In contrast to the WT strain, disrupting any gene within either the Pta/AckA or the ACS pathway resulted in a significantly reduced steady-state growth rate at 61 mM NaOAc, indicating that both pathways are simultaneously active under these conditions. Remarkably, the AE strains retained a low growth rate even with double knockouts of either *acsA* or *pta* at low or high concentrations of NaOAc. This resilience likely stems from the increased tolerance to NaOAc and/or the compensatory expression of the alternative metabolic pathway.

We subsequently analyzed transcriptional and metabolic adaptations in the TCA cycle (**Fig. 2D, F**). Most genes in the TCA cycle are downregulated on glucose compared to acetate, indicating a higher demand to regenerate energy and cofactors through the TCA cycle when acetate is the sole carbon source. After adaptive evolution, the expression of most TCA cycle genes remains unchanged, except for the *frd* operon, which encodes fumarate reductase—an enzyme that uses FADH[to convert fumarate to succinate. Its downregulation likely helps conserve FADH[and redirects metabolic flux toward malate, enhancing the efficiency of the TCA cycle and supporting improved energy balance during growth on acetate.

We then examined the downstream steps of acetate metabolism via the glyoxylate cycle (**Fig. 2D, F**). As expected, the WT strain showed strong upregulation of glyoxylate cycle genes during growth on acetate compared to glucose (log2FC = 8.15 for *aceA*, 6.53 for *aceB1*, and 6.77 for *aceB2*). After adaptive evolution, the AE strains exhibited further upregulation of these genes relative to WT during growth on NaOAc (log2FC = 1.00 and 1.66 for *aceA* and *aceB1* in AE2; 0.51 and 0.81 in AE3). This enhanced expression correlated with downregulation of one of the two *iclR* repressor genes (RS22730) in both AE strains (log2FC = –2.54 in AE2, –1.31 in AE3). These changes suggest that adaptive evolution promoted a shift toward greater carbon conservation through increased glyoxylate cycle activity, likely improving acetate assimilation and biomass production.

#### Characterization of the bccT1 mutation

A striking observation was that the *bccT1* gene was independently mutated in all three evolved cell lines. Proteins in the betaine/choline/carnitine transporter (BCCT) family mediate the bidirectional transport of compatible solutes across the cytoplasmic membrane, coupled to the symport of two H^+^ or sodium ions^43^. *V. natriegens* encodes seven *bccT* gene copies–two on chromosome 1 and five on chromosome 2– highlighting the importance of compatible solute homeostasis in this halophilic bacterium. Among these, BCCT1 shares the highest homology with *Vibrio parahaemolyticus* VP1456 (**Fig. S3, S4**), a trimeric membrane protein known for sodium-dependent transport of glycine betaine, proline, choline, and ectoine^42^. Notably, VP1456 and BCCT1 are the only BCCT transporters shown to mediate ectoine uptake in *V. parahaemolyticus*^42^ and *V. natriegens*^45^, respectively.

To evaluate the functional impact of the mutations, we performed *in silico* protein structure analysis of BCCT1 using AlphaFold3 (**Fig. 4A, B**). While none of the mutations disrupted the conserved ligand- or sodium-binding sites, AE2 harbored a D452V substitution that likely destabilizes the trimer by disrupting a key hydrogen bond between Q466 and R467. The deletions in AE1 and AE3 clustered at the multimerization interface in transmembrane helices 6 and 7, further supporting impaired trimer assembly and transporter function (**Fig. 4 A, B**).

**Figure 4.**
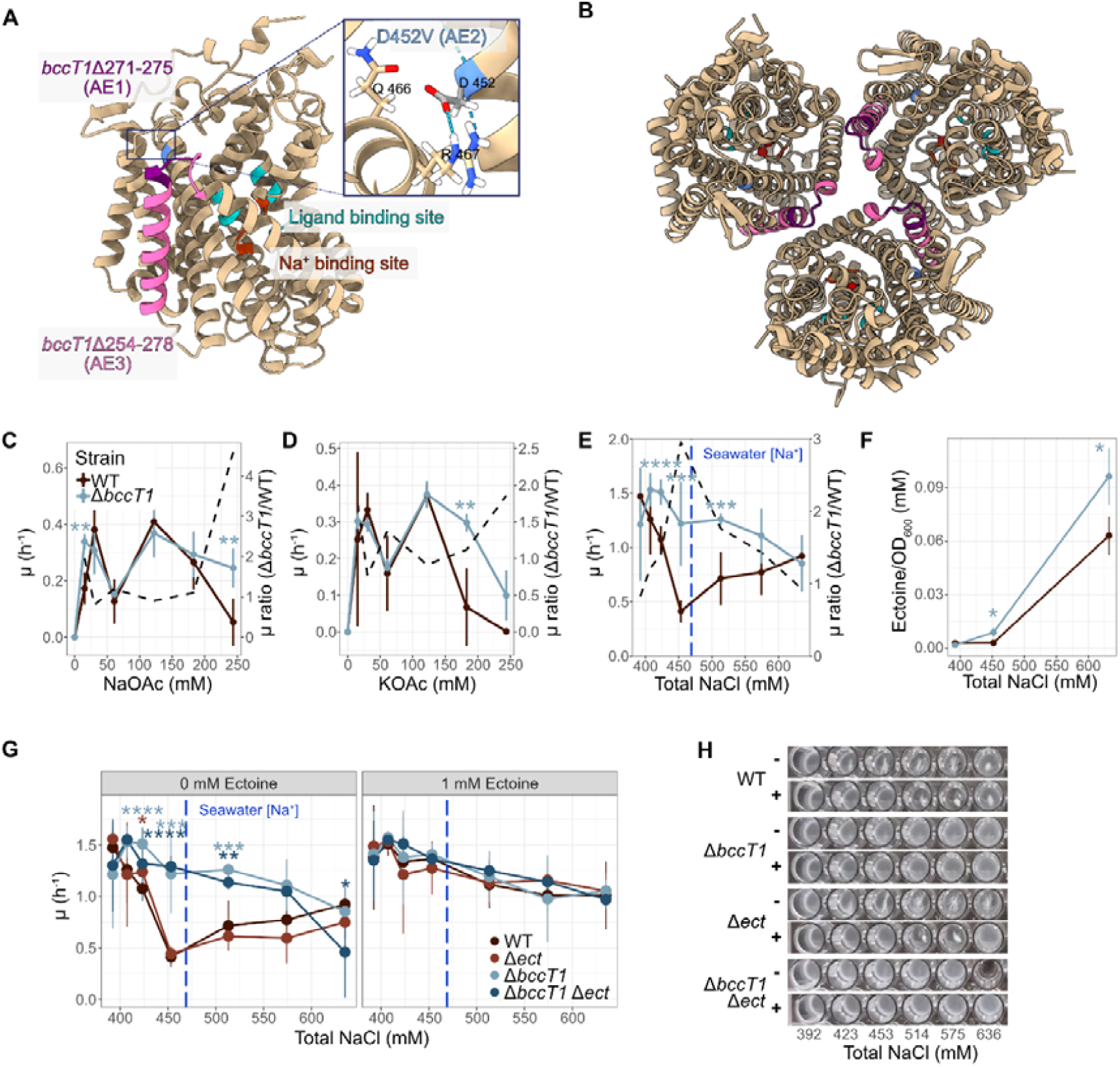
Phenotypic characterization of the *bccT1* mutation. A) Visualization of the BCCT1 protein prediction by AlphaFold3 from the side, B) from a top view as a trimer. C-E) Growth rate of the Δ*bccT1* mutant on defined MOPS2 media supplemented C) with increasing NaOAc titration measured at OD_600_ = 0.025 - 0.1 (n = 6), D) on KOAc titration (n = 4), E) on 28 mM glucose with increasing sodium titration measured at OD_600_ = 0.015 - 0.1 (n = 8). F) Intracellular ectoine levels of the Δ*bccT1* mutant measured using the HPLC and normalized by the OD_600_ (n = 3). G) Comparison of the growth rate between the Δ*bccT1*, Δ*ect*, Δ*bccT1* Δ*ect* mutant with (+) and without (-) supplementation of 1 mM ectoine with H) a representative microplate image of G). Growth was absent in the Δ*bccT1* Δ*ect* mutant at 636 mM NaCl in two out of four independent experiments. Data and error bars represent mean and standard deviations from independent biological replicates. Statistical analysis in C)-G) were performed with unpaired two-sided t-test with the WT strain as the control. (*p<0.05, **p< 0.01, ***p<0.001, ****p<0.0001)

To verify that the *bccT1* mutations in the AE strains cause a loss of transporter function, we constructed a *bccT1* knockout in the WT background and assessed growth on NaOAc and KOAc. Up to 182 mM NaOAc and 122 mM KOAc, the Δ*bccT1* and WT grew similarly (**Fig. 4C, D**). However, at 244 mM NaOAc and 182 mM KOAc, WT growth declined sharply, while the Δ*bccT1* mutant retained its growth rate. This suggests that Δ*bccT1* does enhance tolerance to higher concentrations of acetate salts, while the mutation does not improve acetate utilization per se.

Since BCCT transporters also facilitate bidirectional sodium ion flux, we hypothesized that a *bccT1* deletion might confer enhanced sodium tolerance independent of acetate metabolism. We therefore grew Δ*bccT1* and WT on 28 mM glucose with increasing NaCl, excluding acetate as a factor (**Fig. 4E**). The mutant exhibited stable growth up to 514 mM NaCl, above seawater salinity, whereas WT growth dropped abruptly at 453 mM (0.41[±[0.10[h[¹). Interestingly, WT cultures showed an apparent increase in growth rate at 514 mM NaCl (0.72[±[0.24[h[¹), but this coincided with pronounced flocculation and biofilm formation in the microplate wells **(Fig.**0**4H)**, suggesting that this rise in optical density may reflect physical aggregation rather than true growth recovery. At higher NaCl concentrations, no significant growth rate difference was observed between WT and Δ*bccT1*, likely due to the masking effects of flocculation on OD-based measurements. These results support a model in which *bccT1* deletion enhances osmotic resilience under salt stress—likely the key mechanism underlying its adaptive benefit during acetate salt challenge.

To further investigate the basis of this salt resilience, we considered the ecological niche of *V. natriegens* as a marine bacterium that routinely encounters external sources of compatible solutes. Under such conditions, uptake of osmolytes via BCCT1 would be advantageous, conserving the energy required for de novo biosynthesis. In contrast, our experimental setup lacked exogenous osmolytes, forcing cells to rely entirely on internally synthesized ones. We hypothesized that in the absence of BCCT1, cells may retain osmolytes more effectively, thereby enhancing salt tolerance under nutrient-limited conditions. Given that BCCT1 is the only BCCT transporter able to transport ectoine in *V. natriegens*^45^, we focused on ectoine as a model compound to test this hypothesis. Intracellular ectoine levels were quantified in WT and Δ*bccT1* strains grown at different NaCl concentrations. At 453 mM NaCl, the Δ*bccT1* strain contained 197% more ectoine per cell than WT, and 53% more at 636 mM (**Fig. 4F**). These findings suggest that loss of BCCT1 leads to enhanced ectoine retention under salt stress, supporting the mutant’s superior growth at NaCl concentrations that inhibit WT. Notably, at 636 mM NaCl, ectoine levels in the WT also increased, which likely contributes to its partial recovery at this concentration despite reduced growth at 453 mM (**Fig. 4E**).

To assess the role of ectoine biosynthesis, we deleted the *ectABC-aspC* operon and compared growth with and without 1 mM ectoine supplementation. The Δ*ect* mutant exhibited similar growth to WT at all tested NaCl concentrations (**Fig. 4G**), and both strains showed strong flocculation, unlike Δ*bccT1*, which remained planktonic (**Fig. 4H**). Surprisingly, the Δ*bccT1* Δ*ect* double mutant grew comparably to Δ*bccT1,* except at 636 mM, where growth failed in some replicates. This suggests that ectoine contributes primarily to survival under extreme salt stress, while the Δ*bccT1* phenotype at intermediate concentrations involves additional mechanisms. Supplementing ectoine rescued growth in Δ*ect* and WT, phenocopying Δ*bccT1* mutant growth rates, but it did not resolve the flocculation phenotype (**Fig. 4G, H**). Together, these findings indicate that although ectoine accumulation (**Fig. 4F**) enhances salt tolerance, it is not the sole contributor to the Δ*bccT1* phenotype. At intermediate NaCl levels, the dominant advantage likely stems from other mechanisms—such as reduced sodium influx or retention of alternative osmolytes—as evidenced by the similar performance of Δ*bccT1* and Δ*bccT1* Δ*ect* strains.

Taken together, these results suggest that the *bccT1* mutations in the adaptively evolved strains enhance tolerance to high acetate salt concentrations by promoting intracellular retention of ectoine and potentially other osmolytes, and/or by limiting sodium influx. Supporting the osmolyte retention model, we observed significant upregulation of the *ectABC-aspC* operon in AE2 and AE3 compared to WT (log[FC = 1.9 and 2.1, respectively; Table S4), indicating that these strains enhance growth under high NaOAc by both boosting ectoine biosynthesis and reducing its loss. Moreover, the superior growth of the Δ*bccT1* mutant at high KOAc concentrations (**Fig. 4D**)—despite potassium not being a known BCCT substrate—suggests a general mechanism of osmolyte retention rather than a specific ion effect. Altogether, our findings imply that while BCCT1 allows *V. natriegens* to exploit environmental osmolytes in its native habitat, its loss provides a selective advantage under minimal, ectoine-poor conditions by favoring internal accumulation of compatible solutes.

#### Characterization of luxU mutations

We observed LuxU frameshift mutations in two independent cell lines (**Fig. S6A**), with one strain (AE2) demonstrating the most highly accelerated growth rates on acetate, whereas the slower growing AE3 is lacking this mutation (**Fig. 2A, C, Table S5**). Since quorum sensing in *V. natriegens* has not been thoroughly investigated, we decided to explore how this cell density-based system impacts growth on acetate. Unlike its close relatives *V. parahaemolyticus* and *V. harveyii*, which have multiple enzymes for autoinducer synthesis, *V. natriegens* possesses only one annotated enzyme (LuxS) for autoinducer synthesis (**Fig. 5A**). However, similar to its relatives it encodes homologs of different sensor kinases (CqsS, LuxN, and LuxQ) for various autoinducer molecules and a two-component phosphorelay system (LuxU-LuxO), along with downstream global regulators AphA and HapR (LuxR homologue), which are known to control multiple genes involved in multicellularity, including the biofilm operon (*vps,* also known as *cps*), in *V. parahaemolyticus*^46,47^ (**Fig. 5A**).

**Figure 5.**
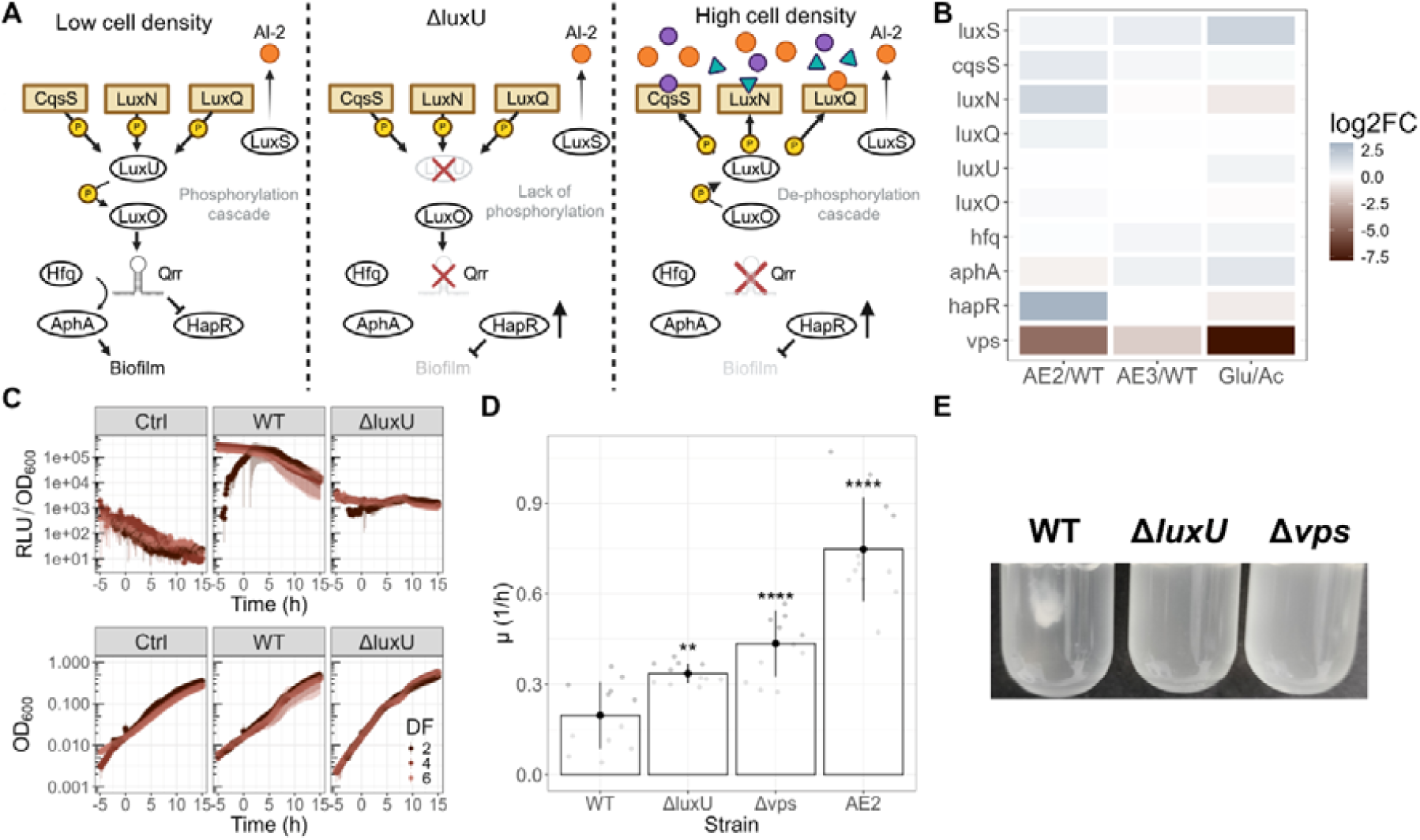
Characterization of the *luxU* deletion. A) Schematics of the suggested quorum sensing system in *V. natriegens*. B) Transcriptomics response of AE2, AE3 relative to WT during steady state growth on 61 mM NaOAc, as well as WT during growth on 28 mM glucose vs. 61 mM NaOAc (n = 3). C) Growth and relative luminescence unit (RLU/OD_600_) intensity of WT and Δ*luxU* strain transformed with a P*_vps_*-*luxCDABE* biosensor on 61 mM NaOAc with 3 different dilution factors (10^-DF^); n = 6). The control (Ctrl) strain harbors an empty plasmid. D) Growth rate of Δ*luxU* and Δ*vps* mutants in comparison to WT and AE2 at 61 mM NaOAc, as measured between OD_600_ = 0.025-0.1 (n = 12). E) Biofilm formation of different mutants statically grown on 6 mM glucose for 10 days at room temperature. Data and error bars represent mean and standard deviations from independent biological replicates. Statistical analysis in D) was done by a two-way unpaired t-test compared to the WT strain. (*p< 0.05, **p< 0.01, ***p<0.001, ****p<0.0001)

To understand how the common *luxU* frameshift mutation affects the cell, we conducted an *in silico* analysis, showing that in both evolved strains the functional domain H58, which is critical for its phosphorelay function^48^, was deleted (**Fig. S6**). An inability to phosphorylate LuxU is predicted to interrupt the Lux signaling cascade, leading to the upregulation of HapR (**Fig. 5A**, center). Consequently, we predict that the *luxU* frameshift mutation would mimic a high cell density state (**Fig. 5A**, right), regardless of their actual cell density, ultimately resulting in the downregulation of the *vps* biofilm operon.

To verify this, we first compared transcriptomic data between an evolved strain carrying the *luxU* frameshift mutation (AE2) and a strain without frameshift mutation (AE3) against the ancestral strain (WT) (**Fig. 5B, Table S6-S7**). As predicted, AE2/WT exhibited a stronger downregulation of the *vps* operon (log2FC = -4.6) compared to AE3/WT (log2FC = -1.3), suggesting that the *vps* operon might indeed be controlled by the LuxU-LuxO-AphA/LuxR pathway. However, this difference might be influenced by the higher NaOAc tolerance of AE3 relative to AE2 (**Fig. 2A**).

To further confirm that *vps* downregulation is due to *luxU* deletion, we constructed a Δ*luxU* mutant and transformed it along with the WT with a P*_vps_*-*luxCDABE* luminescence reporter, allowing us to monitor the activity of the P*_vps_* promoter in real time. Since the quorum sensing system depends on the autoinducer concentration in the media, stationary phase cultures were washed and resuspended in fresh MOPS2 media at three dilution factors (1:10^2^, 1:10^4^, and 1:10^6^), and incubated in a microplate reader (as the initial OD_600_ of the higher dilutions were below the detection limit, the time-axis of the three datasets was aligned at OD_600_ = 0.15, defined as t = 0 h). As expected, the WT displayed significantly higher luminescence signals than the Δ*luxU* mutant for every dilution factor (**Fig. 4C**), suggesting that the dilution in fresh media triggered the low cell density response in a *luxU*-dependent way. Interestingly, for the WT sample with the lowest dilution (1:10^2^), for which the initial OD was above the detection limit, luminescence increased sharply after dilution, reflecting the kinetics of P*_vps_* activation. In contrast, samples with higher dilution factors already display high luminescence signals at t = 0 h, consistent with the fact that these cultures had enough time to establish full P*_vps_* activity before reaching OD_600_ = 0.15. Regardless of the dilution factor, in the WT luminescence drops with increasing OD_600_, consistent with the deactivation of the quorum sensing system at high cell densities (**Fig. 5A**). This peak RLU/OD_600_ was also evident when glucose was used as a carbon source (**Fig. S6B**), though it decreased faster, likely because of the significantly increased growth rate on glucose^8^ (**Fig. 4E**) resulted in the cell spending less time in the low cell density state. This suggests that the *vps* operon was activated due to sudden depletion of autoinducers, triggering the low-cell density response through the Lux signaling cascade. Nevertheless, the RLU/OD_600_ signal of the Δ*luxU* mutant is still higher compared to the control strain harboring an empty plasmid, suggesting that there are additional regulatory mechanisms controlling the expression of the *vps* operon.

To further investigate the impact of *luxU* deletion on the growth rate, we grew Δ*luxU,* Δ*vps,* and WT strains in 61 mM NaOAc. Intriguingly, the growth rates of both the Δ*luxU* (0.34 ± 0.03 h^-1^) and the Δ*vps* (0.43 ± 0.11 h^-1^) strains were significantly higher than the WT (**Fig. 5C**), even though not as high as AE2 (0.75 ± 0.17 h^-1^). This confirms that reduced biofilm formation most likely contributes to the increase in growth rate in AE1 and AE2, both carrying a *luxU* frameshift mutation (cf. **Fig. 2B**). However, as noted before, this mutation is not the sole determinant of enhanced growth, as AE3—despite lacking the *luxU* mutation— also exhibits significantly improved acetate assimilation and biomass accumulation.

#### Putative role of additional mutations

All evolved strains carry nonsynonymous mutations in the transcriptional regulator *yheO* (**Fig. S2A,B**), although these affect different residues in each strain and their functional consequences remain unknown. In *E. coli*, YheO has been implicated in the regulation of the Pta/AckA pathway and acetate metabolism^49^. However, in our transcriptomic dataset, only *ackA2* is upregulated in both AE2 and AE3 while *pta1-ackA1* upregulation is restricted to AE2 (cf. **Fig. 2D**), suggesting that YheO may have a more limited regulon in *V. natriegens*, or that its effects depend on the specific mutations acquired during evolution. Additionally, AE2 and AE3 each harbor distinct mutations in the gene encoding a DUF3612 domain-containing protein, which is predicted to possess a DNA-binding function via its N-terminal Cro/C1-type helix-turn-helix motif (**Fig. S2C,D**). The divergence of these mutations across strains—alongside those in *yheO*—suggests that they may exert differential effects on global gene expression. Further transcriptomic analysis will be required to clarify their individual roles and contributions to the observed phenotypes. In addition to the distinct *bccT1* mutations found in all AE strains (see above), AE3 also carries a unique mutation in the intergenic region upstream of *proVWX*, an ABC transporter operon involved in osmoprotectant uptake. Collectively, these mutations may modulate stress response pathways or transport functions, thereby indirectly enhancing cellular fitness under acetate- or salt-stressed conditions.

### PHB production from acetate with adaptively evolved *V. natriegens*

#### PHB production in shake flasks

After demonstrating the enhanced growth rate, NaOAc tolerance, and reduced biofilm formation of the AE strains, we sought to determine whether these traits translated into improved bioproduction capabilities. To this end, we cloned the *phaBAPC* operon from *V. natriegens*–encoding three catalytic enzymes (PhaB, PhaA, and PhaC) that convert acetyl-CoA into PHB, along with the accessory phasin protein PhaP– into a high-copy plasmid under an anhydrotetracycline (ATc)-inducible promoter. This allowed us to temporally separate the growth and production phases by timed induction. We introduced the PHB-overproduction plasmid into two of the fastest-growing AE strains and the parental strain to compare poly-3-hydroxybutyrate (PHB) production.

To this end, we cultivated cells in MOPS2 minimal media containing 61 mM (3.60 g/L) acetate as the sole carbon source and induced PHB production at OD_600_ = 0.5. Intracellular PHB granules were visualized and quantified using BODIPY 493/503, a fluorescent dye that selectively stains hydrophobic lipid inclusions such as PHB. Fluorescence intensity per cell was measured via flow cytometry and correlated with PHB levels determined by HPLC (crotonic acid assay), confirming a linear relationship between BODIPY signal and PHB content per cell dry weight (**Fig. 6C**), consistent with prior studies^50^. Interestingly, the WT strain exhibited higher PHB fluorescence at 24 h—approximately double that of AE2— indicating a higher PHB content per cell (**Fig. 6D**). However, despite accumulating more PHB per cell, the WT consumed only 1.2 g/L of acetate over 48 hours, while AE strains nearly exhausted the acetate supply but produced less PHB overall. Specifically, AE2 and AE3 yielded PHB/acetate conversion ratios of 0.033 and 0.021 g/g, respectively, compared to 0.20 g/g in WT. Notably, AE strains produced approximately twice the biomass of WT (**Fig. 6E**), highlighting a shift in carbon allocation: while WT consumes little acetate and prioritizes PHB accumulation, AE strains can assimilate more acetate and favour biomass generation. This trade-off suggests that AE strains have evolved a more balanced strategy for carbon and energy usage, potentially offering improved process performance under nutrient-limited conditions. After 24 h, BODIPY fluorescence dropped markedly in all strains—by ∼50% in AE strains and ∼25% in the WT—indicating PHB degradation. AE strains showed a steady decline in fluorescence, with a subpopulation reverting to pre-induction levels (**Fig. 6F**), consistent with complete acetate depletion.

**Figure 6.**
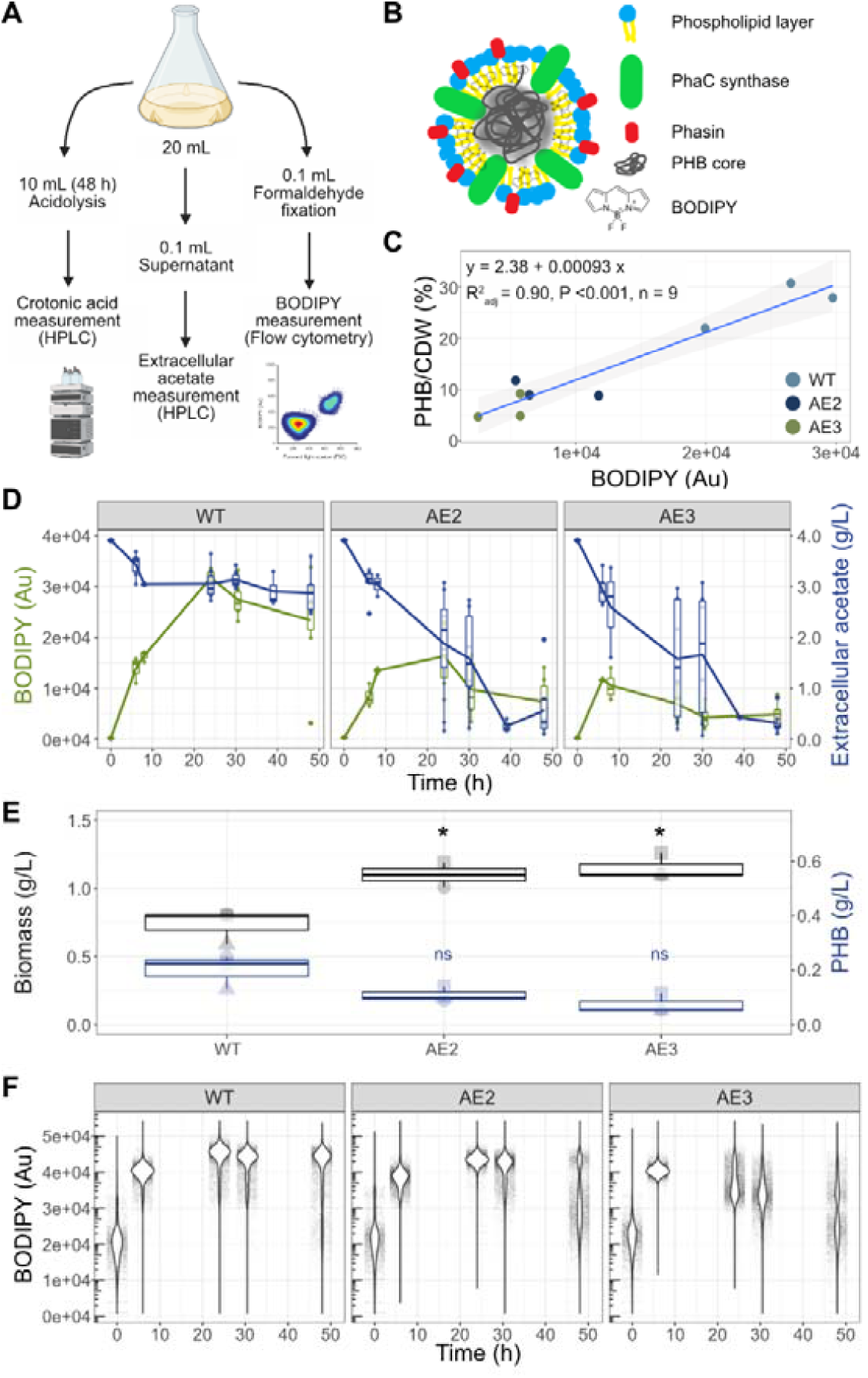
Shake flask experiment for PHB production from acetate using adaptively evolved *V. natriegens*. A) Diagram showcasing the sampling workflow, created using Biorender. B) Diagram illustrating how BODIPY 493/503 selectively stains intracellular PHB granules for fluorescence-based detection. C) Standard curve showing the relationship between average BODIPY fluorescence intensity (measured by flow cytometry) and PHB content (g PHB per g cell dry weight), quantified as crotonic acid by HPLC. Data were obtained after 48 h of cultivation at 30 °C in MOPS2 minimal medium containing 61 mM sodium acetate as the sole carbon source. PHB production was induced with 100 ng/mL anhydrotetracycline (ATc) at ODLJLJLJ = 0.5 in all three tested strains. D) Time-course analysis of intracellular PHB accumulation and substrate consumption. BODIPY fluorescence (box plots), measured by flow cytometry, reflects intracellular PHB content across timepoints. Extracellular acetate concentrations were determined by HPLC from culture supernatants. Data are shown for WT, AE2, and AE3 strains cultivated in MOPS2 minimal medium with 61 mM sodium acetate at 30 °C, with PHB production induced at ODLJLJLJ = 0.5 using 100 ng/mL ATc (n = 6). E) Final cell dry weight (biomass, black) and PHB content (blue), measured after 48 h of cultivation. Statistical significance was assessed using a two-way unpaired t-test relative to WT (*p < 0.05, **p < 0.01, ***p < 0.001, ****p < 0.0001; ns = not significant). **F)** Violin plots showing the distribution of BODIPY fluorescence (arbitrary units) within the population over time for each strain. Each point represents a single cell measured by flow cytometry (n > 10,000 per sample). A pronounced decrease in fluorescence—especially in AE strains after 24 h—indicates PHB degradation as acetate becomes limiting, with a subpopulation in AE2 and AE3 reverting to baseline PHB levels.

These observations can be rationalized by known differences in acetate assimilation pathways between the strains. The WT relies on a high-affinity, low-capacity pathway that functions efficiently at low substrate levels but limits overall flux (**Fig. 1E**). AE2, by contrast, predominantly employs a low-affinity, high-capacity pathway that is more energy-efficient and capable of supporting higher acetate uptake rates (**Fig. 2D–E**). This altered flux control plausibly explains the divergent PHB phenotypes. Assuming similar *phaBAPC* expression after induction, the WT may channel most of its limited acetate flux into PHB production, limiting energy and carbon availability for biomass formation. This would account for its lower growth rate, minimal acetate consumption, and higher intracellular PHB levels per cell. In contrast, AE2’s efficient acetate assimilation supports both biomass accumulation and PHB production, leading to continued acetate consumption following induction of *phaBAPC* operon expression. When external acetate eventually becomes limiting, AE2 appears to mobilize stored PHB to sustain growth under carbon starvation, resulting in a lower per-cell PHB content than the WT.

To investigate whether PHB degradation was enzymatically mediated, we examined potential depolymerases. A comparative genomics study identified two extracellular PHA oligomer hydrolases (encoded by *phaY* and *phaY2*) in *V. natriegens* NBRC15636 (= DSM 759), but no homologs of the intracellular PHA depolymerase gene *phaZ* were found across *Vibrio* species^51^. We constructed deletion mutants of *phaY* (RS21480), *phaY2* (RS22040), and a double mutant, and assessed PHB degradation in AE2 under carbon-starved conditions. However, degradation persisted in all cases (**Fig. S7C**), suggesting that yet-unidentified depolymerases are responsible for PHB turnover in *V. natriegens*.

#### Fed-batch fermentation

Given AE2’s enhanced capacity for acetate assimilation, particularly following *phaBAPC* operon induction, we hypothesized that the PHB production titer, rate, and yield could be improved by implementing a fed-batch process. Namely, by matching substrate supply to the strain’s increased demand, we aimed to promote PHB accumulation while preventing its degradation due to carbon depletion. An overview of the fed-batch fermentation system is provided in **Fig. 7A**. Following initial acetate depletion in the batch phase, we implemented substrate feeding to attain high cell densities, then induced PHB production via ATc addition. To prevent growth inhibition from excess residual acetate accumulation, we tested reactive feeding strategies, initially using a DO-stat system, where rapid increases in dissolved oxygen (DO signal), indicative of acetate depletion, triggered substrate feeding^52–54^. While this maintained low residual acetate levels (≤ 1 g/l) during growth, it failed to prevent carbon depletion post-induction. With a large fraction of the carbon flux being directed to PHB biosynthesis instead of respiration, the DO concentration no longer responds proportionally to substrate depletion. This leads to a decrease in feeding rate, triggering PHB consumption in the latter stages of fermentation (**Fig. S8**).

**Figure 7.**
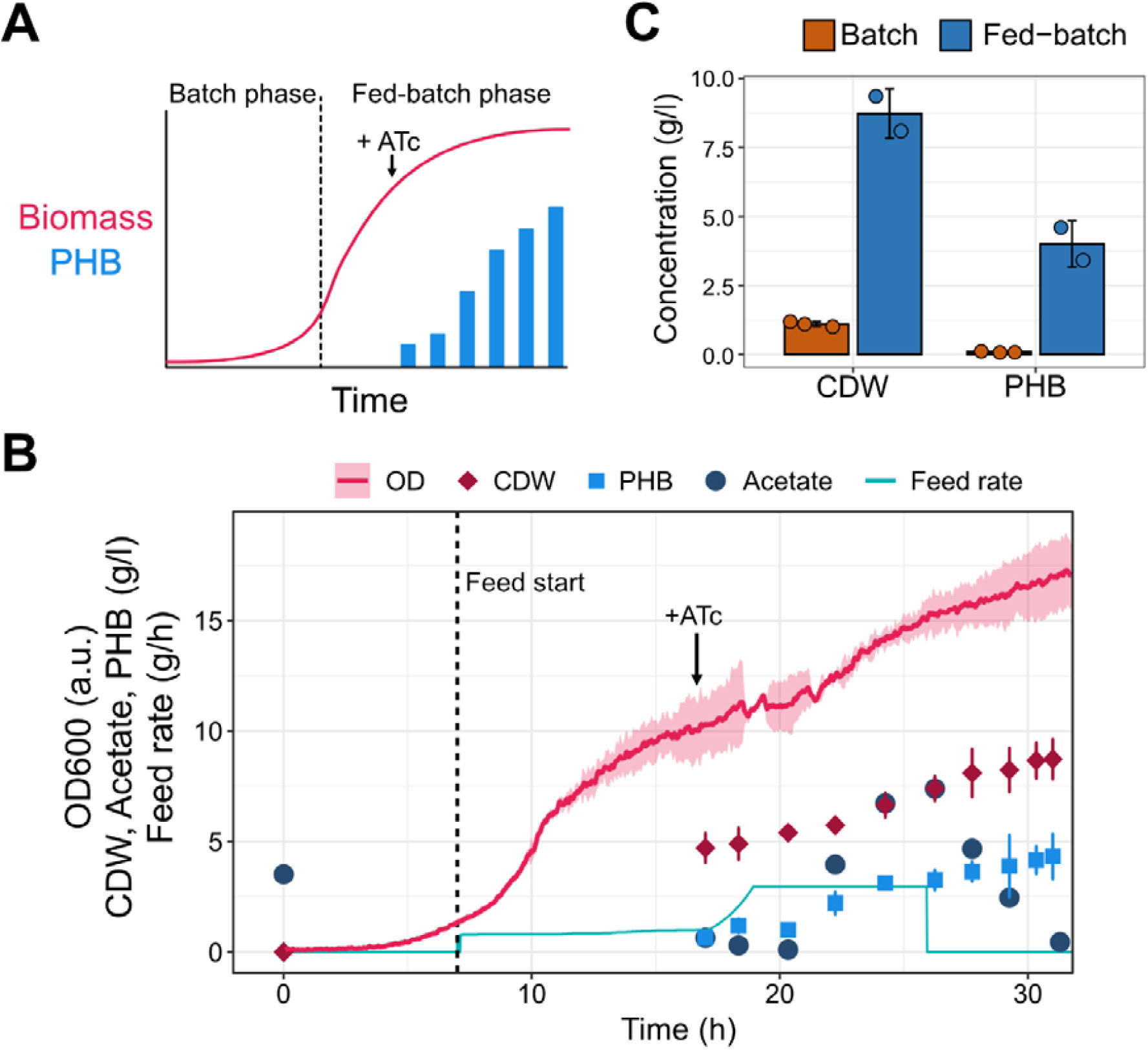
Improving PHB production through fed-batch fermentation. A) Schematic diagram of the fed-batch fermentation system. Feeding starts after the initial amount of substrate is depleted in the batch phase (dashed line). PHB production is triggered in the latter stages of fermentation by addition of the inducer ATc (black arrow). B) Fed-batch PHB production. The start of substrate feeding and induction time are indicated as in A). The mean and standard deviation of the optical density (OD, measured online) are shown as a solid pink line and pink shading, respectively. At each sampling timepoint, the mean CDW (red diamonds) and PHB (blue squares) titre is shown, with error bars indicating the standard deviation. For simplicity, representative feed rate and residual acetate traces are shown (solid green line and dark teal circles, respectively). All other data correspond to duplicate experiments. C) Comparison of maximum CDW and PHB titres attained for strain AE2 in batch (shake flask) and fed-batch fermentations. Bars represent the mean and standard deviation of triplicate (batch) or duplicate (fed-batch) experiments, with experimental data points shown as individual dots.

To circumvent this, we instead adjusted feeding dynamically as a function of the pH control. As acetate was consumed, rising pH triggered acid addition, which we used as a proxy for acetate demand. Unlike pH-stat fermentation—where the carbon source itself controls the culture pH^55^—this approach maintained a small residual acetate pool, promoting PHB synthesis while preventing degradation. However, even this strategy could not fully meet AE2’s demand for carbon, which increased exponentially following induction (**Fig. S9**). To resolve this, we developed a hybrid feeding strategy: before induction, the feed rate was proportional to acid addition; after induction, it increased exponentially up to a set maximum (**Fig. S10**). This ensured excess carbon supply only during PHB production, where the benefit of a high residual substrate concentration outweighs the potential growth inhibition from overfeeding.

With this strategy, PHB production reached 4.01 ± 0.84 g/L, corresponding to 45.66 ± 4.97 % of cell dry weight (CDW) (**Fig. 7B, C** and **Table 1**). Remarkably, AE2 achieved a volumetric productivity of 0.13 ± 0.03 g/L/h over the full fermentation, or 0.27 ± 0.06 g/L/h in the production phase (i.e. the time following ATc addition). These results demonstrate that dynamic carbon supply, tailored to AE2’s metabolic needs, was both necessary and sufficient for enhanced biopolymer production.

**Table 1.** Fed-batch production parameters. All values are reported as the mean ± one standard deviation of duplicate experiments.

|  |  |
| --- | --- |
| PHB (g/L) | $4.01 \pm 0.84$ |
| CDW (g/L) | $8.73 \pm 0.89$ |
| PHB (%CDW) | $45.66 \pm 4.97$ |
| g PHB / g acetate (%) | $12.98 \pm 1.52$ |
| mol PHB / mol acetate (%) | $7.36 \pm 0.86$ |
| Volumetric productivity (g/L/h) | $0.13 \pm 0.03$ |

## Discussion

This study establishes *Vibrio natriegens* as a viable platform for sustainable, cost-effective bioproduction using acetate as a feedstock. Through ALE and optimized fed-batch fermentation, strain AE2 achieved a remarkable PHB productivity of 0.27 ± 0.06 g/L/h during a 15 h production phase, reaching a final titre of 4.01 ± 0.84 g/L (45.66 ± 4.97% of CDW). This productivity compares favourably to established industrial microorganisms, including *Halomonas* sp. (0.04 g/L/h, 112 h)^56^*, Cupriviadus necator* DSM 545 (0.17 g/L/h, 29 h)^57^, *Halomonas bluephagenesis* B71 (0.33 g/L/h, 150 h)^58^, and *Salinivibrio* spp TGB19 (0.37 g/L/h, 144 h)^59^, despite our deliberate omission of complex nitrogen and carbon supplements.

The omission of complex nutrient sources, such as yeast extract or peptone, aimed to minimize production costs, which remain a major barrier to commercializing PHB production^60,61^. Nevertheless, previous research has indicated substantial productivity enhancements with these supplements; for example, productivity in *Salinivibrio* spp. increased 7.4-fold with tryptone supplementation^59^, *Yarrowia lipolytica* productivity improved 1.7-fold with yeast extract and peptone^62^, and *E. coli* JM109 achieved a remarkable 64-fold increase with yeast extract^63^. Thus, although our minimalist strategy demonstrates economic benefits, carefully balancing production expenses against productivity improvements through rigorous technoeconomic analyses will be critical for industrial adoption.

Adaptive laboratory evolution significantly improved the acetate assimilation capability and tolerance of *V. natriegens*, especially at higher acetate concentrations, as previously observed^34^. Beneficial mutations contributing to this phenotype were identified in genes encoding the betaine-choline-carnitine transporter *bccT1*, the transcriptional regulator *yheO*, and the quorum sensing regulator *luxU*.

A previous study in *E. coli* also noted mutations in a compatible solute transporter *proV* when performing ALE towards various commodity chemicals with high osmolarity^64^. These *proV*-deficient strains were a result of increased salinity due to the presence of salt counterions. As recent studies suggest that the optimal sodium concentration for *V. natriegens* growth ranges around 7.5-17.5 g/L^15,65^ (lower than the salinity in MOPS2 media), it is not unexpected to observe a common mutation enhancing sodium tolerance during our ALE experiments. We further observed that this mutation conferred tolerance towards acetate. This is in line with previous observations in *V. cholerae* and *V. parahaemolyticus,* where preadaptation to salt has conferred tolerance towards acidic stress^66^, but the underlying mechanisms are yet to be identified.

Intriguingly, we identified a frameshift mutation in the quorum sensing gene *luxU* in strain AE2. To rationalize the impact of this mutation, we developed a first quorum sensing model in *V. natriegens* based homology to regulators in well-studied *Vibrio* species^48,67–70^ (**Fig. 5A**). The model suggests that the exopolysaccharide biosynthetic operon (*vps*) is normally upregulated at low cell densities (**Fig. 5A**). Consequently, a *luxU* deletion is predicted to maintain cells in a continuous "high cell density state"^46–48,67,71^, suppressing *vps* expression, consistent with our experimental data (**Fig. 5B, C**). However, this contrasts with findings from a previous study reporting an increase in EPS production during bioproduction at higher cell densities^72^. This discrepancy is likely methodological, as the prior report used a washing step before the production phase, likely removing autoinducers and triggering a low-density response^72^. In our context, continual suppression of *vps* expression prevents unnecessary energy expenditure on exopolysaccharide production, especially beneficial when acetate is the sole, energetically challenging carbon source. Consequently, reduced EPS formation likely contributed significantly to improved growth efficiency and elevated growth rates observed in AE2 (**Fig. 5D**).

The *vps* operon was one of the most upregulated gene clusters in the parental *V. natriegens* strain when grown on acetate compared to glucose (**Table S6**). This upregulation may represent an adaptive strategy to counteract acetic acid-induced stress, similar to the protective function of capsular polysaccharides observed in *Acetobacter tropicalis*^73^. Conversely, in AE strains, particularly AE2, the *vps* operon was notably downregulated. This downregulation likely results from enhanced acetate and sodium tolerance conferred by mutations such as *bccT1*, rendering EPS-based protective mechanisms less critical (**Fig. 3**). Importantly, previous research demonstrated that reduced *vps* expression decreased culture viscosity 5-6 fold and improved product yields by 27%, both advantageous traits for bioprocesses^72^. Thus, the downregulation of EPS production not only reduces metabolic burdens but also positively impacts biotechnological performance.

Furthermore, quorum sensing has been shown to regulate the Pta/AckA pathway in *Vibrio harveyi*^70^, which may explain the upregulation of the *pta1-ackA1* operon observed in AE2 compared to AE3 (**Fig. 2D**), as the latter does not carry the *luxU* frameshift mutation. Although *yheO* has been implicated in controlling the Pta/AckA pathway in *E. coli*^49^, our data suggests that in *V. natriegens* its influence may be limited to upregulating *ackA2*. During acetate metabolism, AckA catalyzes the conversion of acetate to acetyl-phosphate, a key intermediate known to influence various two-component regulatory systems across different bacterial species^40^. The increased acetyl-phosphate levels in AE strains could thus drive their enhanced acetate utilization and overall improved growth performance. However, due to the differing mutation sites between strains (**Fig. S2**), further transcriptomic investigation is necessary to elucidate the precise regulatory mechanisms at play.

These results collectively suggest that during growth on poor carbon sources like acetate, it is advantageous for cells to minimize investment in biofilm formation and other energetically costly processes—a pattern commonly observed in ALE experiments. This strategy aligns with established metabolic engineering approaches that redirect flux away from non-essential functions, as demonstrated in *Halomonas* spp.^74,75^ and *V. natriegens*^72^. Our comparative analysis of AE2 and AE3 further illustrates how distinct genetic changes can converge on similar phenotypic outcomes. AE2 carries a frameshift mutation in *luxU*, leading to strong downregulation of the *vps* operon and reduced biofilm formation. Beyond biofilm regulation, broader effects on growth may also arise from changes in quorum sensing circuitry, particularly given that HapR—a LuxR homologue—regulates hundreds of genes in *V. parahaemolyticus*^70^, suggesting that *luxU* inactivation could exert pleiotropic effects on cellular physiology. Both AE2 and AE3 exhibit strong upregulation of *ackA2*, potentially driven by the shared but distinct mutations in *yheO*, a transcriptional regulator known to influence acetate metabolism in *E. coli*. Notably, only AE2 shows additional upregulation of the *pta1-ackA1* operon, which may be linked to *luxU* inactivation and its downstream regulatory impact. This more extensive transcriptional response could explain AE2’s enhanced acetate flux and growth phenotype. In contrast, AE3—lacking the *luxU* mutation— harbors a unique combination of other mutations, including a mutation in the intergenic region upstream of *proVWX* and distinct strain-specific changes in both the DUF3612-domain protein and *bccT1*, all of which may collectively modulate stress response, transport, or global regulatory dynamics. These findings underscore the regulatory flexibility of *V. natriegens* and highlight multiple viable evolutionary trajectories toward optimizing growth and bioproduction from acetate.

In conclusion, these findings demonstrate the potential of *V. natriegens* as a promising chassis for sustainable PHB production from acetate. The combination of enhanced growth rate, improved acetate and salt tolerance, upregulation of the Pta/AckA pathway, and reduced EPS production makes the AE strains, particularly AE2, promising candidates for scalable acetate-based bioproduction. While further optimization can maximize PHB yields, the progress achieved through adaptive evolution, genetic engineering, and bioprocess engineering demonstrates a strong foundation for future advancements. By further refining key metabolic pathways, such as deleting any (yet undiscovered) depolymerase genes, and improving bioprocess strategies, *V. natriegens* can be developed into an efficient and scalable system for marine-based biopolymer production, offering significant promise for industrial applications.

## Methods

### Bacterial strain and growth conditions

All bacterial strains used in this study are listed in **Table S8**. For transformation and genome engineering protocols, *V. natriegens* DSM 759 ΔVNP1+2 Δ*dns* cells were grown on LBv2 agar solid media, as previously described^20^. For plate reader and shake flask experiments, *V. natriegens* was streaked, inoculated, and diluted in modified “MOPS2” minimal media, which is based on Neidhardt medium^76^ with minor adjustments, including the supplementation of 2% (w/v) (342.2 mM) NaCl to simulate ocean salinity, mimicking a previously published strategy^32^. This medium was chosen due to its independence from a phosphate-based buffering system, since phosphorus levels may affect PHB production^77,78^. The 1X MOPS2 media (pH 7.2) contained the following components: MOPS (42 mM), Tricine (4 mM), K_2_SO_4_ (0.276 mM), CaCl_2_ (0.5 μM), MgCl_2_ (0.525 mM), NaCl (total 392.2 mM), FeSO_4_ (10 μM), K_2_HPO_4_ (1.32 mM), NH_4_CL (9.5 mM), and the micronutrients, (NH_4_)_6_Mo7O_24_ (3 nM), H_3_BO_3_ (0.2 μM), CoCl_2_ (1.5 nM), CuSO_4_ (0.48 nM), MnCl_2_ (4 nM), and ZnSo_4_ (0.48 nM), as well as the tested carbon source (61 mM sodium acetate (NaOAc) for standard plate reader experiments and overnight culture). All components were filter-sterilized using a 0.22 μm filter. For solid MOPS2 medium, agar was added into ddH_2_O with a final concentration of 3% (2X agar solution) and was microwaved in a sterile container. The 2X agar solution was left at room temperature until reaching approximately 70 °C prior to addition of the MOPS2 components. The NaOAc stock solution (20% w/v) was neutralized with HCl to pH 7. Kanamycin was added at a concentration of 200 ng/mL to plasmid-based experiments with *V. natriegens*.

### Generation of *V. natriegens* mutants

Genome modification was carried out following the NT-CRISPR workflow described by Stukenberg et al. (2022)^21^, with minor modifications. In brief, transfer DNA (tDNA) assembly was performed using either fusion PCR or Golden Gate assembly (see oligonucleotides in **Table S9**). *V. natriegens* cells, transformed with the NT-CRISPR plasmid containing the desired gRNA (see oligonucleotides in **Table S10**), were grown overnight in LBv2 medium supplemented with 100 µM IPTG and 4 µg/mL chloramphenicol at 30°C (OD_600_ ∼14-16). The overnight culture was then diluted 1:100 into 350 µL of sea-salt medium (28 g/L) containing approximately 100 ng of tDNA and incubated at 30 °C under static conditions for 5 hours. Following this incubation, 1 mL of LBv2 medium containing 200 ng/mL ATc was added to the culture, which was then shaken at 400 rpm for 1 hour at 30 °C. After this induction step, the culture was diluted 1:100 in LBv2, and 20 µL was streaked onto a square agar plate prior to an overnight incubation step at 30 °C. The colonies were verified through colony PCR through the method previously described^21^ (see oligonucleotides in **Table S11**).

### Plasmid construction

Plasmids used in this study are listed in **Table S12** and **S13**. The plasmids were constructed using the Marburg collection, a Golden-gate based toolbox optimized for *V. natriegens*^20^. The coding sequences for the *phaBAPC* operon and the P*_vps_* promoter were amplified from genomic DNA using Q5 polymerase with primers listed in **Table S12**, and purified using the EZNA Cycle Pure Kit (Omega BioTek) according to the manufacturer’s instructions. The purified amplicon was then cloned into the pMC_V_01_Part_Entry_sfGFP plasmid^20^ and transformed into *Escherichia coli* DH5α. Successful assembly was confirmed by colony PCR, and plasmid isolation was performed using the EZNA Plasmid DNA Mini Kit (Omega BioTek), following the manufacturer’s protocol. For overnight cultures, chloramphenicol and kanamycin were added at a final concentration of 25 ng/mL and 50 ng/mL, respectively.

### Competent cells and transformation

Competent *V. natriegens* cells and transformation were performed as described in Stukenberg et al. (2021)^20^. Briefly, an overnight *V. natriegens* culture was inoculated in 125 mL of preheated LBv2 medium at 37 °C to an OD_600_ of 0.01 in a baffled Erlenmeyer flask. The culture was then incubated at 200 rpm and 37 °C until reaching an OD_600_ of 0.5-0.8 prior to being transferred to precooled 50 mL falcon tubes and placed on ice for 10 minutes. Cells were then centrifuged at 3.000x g for 10 minutes at 4 °C. The pellets were resuspended in a 40 mL cold TB buffer for each 125 mL of bacterial culture. Following another incubation on ice and centrifugation, pellets were resuspended in 5 mL cold TB buffer, consolidated in a single tube, and 350 µL of DMSO was added. The suspension was incubated on ice for 10 minutes, after which 50 µL aliquots were snap-frozen in liquid nitrogen and stored at −80 °C.

For transformation, DNA was added to aliquots of competent *V. natriegens* cells, which were incubated on ice for 30 minutes. Heat shock was performed at 42°C for 45 seconds, followed by a 10-minute incubation on ice. The cells were then recovered in 1 mL warm LBv2 medium, shaking at 37 °C for 1 hour. After recovery, cells were centrifuged at 3.000x g for 1 minute, the supernatant was removed, and the pellet was resuspended in approximately 50 µL of the residual medium. The full volume was plated on LBv2 plates containing the appropriate antibiotic and incubated overnight at 37 °C.

### Predicting acetate flux with genome scale model

To investigate acetate metabolism in silico, we used the *V. natriegens* genome scale metabolic model (GSMM) devised by Coppens et al. (2023)^15^, iLC85815. The GSMM was obtained from the original publication in SBML format (https://github.com/LucasCoppens/Modelling_Vibrio_natriegens). Three manual modifications were performed. First, the cofactor for the reaction catalyzed by acetoacetyl-CoA reductase was changed to NADPH (from NADH). Second, a boundary (sink) reaction for PHB was added, to enable the simulation of PHB production. Third, the reaction catalyzed by acetyl-CoA synthetase (ACS, E.C. 6.2.1.1), which was found to be absent from the model, was added. This modified version of the model is available at https://github.com/FritzLabUWA/Vnat_GSMM. Using the modified version of the model, flux balance analysis (FBA) was conducted to predict growth rates and carbon flux distributions. Biomass formation was set as the model objective. The acetate uptake rate was constrained to a default value of 10 mmol/g_CDW_/h, enabling the quantification of optimal flux distributions in relative terms only (i.e., quantification of the flux through each model reaction as a percentage of the substrate uptake rate, or calculation of molar ratios). Uptake rates for all other carbon substrates were set to 0 mmol/g_CDW_/h. To simulate growth-only conditions, PHB accumulation was omitted from the FBA simulation by setting the upper and lower bounds of the PHB sink reaction to 0 mmol/g_CDW_/h. All GSMM modifications and analyses were conducted using the COBRApy package in Python. The d3flux package (https://pstjohn.github.io/d3flux/) was used to automatically generate D3.js-based metabolic visualizations of the COBRApy output.

### Adaptive laboratory evolution

*V. natriegens* DSMz 759 ΔVNP1+2 Δ*dns* was revived from cryostock, and three independent colonies were grown in MOPS2 medium supplemented with 20 g/L NaCl and 122 mM NaOAc in a 96-deep-well plate (2.2 mL) at 37 °C, using a shaking incubator set at 220 rpm. The cultures were covered with a Breathe-Easy® sealing membrane to enhance oxygenation. Every 24 hours, the cultures were diluted 1:1,000 into fresh medium, for a total of 100 days, equating to approximately 100 × *log*_2_(1,000) = 997 generations, allowing beneficial Every 24 hours, the cultures were diluted 1:1,000 into fresh medium, for a total of 100 days, mutations to accumulate and outcompete less advantageous variants. After 100 days, each of the three cultures was streaked onto 122 mM NaOAc MOPS2 agar plates, and two colonies from each culture were selected based on the largest colony size. These colonies were stored at -80 °C in 25% glycerol for subsequent experiments.

For each of the adaptively evolved strains, two colonies were chosen for genomic analysis. The ancestral strain was re-sequenced in duplicate to establish a reference genome for comparison. Genomic DNA (gDNA) was extracted from stationary-phase cells using the chloroform: isoamyl alcohol method. Briefly, cells were resuspended and lysed in PL2 buffer (1% SDS, 10 mM Tris pH 7, 10 mM EDTA, 0.2 mg/mL RNase) and incubated at 65 °C for 30 minutes at 400 rpm in a thermoshaker, simultaneously deactivating DNase and degrading RNA. An equal volume of PL3 (2.5 M KOAc) and chloroform: isoamyl alcohol (24:1) was added to the solution, which was then mixed and centrifuged at maximum speed for 10 minutes. The supernatant was transferred to a new tube containing 0.7 volumes of isopropanol and incubated at room temperature for 10 minutes. After centrifugation, the pellet was washed with 70% ethanol, air-dried, and resuspended in the TE buffer. The quality of the gDNA was verified using 0.8% TAE agarose gel electrophoresis before proceeding to next-generation sequencing.

Genome sequencing was performed using the DNBseq method^79^. The resulting FASTQ files were imported into Geneious Prime 2021 for analysis. Reads were mapped to the reference genome using in-built “Geneious” mapper medium-to-low sensitivity settings. Mutations were generated via the in-built “Find Variations/SNPs” function with a frequency threshold of >0.5. Protein models were generated with AlphaFold3^80^ and visualized in ChimeraX^81^.

### Plate reader experiments

The bacterial growth kinetics were measured as OD_600_ using a Biotek Synergy HTX microplate reader. To compare the growth rate of the AE strains to the ancestor WT strain, it was essential to rule out transcriptional adaptation as a confounding factor, considering that the AE strains had been growing on NaOAc for 997 generations. Thus, all cultures were preadapted to NaOAc, with AE strains grown for 16 hours and WT for 24 hours in 122 mM NaOAc MOPS2 media. This pre-adaptation phase allowed the WT strain to adjust transcriptionally to NaOAc, ensuring any observed growth rate differences were not due to initial transcriptional changes. To compare the Δ*acsA*, Δ*ackA*, and Δ*pta* mutants to its parental strain, the cells were pre-adapted in LBv2, as these mutants were unable to grow consistently at 61 mM NaOAc. To compare the growth rate of Δ*bccT1*, Δ*vps*, and Δ*luxU* mutants to the WT, the cultures were preadapted at 61 mM NaOAc for 24 hours. Unless indicated, stationary phase cultures (OD_600_ ∼3.5 on 61 mM NaOAc MOPS2) were diluted 10^4^-fold into a 100 µL of fresh medium in a transparent flat-bottom 96 well plate for OD_600_ measurement. This high dilution factor allows the cells to reach enough doublings required for steady state growth^82,83^, during which the growth rate was measured (described below). Spaces between wells, the outer wells, and unused wells were filled with ddH2O water to minimize evaporation. Experiments involving luminescence were performed in the same manner but using a black flat-bottom 96-well plate instead.

The characterization of the P*_vps_* promoter was performed by growing the respective cells with plasmids in liquid LBv2 media until reaching an OD_600_ ∼ 1.0 prior to isolation of the cell. The cells were pelleted at 10.000x g for 1 minute and were subject to washing and resuspension in MOPS2 medium without any carbon source, and serial dilution (10^2^, 10^4^, and 10^6^-fold) into the plates within two minutes after resuspension. For the glucose-based experiments, cultures were preadapted on 28 mM glucose to an OD_600_ ∼2 prior to 10^4^ dilution into the corresponding medium.

The plate was subsequently incubated at 37 °C with orbital shaking at 425 cpm in the plate reader. When applicable, luminescence was measured from the top (read height = 4.50 mm) with a gain of 135 and an integration time of 1 s. The raw data was blank-subtracted with the average luminescence reading of three blank wells containing uncultured media. The blank-subtracted data were computationally shifted to start at the same OD_600_ = 0.025 for acetate-based experiments and OD_600_ = 0.015 for glucose-based experiments. Unless indicated, the growth rate was determined by measuring at the mean slope of a linear regression model of each growth curve with the lm() function in R during steady state growth between plate reader OD_600_ of 0.025-0.1.

### Shake flask experiments

Cultures for shake flask experiments were plated out in solid MOPS2 medium with 61 mM NaOAc and inoculated in liquid MOPS2 media with 61 mM NaOAc for 24 hours (WT) or 16 hours (AE) at 37 °C with 220 rpm shaking. Stationary phase cultures (4 mL of OD ∼3.5) were pelleted, washed and resuspended in 20 mL volume of fresh media supplemented with 100 ng/mL anhydrotetracycline (ATc) in 100 mL triple-baffled flasks. The induced cultures were grown at 37 °C with 220 rpm shaking for 48 hours in baffled flasks. Time point samples were taken regularly for acetate quantification (200 μL, see below) and flow cytometry (200 μL, see below). At 48 hours, the final PHB titer was measured as crotonic acid using high pressure liquid chromatography (HPLC).

### Intracellular ectoine measurement

Cells were cultured in MOPS2 medium containing 28 mM glucose and grown to stationary phase (OD_600_ ∼2), followed by a 10^3^-fold dilution into fresh MOPS2 medium with 28 mM glucose and varying NaCl concentrations (392, 453, and 636 mM). Once the cells reached an OD_600_ of approximately 0.5, 1 mL of culture were harvested by centrifugation at 5000x g for 1 minute. The supernatant was carefully removed, and the cell pellet was resuspended in 0.5 mL ddH2O. The resuspended cells were stored at -20°C until further processing. For cell lysis, a VCX400 sonicator (Sonics and Materials) equipped with a 3 mm tapered microtip was used at 10% amplitude, employing 10-second on and 10-second off cycles for 3 minutes. Cell debris was removed by centrifugation at 21,000x g for 2 minutes, and the resulting supernatant was used to quantify intracellular ectoine levels using the protocol described below.

### Analytical methods

Samples for extracellular acetate measurement were collected by centrifuging culture samples at 10,000x g for 1 minute and diluting the supernatant 1:10 in ddH_2_O. To quantify the volumetric PHB titer, samples underwent acidolysis and were measured as crotonic acid. Briefly, 10 mL of sample was collected by centrifuging the samples at 7.197x g for 5 minutes. The pellet was rinsed with ddH_2_O and then transferred into a pre-weighed glass tube. The cells were dried overnight at 60 °C and the difference in weight was measured to obtain the cell dry weight (CDW). Acidolysis was executed by adding 1 mL of 98% sulfuric acid into the pellet with a 1-hour incubation in boiling water. The reaction was cooled down to room temperature prior to a dilution with 4x volume ddH_2_O on ice. The samples were further diluted by another 10-fold dilution in ddH_2_O.

Acetate and crotonic acid concentrations were quantified using an Agilent 1100 High Pressure Liquid Chromatography (HPLC) system equipped with a photodiode array detector (DAD) following the protocol from Dalia et al. (2017)^84^ with minor modifications. Briefly, separation was performed on a Phenomenex REZEX ROA-Organic Acid H^+^ (8%) 300 x 7.8 mm column using 3 mM sulfuric acid as the mobile phase. The flow rate was 0.5 mL/minute, injection volume was 20 µL, and the column temperature was maintained at 60 °C. For ectoine measurement, separation was performed on a Reprosil-XR 5 µm 250 x 4.6 mm C18 column using 0.1% trifluoroacetic acid as the mobile phase. The flow rate was 1 mL/minute, injection volume was 20 µL, and the column temperature was maintained at 30 °C. Peak areas recorded with a diode array detector at 210 nm were analysed using ChromatographR^85^ and quantified against a calibration curve generated from the respective standards prepared in ddH_2_O. The PHB-to-crotonic acid conversion efficiency was independently validated using a commercial PHB standard (Sigma Aldrich, 915092), yielding a conversion rate of 91% (**Table S14**). Final PHB titers were calculated from crotonic acid concentrations using this empirically determined conversion factor.

### Flow cytometry

To prepare samples for flow cytometry, bacteria cultures were harvested at different time points from liquid culture and were fixed using formaldehyde. Briefly, cold formaldehyde was added to the liquid culture to a final concentration of 4% and was incubated at 4 °C for 1 hour. Cells were subsequently washed twice, which includes a centrifugation step at 10.000x g for 1 minute, removal of the supernatant, and resuspension in 0.5x volume of fresh medium. Cells were subsequently resuspended in the same media and stored at 4 °C until analysis, which was typically done within 7 days. The fixed samples were diluted 1:200 to a concentration of ∼10^4^-10^5^ cfu/mL in 0.5x PBS with 5 mM EDTA prior to staining. To visualize intracellular PHB^50^ and DNA, cells were double-stained with BODIPY 493/503 (Sigma-Aldrich)^87^ and Syto62 (Thermofisher) at final concentrations of 2 μM and 1 μM, respectively. Staining was performed at 37° C for 15 minutes.

Data acquisition was performed with FACS Canto (BD Biosciences) with 1x PBS as shear fluid with 10,000 events as a limit. The BODIPY and Syto62 signals were quantified at 488 nm (excitation) / 530 ± 30 nm (emission) and 633 nm (excitation) / 660 ± 20 nm (emission) wavelengths, respectively. The forward scatter, side scatter, BODIPY and Syto62 voltage were set to 450, 430, 420, 603, respectively. The resulting output was analyzed and visualized with the RStudio packages, FlowCore^88^ and openCyto^89^. To distinguish the bacteria from debris, the data were gated to events positively stained by Syto62.

### Transcriptomic analysis

Single *V. natriegens* colonies were grown at 37° C at 220 rpm in MOPS2 medium supplemented with 5 g/L glucose (28 mM, WT) or 5 g/L NaOAc (61 mM, WT, AE2, AE3) starting OD_600_ of 2×10^-4^ in a 1 L baffled flask and were grown at 37 °C with 180 rpm until stationary phase. The cultures were diluted into 100 mL of the tested medium with a shaking. 50 mL of cells in steady state exponential growth (OD_600_ ∼ 0.2-0.25) were isolated via centrifugation for 3 minutes at 7,200x g. The supernatant was removed, and the pellets were flash frozen with liquid N_2_. The samples were kept at -80 °C until RNA extraction. The RNA extraction was performed using a Rapid RNA Isolation Kit (BioBasic Inc) and the residual gDNA was trimmed using a Turbo Free Kit (ThermoFisher Scientifics) as per manufacturer’s instructions. The quality of the isolated RNA was confirmed with a TapeStation (Agilent) and the quantity was checked with Qubit (Thermo Fisher Scientific). Library preparation was performed using the NEBNext rRNA depletion (Bacteria) and the TruSeq Stranded Total RNA Library Prep Gold Kit, following the manufacturer’s protocols. The total RNA library was subsequently sequenced with an Illumina sequencer Novaseq6000.

The resulting FASTQ files were analyzed using Geneious Prime 2021. The adapter sequences in the raw paired-read data were trimmed using BBduk with standard sensitivity. The trimmed data was subsequently mapped to the genome of *V. natriegens* DSM 759 ΔVNP1+2 Δ*dns* using BBmap. Individually mapped reads were counted through the “Calculate expression levels” function in Geneious Prime 2021 with “Count as partial matches” as a parameter. The resulting reads were compared between different conditions with DEseq2 using parametric settings to obtain the differentially expressed genes. The resulting genes with an adjusted P-value (padJ) <0.05 were mapped into KEGG using the R package, clusterProfiler^90,91^, with the gseKEGG() function.

### Fed-batch fermentations

Fed-batch fermentations for PHB production with strain AE2 were carried out in 1.5 L CloudReady parallel bioreactors (TJX Bioengineering). Cells were first streaked out on MOPS2 agar solid medium supplemented with 61mM NaOAc. Following overnight incubation (14-16 h) at 37 °C, individual colonies were picked and used to inoculate precultures in 100 mL baffled conical flasks (20 mL culture volume, MOPS2 supplemented with 61 mM NaOAc). Precultures were incubated overnight at 37 °C with 220 rpm shaking. Cells from these precultures were then washed twice in 0.9% (w/v) NaCl and used to inoculate the bioreactor cultures at a starting OD_600_ of 0.1. The initial bioreactor culture volume was 500 mL of MOPS2 supplemented with 61 mM NaOAc and 0.01% (v v^-1^) antifoaming agent J647 (Struktol). Initial values for agitation speed and air sparging were 500 rpm and 0.5 L/min, respectively. To maintain the dissolved oxygen (DO) concentration in the bioreactors above 40%^92^, both parameters were controlled by an automated cascade (500-1200 rpm agitation and 0.5-2 L/min air sparging). The temperature set point was 37 °C. The culture pH was controlled at 7.2 by addition of HCl. After acetate depletion, the bioreactor cultures were fed following a DO-stat, acid-based, or hybrid feeding function, as specified, up to a final working volume of 1 L (unless otherwise stated). The feed solution contained 100 g/L NaOAc, 10 g/L (NH_4_)_2_SO_4_, 5 g/L MgSO_4_, 1.5 g/L K_2_HPO_4_, 16.4 mg/L FeSO_4_·7H_2_O, 1% (v v^-1^) micronutrient solution (as used for MOPS2), and 0.01 % (v v^-1^) antifoaming agent J647. Where specified, PHB biosynthesis was induced by adding anhydrotetracycline (ATc) to the culture medium and feed solution to a final concentration of 100 ng/mL. The reactors were sampled periodically following induction. Cell dry weight (CDW), extracellular acetate, and PHB were quantified from these samples as outlined above.

### Error propagation

To find the standard deviation of the observable 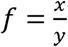 using the Gaussian error propagation, we use the propagation of uncertainty formula. Given two experimental observables *x* and *y* with their respective means *μ_x_* and *μ_y_* and standard deviations *σ_x_* and *σ_y_*, the formula for the standard deviation of *f* (denoted as *σ_f_*) is given as follows:

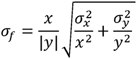

### Statistical reproducibility and analyses

Experiments other than transcriptomics analysis and fed-batch fermentation were repeated at least twice on different days to ensure statistical reproducibility. Statistical analyses for plate reader experiments and shake flask experiments were performed using a two-sample unpaired t-test method. Statistical analyses for data that have undergone error propagation is performed using one-sample t-test.

## Acknowledgements

This work was supported by funding from the Australian Research Council (ARC) through the Linkage Project grant LP220100185, Future Fellowship FT230100283, and Woodside Energy Ltd. The authors thank Prof. Terry Hwa for insightful discussions regarding adaptive evolution results. Method development was supported by valuable contributions from Prof. Nicolas Taylor and Ewen McCabe. Dr. Catherine Colas des Francs-Small, Dr. Joanna Melonek, and Gilang Fajar Suhorno are acknowledged for their assistance with RNA extraction. We extend our gratitude to Prof. Ian Small, Dr. James Lloyd, and Dr. Muhammad Kamran for their significant help with transcriptomic analysis. Macrogen Inc. and BGI Genomics performed library preparation and sequencing for RNAseq and genomic DNA samples, respectively. We appreciate Prof. Charlie Bond’s valuable suggestions regarding protein models. The authors also thank Dr. Brady Johnston and Dr. Philipp Bayer from UWA Hacky Hour for assistance in coding and data analysis. Dr. Ben Roller provided the protocols for measuring PHB via flow cytometry and shake flask experiments. We gratefully acknowledge Dr. Catherine Rinaldi for her valuable assistance in establishing and troubleshooting the flow cytometry methodology. Additionally, we thank Dr. Clara Woodcraft for the gDNA extraction methods and Katy Morris for her diligent laboratory support.

## Author Contributions

RJP: Conceptualization, Investigation, Methodology, Writing – Original Draft. SDV: Conceptualisation, Investigation, Methodology, Writing – Original Draft. LP: Investigation. JJ: Conceptualisation. GFl: Methodology. CE: Methodology. GFr: Supervision, Conceptualization, Methodology, Writing – Original Draft. All authors contributed to manuscript review and editing.

## Declaration of generative AI and AI-assisted technologies in the writing process

During the preparation of this work, the authors used ChatGPT (OpenAI, GPT-4) in order to refine scientific writing, improve clarity and logical flow, and assist in editing for grammar and style. After using this tool, the authors reviewed and edited the content as needed and take full responsibility for the content of the publication.

