## Supplementary Information for "Establishing *Vibrio natriegens* as a high-performance host for acetate-based poly-3-hydroxybutyrate production"

### Content

**Figure S1.** Growth kinetics of *V. natriegens* before and after ALE on MOPS2 medium.

**Figure S2.** Bioinformatics analysis of the mutations in *yheO* and DUF3612 accumulated during the ALE experiment.

**Figure S3.** Amino acid sequence alignment of *V. natriegens* BccT1 with other ectoine-transporting BCCT transporters in *V. parahaemolyticus* (VP1456) and *Corynebacterium glutanicum* (CgBetP).

**Figure S4.** Comparison of the BCCT transporters in *V. natriegens* compared to *V. parahaemolyticus* VP1456.

**Figure S5.** Characterization of the *bccT1* mutation.

**Figure S6.** Characterization of the *luxU* deletion.

**Figure S7.** PHB depolymerase knockout verification in AE2 strain and its mutants through plasmid based *pha* overexpression followed by carbon starvation.

**Figure S8.** DO-stat feeding in growth and PHB production experiments.

**Figure S9.** Acid-based feeding in growth and PHB production experiments.

**Figure S10.** Logic flow diagram of the hybrid feeding function.

**Table S1.** Locus tags of genes related to acetate metabolism, glyoxylate cycle, and TCA cycle.

**Table S2.** Genetic mutations in the three adaptively evolved (AE) strains.

**Table S3.** [not included – provided as separate file for download]

**Table S4.** Differential expression of the compatible-solute related genes and their locus tags in *V. natriegens* with the calculated $\log_2$ fold-change (log2FC) and adjusted p-value (padJ).

**Table S5.** Growth rate measured during RNAseq experiment.

**Table S6.** Differentially expressed genes in AE2 strain that are upregulated with a log2 fold change (log2FC) of >2 along with its adjusted p-value (padJ).

**Table S7.** Differentially expressed genes and their locus tags (LT) that are most downregulated in AE2 with a $\log_2$ fold change (log2FC) of <-2 along with its adjusted p-value (padJ).

**Table S8.** List of strains used in this study.

**Table S9.** Primers used in this study for the generation of the tDNA used in the genome engineering process.

**Table S10.** Oligonucleotides used to assemble the gRNA sequences for the NT-CRISPR plasmid.

**Table S11.** Oligonucleotides used for the verification of correct deletion through colony PCR and sanger sequencing.

**Table S12.** Oligonucleotides used to create Marburg Collection-compatible genetic parts in this study.

**Table S13.** Plasmids used in this study with their parts.

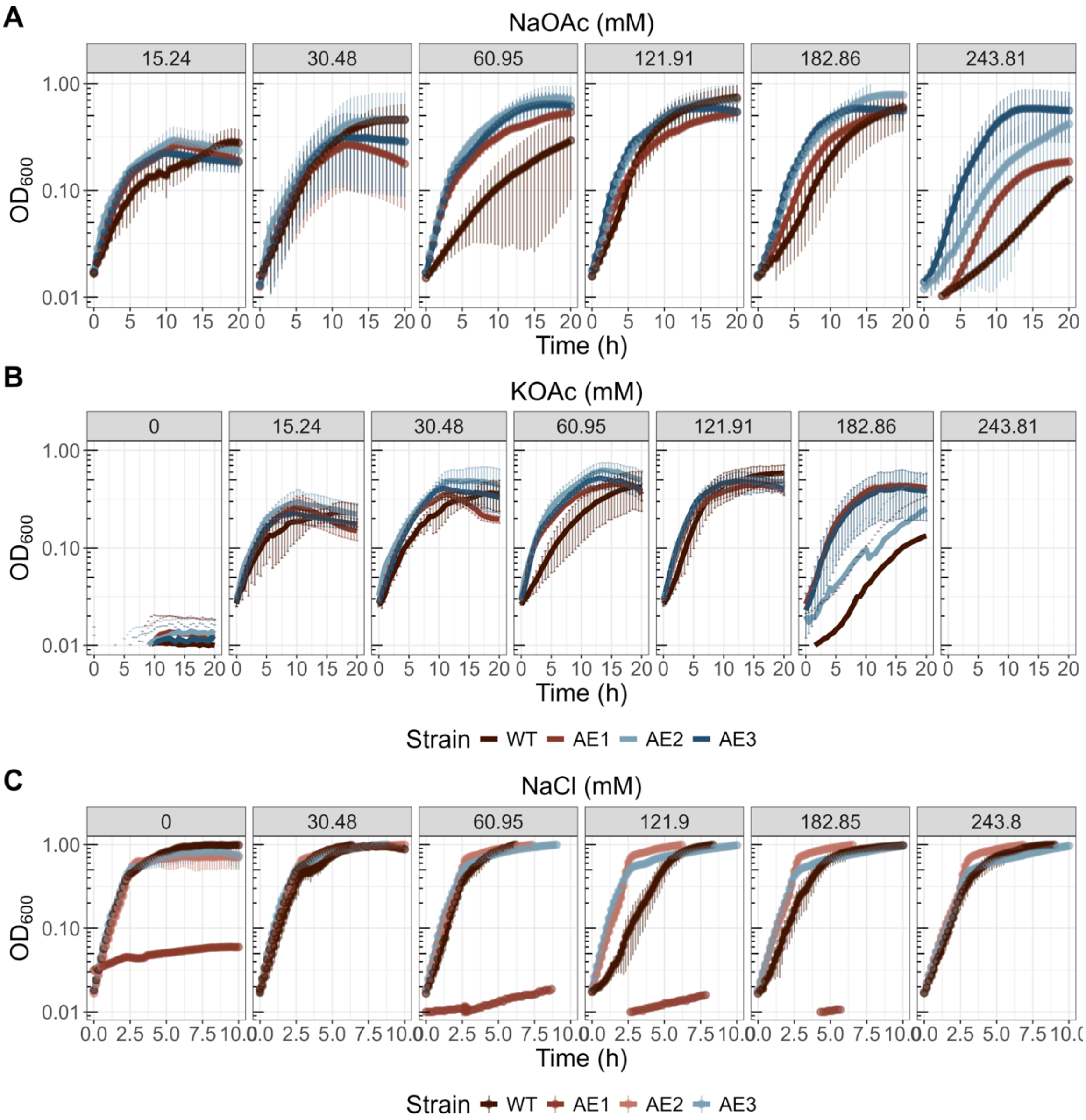


**Figure S1. Growth kinetics of *V. natriegens* before and after ALE on MOPS2 medium.** A) Growth curves with increasing NaOAc concentration, B) with increasing KOAc concentration, C) on 28 mM glucose with increasing NaCl concentration.

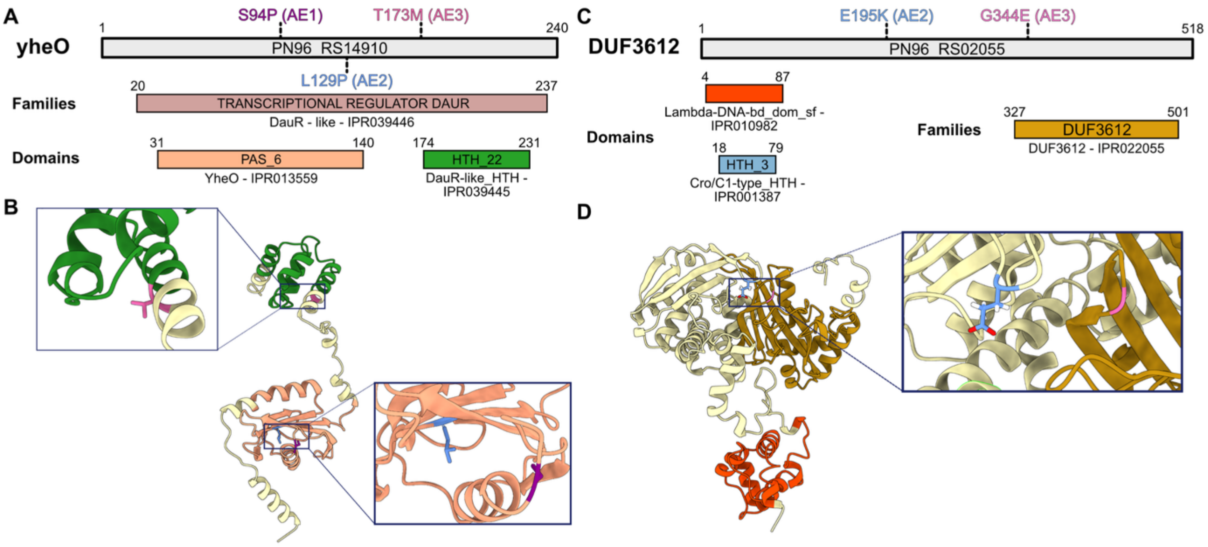


**Figure S2. Bioinformatics analysis of the mutations in YheO (A-B) and DUF3612 (C-D) accumulated during the ALE experiment. A, C)** Family and domain predictions were obtained from InterPro. **B, D)** AlphaFold3 protein models were generated and visualized using ChimeraX. The mutations in each AE strain were mapped onto the corresponding protein models to predict their effects. The mutations were found in distinct regions of the proteins for both YheO and DUF3612. For YheO, AE1 and AE2 carry mutations within the PAS_6 domain, while AE3 exhibits a mutation just upstream of the HTH_22 (DauR-like HTH) domain. These mutations suggest potential alterations in transcriptional activity. For DUF3612, the mutations in AE2 and AE3 occur in a similar region near the DUF3612 family, the function of which remains unknown.

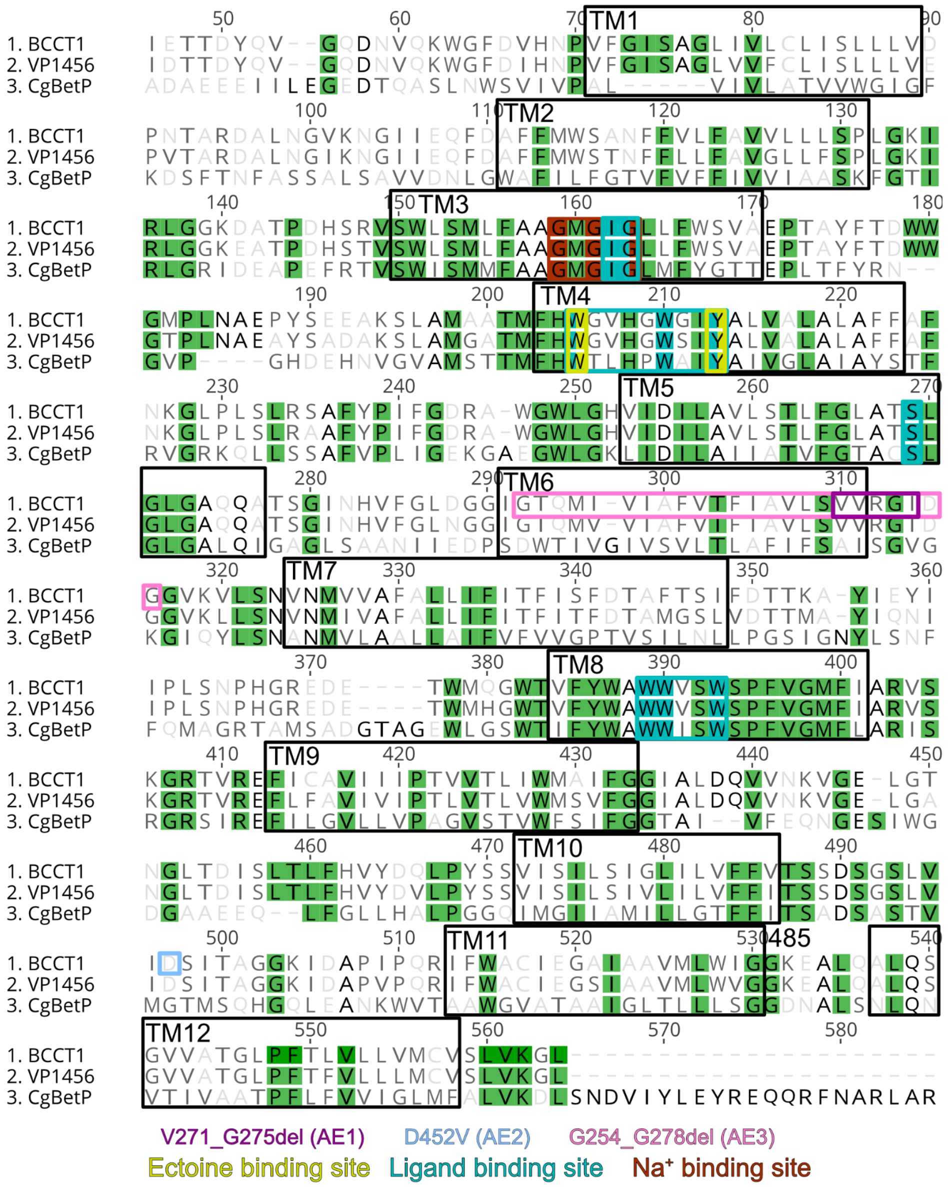


**Figure S3. Amino acid sequence alignment of *V. natriegens* BccT1 with other ectoine-transporting BCCT transporters in *V. parahaemolyticus* (VP1456) and *Corynebacterium glutanicum* (CgBetP).** Sequence alignment was performed using Geneious alignment with a cost matrix of Blosum62.

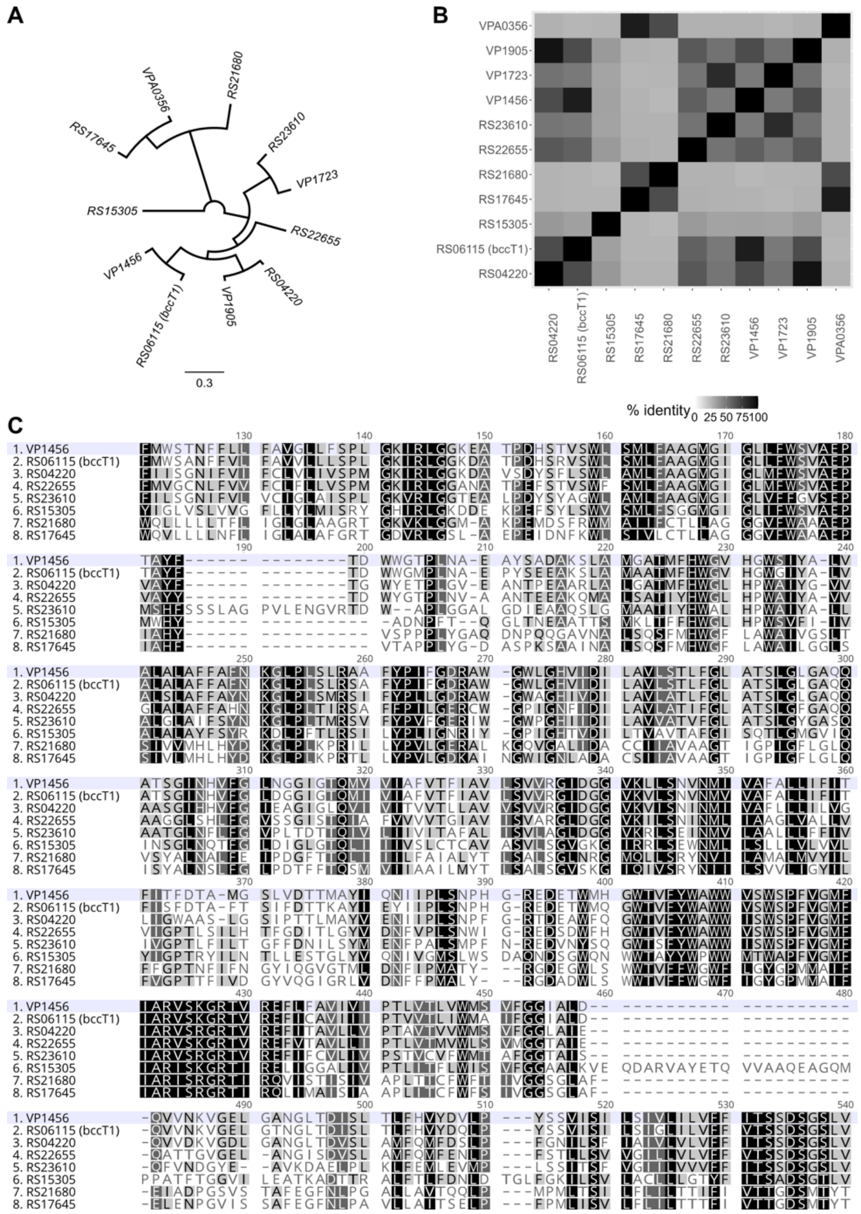


**Figure S4. Comparison of the BCCT transporters in *V. natriegens* compared to *V. parahaemolyticus* VP1456**. A) Phylogenetic tree of all the transporters. B) Heatmap of % amino acid identity between the BCCT transporters. C) Multiple sequence alignment showcasing the conserved amino acids. Sequence alignment was performed using Geneious alignment with a cost matrix of Blosum62.

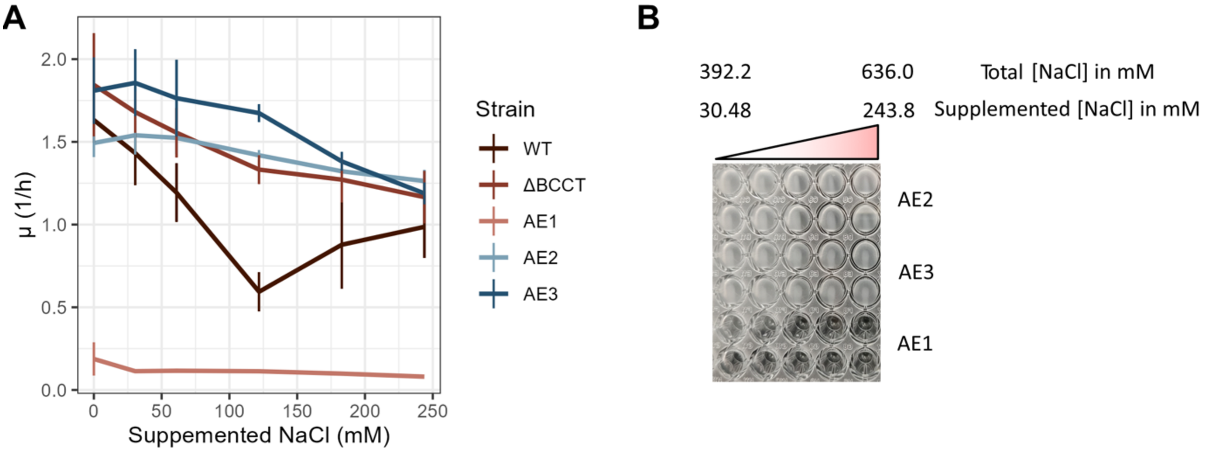


**Figure S5. Characterization of the *bccT1* mutation.** A) Comparison of growth rates of the WT, Δ*bccT1* mutant, and AE strains growing on 28 mM glucose supplemented with different titrations of NaCl. B) Image of 96-well plate used in this experiment showing similar phenotype to the Δ*bccT1* mutant and the absence of flocculation that was observed in the WT strain in **Fig. 4H**.

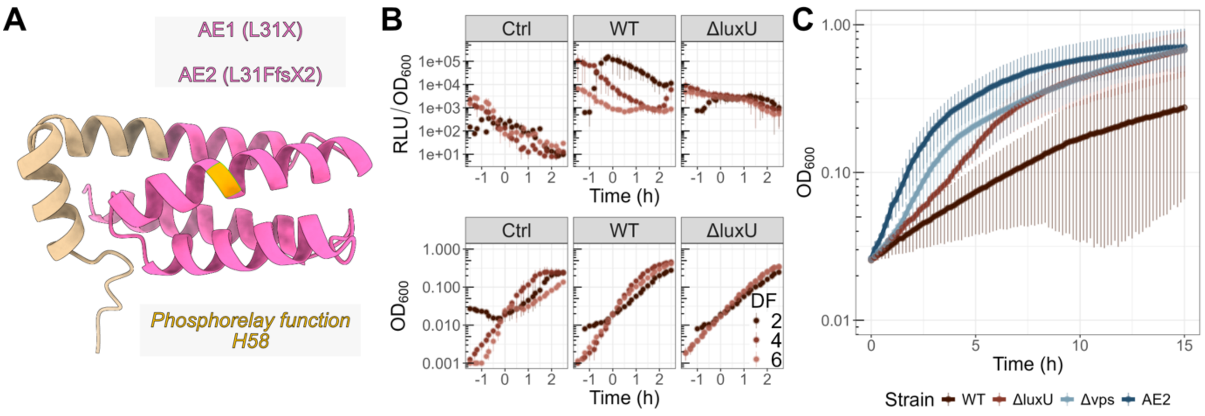


**Figure S6. Characterization of the *luxU* deletion.** A) Protein model of LuxU created with AlphaFold3 with the mutation in the AE strains. The strains harboring $P_{vps}$-luxCDABE biosensor from Figure 5C were grown similarly as described but were resuspended in 5.6 mM glucose containing MOPS2 medium instead of NaOAc. The growth and expression kinetics were measured with the plate reader, and $OD_{600}$ values below the detection limit (<0.001) were computationally mutated to 0.001 B) Relative luminescence intensity ($RLU/OD_{600}$) and growth kinetics of WT and Δ*luxU* strain transformed with a $P_{vps}$-luxCDABE biosensor on 5.6 mM glucose with 3 different dilution factors in $log_{10}$ (DF) (n = 6). An increase in $RLU/OD_{600}$ value was observed with a DF of 2 in the WT strain, and was observed minimally in the Δ*luxU* strain, reminiscent of the pattern observed in **Fig. 5C**. A decrease in $RLU/OD_{600}$ value was observed after reaching higher cell densities, suggesting dependency on cell density as observed in **Fig. 5C**. C) Growth kinetics of the mutants compared to WT on 61 mM NaOAc MOPS2 after a computational shift to $OD_{600}$ = 0.025 (n = 12).

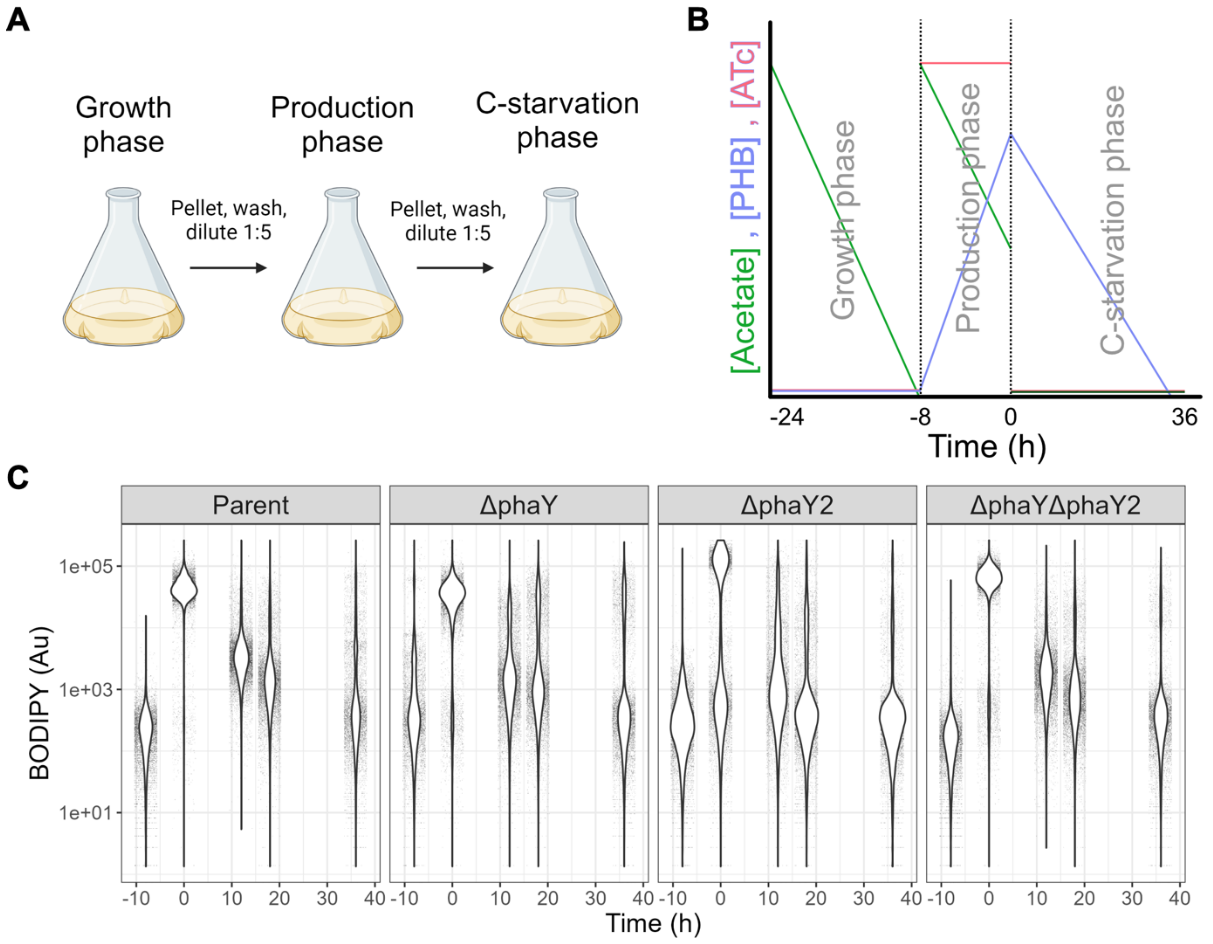


**Figure S7. PHB depolymerase knockout verification in AE2 strain (Parent) and its mutants through plasmid based *pha* overexpression followed by carbon starvation.** A) Flow diagram of the experiment. Cells were grown to $OD_{600}$ = 0.4-0.6 in 61 mM NaOAc MOPS2 prior to a centrifugation step (7,197x g for 10 minutes). The supernatant was removed, and the cells were resuspended in a fresh medium containing 100 ng/mL ATc and was incubated at 37 °C with shaking at 220 rpm for 8 hours to accumulate PHB. Afterwards, the pellet was isolated again and was washed and resuspended in C-deplete MOPS2 medium to trigger the PHB depolymerization for another 26 hours. B) Nutrient diagram of the experiment with the predicted outcome for cells with depolymerase. C) Measured BODIPY values before the plasmid overexpression (-8 h), after overexpression (0 h), and throughout the carbon starvation process (0-36 h).

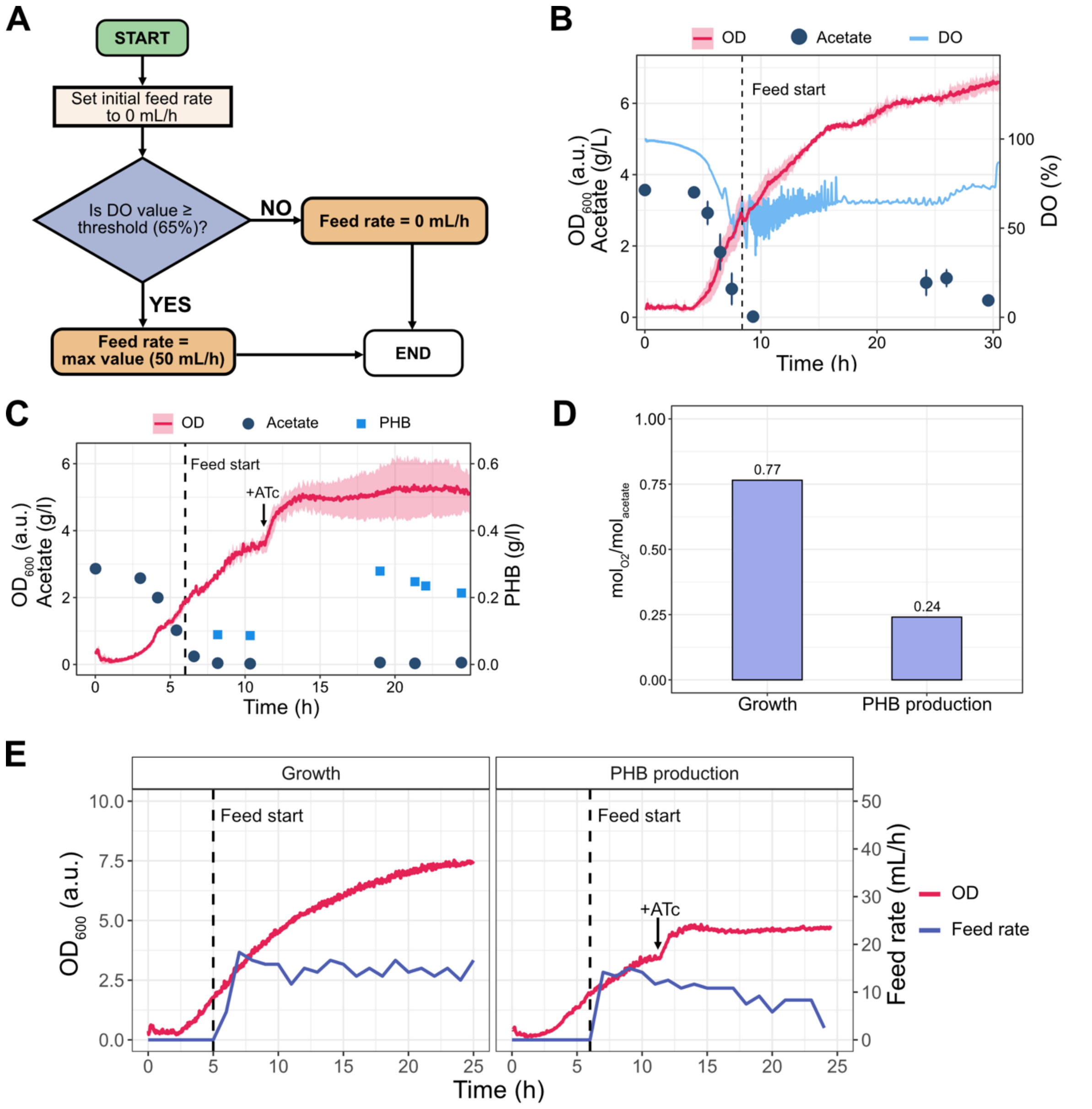


**Figure S8. DO-stat feeding in growth and PHB production experiments.** A) Logic flow diagram of the DO-stat feeding function, used to control feeding rate in the bioreactors. The function, implemented in Python, is executed once every second after the onset of feeding to detect spikes in the DO signal, indicative of carbon depletion. If the DO value surpasses a given threshold (65%), the feed rate is set to a maximum value of 50 mL/h (~3.5 g/h), providing a rapid pulse of substrate addition. Once the DO signal dips below the threshold, the feeding is stopped (i.e. the feed rate is set to 0 mL/h). Over the course of cultivation, the function ensures a self-adjusting feed rate which follows the biological demand. B) DO-stat fed-batch growth and C) PHB production experiments. The start of substrate feeding and inducer addition (where relevant) are indicated by a vertical dashed line and solid black arrow, respectively. The mean and standard deviation of the optical density ($OD_{600}$, measured online) are shown as a solid pink line and pink shading, respectively. At each sampling timepoint, the mean residual acetate (dark teal circles) and PHB (blue squares, where relevant) concentration is shown, with error bars indicating the standard deviation. For the growth experiments, a single representative DO trace is shown (solid blue line). All other data correspond to duplicate experiments. The final volume for all fermentations was 750 mL (initial volume 500mL, feeding solution volume 250mL). D) Genome-scale metabolic modelling predictions of oxygen consumption ($mol_{O2}/mol_{acetate}$) under growth and PHB production regimes. E) Representative feed rates for DO-stat fed-batch growth (left) and PHB production (right) experiments. For simplicity, a single OD (solid pink line) and feed rate (solid dark blue line) are shown for each type of experiment.

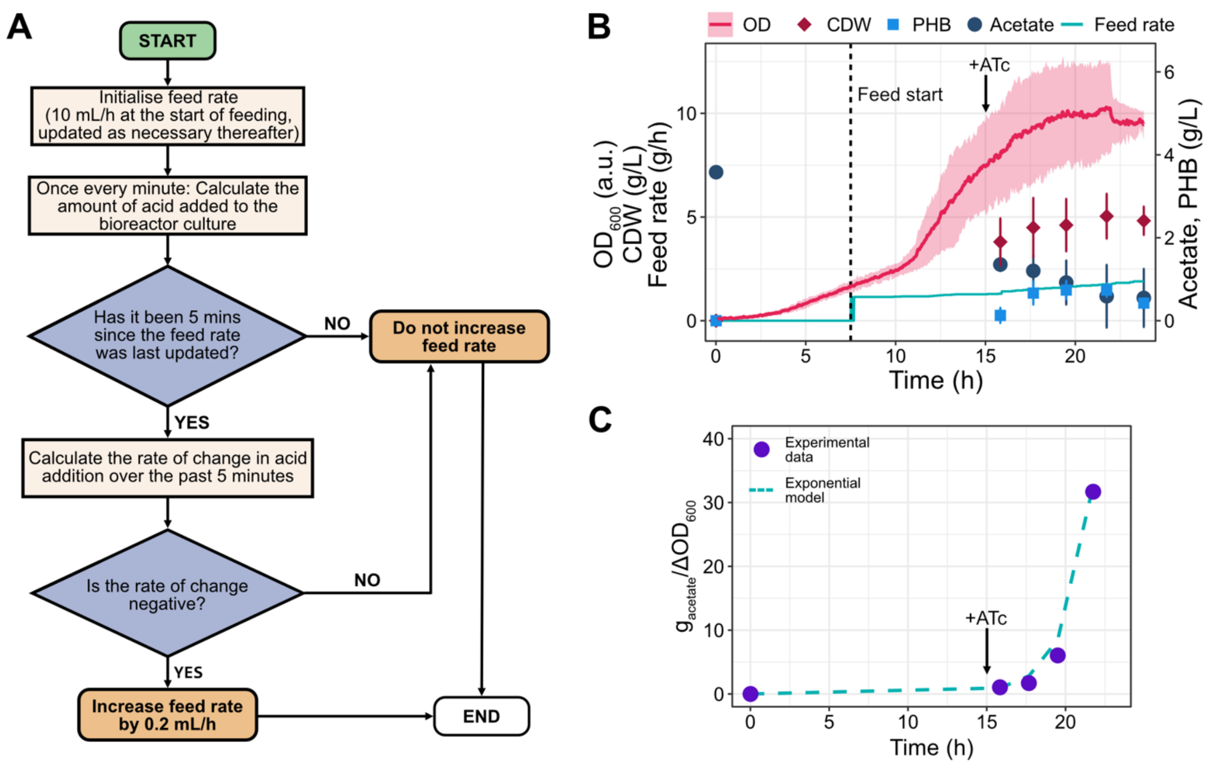


**Figure S9. Acid-based feeding in growth and PHB production experiments.** A) Logic flow diagram of the acid-based feeding function, used to control feeding rate in the bioreactors. The function, implemented in Python, is executed once every second after the onset of feeding, but new feeding values are only computed every five minutes to minimize the effect of random fluctuations. Namely, every five minutes, the rate of change in the volume of acid added per minute (mL/min) is computed. A negative rate of change will trigger an increase in the feeding rate (+ 0.2 mL/h). B) Acid-based feeding PHB production experiment. The start of substrate feeding and inducer addition are indicated by a vertical dashed line and solid black arrow, respectively. The mean and standard deviation of the optical density (OD, measured online) are shown as a solid pink line and pink shading, respectively. At each sampling timepoint, the mean CDW (red diamonds), residual acetate (dark teal circles), and PHB (blue squares, where relevant) concentration is shown, with error bars indicating the standard deviation. For simplicity, a single representative feed rate trace is shown (solid green line). All other data correspond to duplicate experiments. The final volume for all fermentations was 750 mL (initial volume 500mL, feeding solution volume 250mL). C) Acetate consumption increases exponentially following induction. An exponential growth model is fitted to experimental data points derived from one of the PHB production experiments shown in C).

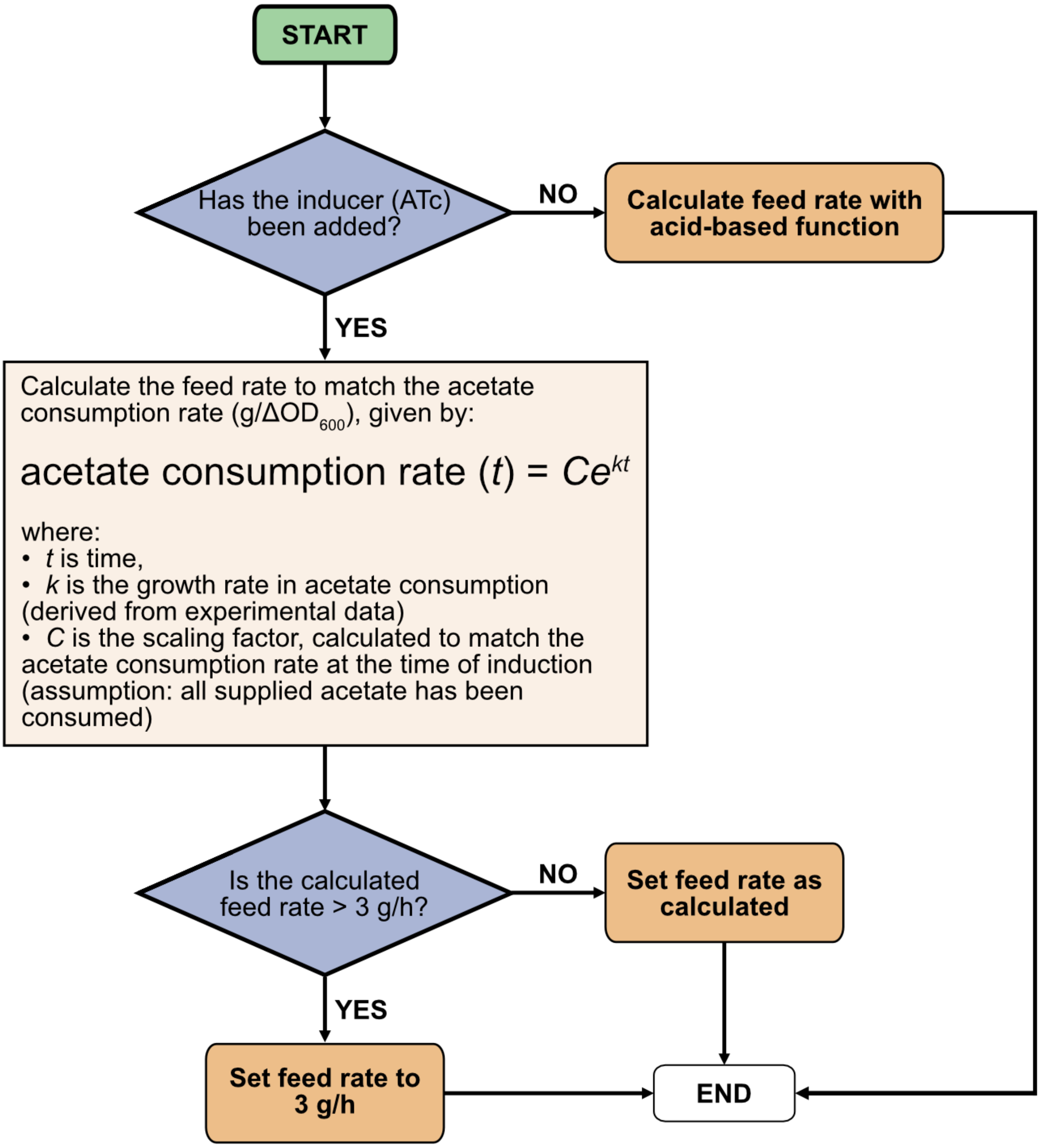


**Figure S10. Logic flow diagram of the hybrid feeding function.** The function, implemented in Python, is executed once every second after the onset of feeding. If the inducer (ATc) has not yet been added to the cultures, an input provided by the user during the cultivation, the feed rate is calculated using the acid-based feeding function. Following induction, the feed rate at the time of substrate addition is used to calculate the scaling factor C, anchoring the exponential feeding function to the current feed rate value. At every iteration of the function, a new feed rate is calculated using the exponential function. The feed rate is capped at a maximum value of 3 g/h to prevent excessive accumulation of residual acetate in the bioreactors.

**Table S1. Locus tags of genes related to acetate metabolism, glyoxylate cycle, and TCA cycle**. Locus tags from the Refseq and Genbank databases are annotated as PN96_RSXXXXX and PN96_XXXXX, respectively, with XXXXX indicating the locus listed in the table.

| Locus tag (Refseq) [PN96_RS] | Locus tag (Genbank) [PN96_] | Description (protein/gene name) | Gene |
|---|---|---|---|
| **Acetyl-CoA synthesis** | | | |
| 14410 | 14210 | *acsA1* | *acsA1* |
| 03410 | 03360 | *pta1* | *pta1* |
| 03415 | 03365 | acetate kinase | *ackA1* |
| 21805 | 21510 | acetate/propionate family kinase | *ackA2* |
| 22305 | 22000 | *pta2* | *pta2* |
| 21410 | 21120 | *acsA2* | *acsA2* |
| **Glyoxylate cycle** | | | |
| 10725 | 10580 | *aceA* | *aceA* |
| 10730 | 10585 | malate synthase A | *aceB1* |
| 17175 | 16960 | malate synthase | *aceB2* |
| 22310 | 22005 | IclR family transcriptional regulator | - |
| 22730 | 22415 | IclR family transcriptional regulator | - |
| 22310 | 22005 | IclR family transcriptional regulator | - |
| **Acetate transporter** | | | |
| 14445 | 14245 | cation acetate symporter | *actP* |
| **Tricarboxylic acid cycle** | | | |
| 09420 | 09305 | citrate synthase | *gltA* |
| 01460 | 01430 | *acnB* | *acnB* |
| 08465 | 08360 | NADP-dependent isocitrate dehydrogenase | *icd* |
| 01375 | 01345 | *lpdA* | *lpdA* |

| | | | |
|---|---|---|---|
| 09390 | 09275 | *odhB* | *sucB* |
| 09395 | 09280 | *sucA* | *sucA* |
| 09380 | 09265 | *sucD* | *sucD* |
| 09385 | 09270 | *sucC* | *sucC* |
| 09400 | 09285 | succinate dehydrogenase iron-sulfur subunit | *sdhB* |
| 09405 | 09290 | *sdhA* | *sdhA* |
| 09410 | 09295 | *sdhD* | *sdhD* |
| 09415 | 09300 | *sdhC* | *sdhC* |
| 14570 | 14370 | *frdD* | *frdD* |
| 14575 | 14375 | *frdC* | *frdC* |
| 14580 | 14380 | succinate dehydrogenase/fumarate reductase iron-sulfur subunit | *frdB* |
| 14585 | 14385 | *frdA* | *frdA* |
| 04300 | 04250 | fumarate hydratase | *fumA* |
| 14435 | 14235 | class II fumarate hydratase | *fumC* |
| 06560 | 06470 | *mqo* | - |
| 19725 | 19465 | *mqo* | - |
| 11855 | 11695 | *mdh* | *mdh* |
| 07395 | 07295 | NAD-dependent malic enzyme | *maeA* |
| 14960 | 14755 | malate dehydrogenase | *maeB* |
| 12890 | 12720 | *pckA* | *pckA* |
| 14995 | 14785 | *ppc* | *ppc* |
| 01365 | 01335 | *aceE* | *aceE* |
| 01370 | 01340 | *aceF* | *aceF* |
| 01375 | 01345 | *lpdA* | *lpdA* |
| 21420 | 21130 | 2-oxo acid dehydrogenase subunit E2 | - |

| | | | |
|---|---|---|---|
| 22095 | 21795 | *lpdA* | - |
| 17695 | 17465 | *ppsA* | *ppsA* |
| 21430 | 21140 | *ppsA* | - |
| 03630 | 03585 | *pyk* | *pykA* |
| 11710 | 11555 | *pykF* | *pykF* |
| 01105 | 01075 | *eno* | *eno* |

**Table S2. Genetic mutations in the three adaptively evolved (AE) strains.** For each of the evolved cell lines AE1, AE2, and AE3, two independent clones were sequenced (indicated by the number after the dot) and the shared mutations are denoted. Locus tags from the Refseq and Genbank databases are annotated as PN96_RSXXXXX and PN96_XXXXX, respectively, with XXXXX indicating the locus listed in the table. Abbreviation: coding sequence (CDS)

| **Locus tag (Refseq) [PN96_RS]** | **Locus tag (Genbank) [PN96_]** | **Target description (protein/gene name)** | **Location** | **Strain and mutation** |
|---|---|---|---|---|
| 01365 | 01335 | *aceE* | CDS | AE1.1, AE1.2 (N589K) |
| 06115 | 06025 | *bccT1* | CDS | AE1.1, AE1.2 (ΔV271-G275), AE2.1, AE2.2 (D452V), AE3.1, AE3.2 (ΔG254-G278) |
| 17170 | 16955 | DUF3612 domain-containing protein | CDS | AE2.1, AE2.2 (E195K), AE3.1, AE3.2 (G344E) |
| 14185 | 13995 | *ftsY* | CDS | AE2.1, AE2.2 (E81AfsX2) |
| 08045 | 07940 | *lrp* | CDS | AE1.1 (D114G) |
| 03340 | 03290 | *luxU* | CDS | AE1.1, AE1.2 (L31X), AE2.1, AE2.2 (L31FfsX2) |
| 00390 | NA | hypothetical protein | CDS | AE3.1, AE3.2 (C275Y) |
| 12405 | 12240 | NAD-dependent epimerase/dehydratase family protein | CDS | AE3.1, AE3.2 (G258GfsX12) |
| 19835 | 19575 | *nhaC* | CDS | AE1.2 (M108RfsX11) |
| 10715 | 10570 | serine protease | CDS | AE2.1, AE2.2 (P194S) |
| 14910 | 14710 | *yheO* | CDS | AE1.1 (S94P), AE2.1, AE2.2 (L129P), AE3.1, AE3.2 (T173M) |
| 03035 | 02985 | *yeiW* | CDS | AE3.1, AE3.2 (G65E) |
| 05290 | 05210 | *proV* | Intergenic (18 bp upstream) | AE3.1, AE3.2 (2 bp deletion) |
| 13840 | 13660 | *thiC* | Intergenic (1 bp upstream) | AE2.1, AE2.2 (1 bp deletion) |

**Table S4. Differential expression of the compatible-solute related genes and their locus tags in *V. natriegens* with the calculated $log_2$ fold-change (log2FC) and adjusted p-value (padJ).** Locus tags from the Refseq and Genbank databases are annotated as PN96_RSXXXXXX and PN96_XXXXX, respectively, with XXXXX indicating the locus listed in the table. The full dataset is shown in Table S3.

| **Locus tag** | | **Gene** | **log2FC** | | | **padJ** | | |
|---|---|---|---|---|---|---|---|---|
| **Refseq [PN96_RS]** | **Genbank [PN96_]** | | **AE2(0.5% NaOAc)/ WT(0.5% NaOAc)** | **AE3(0.5% NaOAc)/ WT(0.5% NaOAc)** | **WT(0.5% NaOAc)/ WT(0.5% Glucose)** | **AE2(0.5% NaOAc)/ WT(0.5% NaOAc)** | **AE3(0.5% NaOAc)/ WT(0.5% NaOAc)** | **WT(0.5% NaOAc)/ WT(0.5% Glucose)** |
| 18285 | 18045 | *phaB* | -0.31 | -1.58 | 7.27 | 0.20 | 7.97e-14 | 2.55e-213 |
| 18290 | 18050 | *phaA* | -0.15 | -1.35 | 6.97 | 0.49 | 2.65e-12 | 4.01e-190 |
| 18295 | 18055 | *phaP* | -0.48 | -1.45 | 7.06 | 0.02 | 4.15e-12 | 3.71e-221 |
| 18300 | 18060 | *phaC* | -0.20 | -1.21 | 6.73 | 0.40 | 4.49e-8 | 4.74e-205 |
| 19345 | 19085 | *betI* | -0.02 | 0.39 | 0.21 | 0.96 | 0.34 | 0.58 |
| 19350 | 19090 | *betB* | -0.29 | 0.15 | 1.07 | 0.16 | 0.53 | 0.00 |
| 19355 | 19095 | *betA* | -0.04 | 0.28 | 1.20 | 0.91 | 0.36 | 0.00 |
| 19360 | 19100 | *choX* | -0.13 | 0.03 | 0.34 | 0.78 | 0.96 | 0.50 |
| 19365 | 19105 | *choW* | -0.36 | 0.05 | 0.79 | 0.38 | 0.92 | 0.03 |
| 19370 | 19110 | *choV* | -0.39 | 0.12 | 1.13 | 0.33 | 0.81 | 0.00 |
| 10100 | 09965 | *treC* | -0.17 | -0.42 | 0.41 | 0.56 | 0.13 | 0.24 |
| 10105 | 09970 | *treB* | -0.42 | -0.45 | 1.84 | 0.26 | 0.26 | 0.00 |
| 10110 | 09975 | *treR* | 0.86 | 0.76 | -1.10 | 0.00 | 0.00 | 0.00 |
| 05295 | 05215 | *ectA* | 2.14 | 2.36 | 2.35 | 0.00 | 0.00 | 0.00 |
| 05300 | 05220 | *ectB* | 1.75 | 2.11 | 2.19 | 0.00 | 0.00 | 0.00 |
| 05305 | 05225 | *ectC* | 1.57 | 2.04 | 2.24 | 0.00 | 0.00 | 0.00 |
| 05310 | 05230 | *aspC* | 2.02 | 1.83 | 2.56 | 0.00 | 0.00 | 0.00 |

**Table S5. Growth rate measured during RNAseq experiment.** Cultures were grown in 100 mL MOPS2 medium with 0.5% (61 mM) NaOAc or 0.5% (28 mM) glucose in 1 L baffled flask at 37 ºC 180 rpm. $OD_{600}$ was measured with a hand-held Cell density meter model 40 (Fisher Scientific). The number after the dot represents the biological replicate.

| Strain | Media | GR (1/h) | Average | Standard dev |
|---|---|---|---|---|
| AE2.1 | 0.5% NaOAc MOPS2 | 0.443 | | |
| AE2.2 | 0.5% NaOAc MOPS2 | 0.472 | 0.483 | 0.038 |
| AE2.3 | 0.5% NaOAc MOPS2 | 0.534 | | |
| AE3.1 | 0.5% NaOAc MOPS2 | 0.438 | | |
| AE3.2 | 0.5% NaOAc MOPS2 | 0.443 | 0.459 | 0.026 |
| AE3.3 | 0.5% NaOAc MOPS2 | 0.496 | | |
| WT1 (Ac1) | 0.5% NaOAc MOPS2 | 0.334 | | |
| WT2 (Ac2) | 0.5% NaOAc MOPS2 | 0.319 | 0.320 | 0.011 |
| WT3 (Ac3) | 0.5% NaOAc MOPS2 | 0.308 | | |
| Glu1 | 0.5% Glucose MOPS2 | 1.436 | | |
| Glu2 | 0.5% Glucose MOPS2 | 1.371 | 1.427 | 0.043 |
| Glu3 | 0.5% Glucose MOPS2 | 1.474 | | |

**Table S6. Differentially expressed genes in AE2 strain that are upregulated with a log2 fold change (log2FC) of >2 along with its adjusted p-value (padJ).** Locus tags from the Refseq and Genbank databases are annotated as PN96_RSXXXXXX and PN96_XXXXX, respectively, with XXXXX indicating the locus listed in the table. The full dataset is shown in Table S3.

| **Locus tag** | | **Description** | **Gene** | **log2FC** | | | **padJ** | | |
|---|---|---|---|---|---|---|---|---|---|
| **Refseq [PN96_RS]** | **Genbank [PN96_]** | | | **AE2(0.5% NaOAc)/ WT(0.5% NaOAc)** | **AE3(0.5% NaOAc)/ WT(0.5% NaOAc)** | **WT(0.5% NaOAc)/ WT(0.5% Glucose)** | **AE2(0.5% NaOAc)/WT(0.5% NaOAc)** | **AE3(0.5% NaOAc)/WT(0.5% NaOAc)** | **WT(0.5% NaOAc)/WT(0.5% Glucose)** |
| 01485 | 01455 | P-II family nitrogen regulator | *glnK* | 5.10 | 4.57 | -4.69 | 7.43e-89 | 1.87e-81 | 1.62e-60 |
| 12320 | 12155 | *tviC* | *tviC* | 4.24 | 0.30 | 1.29 | 3.4e-34 | 0.64 | 0.01 |
| 12325 | 12160 | four helix bundle protein | NA | 4.07 | 1.04 | 1.95 | 5.02e-53 | 0.04 | 2.1e-6 |
| 21805 | 21510 | acetate/propionate family kinase | *ackA2* | 4.04 | 3.35 | -1.51 | 2.07e-71 | 4.66e-55 | 2.01e-10 |
| 12265 | 12100 | polysaccharide biosynthesis protein | - | 3.96 | 0.05 | -0.45 | 2.38e-69 | 0.90 | 0.48 |
| 12335 | 12170 | polysaccharide biosynthesis tyrosine autokinase | *wzc* | 3.78 | 0.60 | -0.23 | 2.89e-92 | 0.01 | 0.39 |
| 12295 | 12130 | glycosyltransferase | - | 3.75 | 0.36 | 1.41 | 8.88e-13 | 0.64 | 0.09 |
| 12330 | 12165 | *tviB* | *tviB* | 3.74 | 0.38 | 1.42 | 1.73e-33 | 0.38 | 1.20e-4 |
| 12280 | 12115 | sugar transferase | - | 3.69 | 0.13 | 0.62 | 9.42e-43 | 0.79 | 0.10 |
| 12270 | 12105 | DegT/DnrJ/EryC1/StrS aminotransferase family protein | - | 3.69 | 0.08 | -0.97 | 1.97e-104 | 0.82 | 0.02 |
| 12305 | 12140 | hypothetical protein | - | 3.59 | 0.72 | 1.29 | 1.55e-15 | 0.23 | 0.07 |
| 01490 | 01460 | ammonium transporter | *amtB* | 3.53 | 3.43 | -3.36 | 1.04e-101 | 1.89e-68 | 8.87e-38 |
| 12300 | 12135 | glycosyltransferase family 4 protein | - | 3.52 | 0.01 | 2.18 | 1.69e-11 | 0.99 | 0.00 |

| | | | | | | | | | |
|---|---|---|---|---|---|---|---|---|---|
| 12290 | 12125 | glycosyltransferase | - | 3.48 | -0.01 | 2.45 | 1.19e-13 | 0.99 | 5.59875e-4 |
| 12315 | 12150 | flippase | *rfbX* | 3.48 | -0.20 | 3.15 | 3.14e-19 | 0.80 | 3.5e-8 |
| 13950 | 13765 | 7-cyano-7-deazaguanine/7-aminomethyl-7-deazaguanine transporter | *yhhQ* | 3.48 | 2.82 | 0.41 | 1.75e-28 | 1.61e-20 | 0.27 |
| 12275 | 12110 | acetyltransferase | - | 3.42 | 0.17 | 0.39 | 1.06e-25 | 0.75 | 0.36 |
| 09320 | 09205 | trypsin-like serine protease | - | 3.35 | 0.08 | 0.85 | 2.96e-80 | 0.75 | 0.00 |
| 23555 | 23235 | DUF3015 family protein | - | 3.33 | 1.06 | -1.52 | 2.06e-50 | 1.82e-7 | 2.7e-7 |
| 12345 | 12180 | polysaccharide export protein | *wza* | 3.33 | 0.26 | -0.57 | 2.08e-77 | 0.29 | 0.02 |
| 12285 | 12120 | hypothetical protein aldolase/citrate lyase | *aceB* | 3.26 | 0.11 | 2.23 | 1.03e-17 | 0.88 | 1.17137e-4 |
| 12340 | 12175 | low molecular weight phosphotyrosine protein phosphatase | *wzb* | 3.24 | 0.40 | -0.65 | 6.43e-55 | 0.08 | 0.01 |
| 10555 | 10410 | hypothetical protein | *yaaH* | 3.21 | 0.13 | -5.21 | 1.10e-27 | 0.79 | 1.41e-87 |
| 01380 | 01350 | TetR/AcrR family transcriptional regulator | *luxR* | 3.16 | 0.02 | 0.61 | 2.26e-32 | 0.96 | 0.02 |
| 23560 | 23240 | DUF4105 domain-containing protein | *NA* | 3.04 | 1.23 | -1.09 | 3.42e-27 | 3.33e-5 | 0.01 |
| 12310 | 12145 | hypothetical protein | - | 3.03 | 0.12 | 1.98 | 8.47e-5 | 0.89 | 0.02 |
| 12260 | 12095 | MBL fold metallo-hydrolase | - | 2.94 | -0.15 | 1.69 | 3.16e-37 | 0.68 | 4.55e-7 |
| 09325 | 09210 | hypothetical protein | - | 2.81 | 0.12 | 0.54 | 1.59e-59 | 0.72 | 0.04 |
| 19550 | 19295 | EamA family transporter | *VPA1529* | 2.75 | 4.04 | 2.09 | 1.81e-24 | 2.43e-71 | 3.39e-7 |

| | | | | | | | | | |
|---|---|---|---|---|---|---|---|---|---|
| 12940 | 12770 | *glnL* | *glnL* | 2.70 | 2.42 | -2.60 | 2.22e-41 | 8.74e-31 | 5.77e-25 |
| 21725 | 21430 | *cyoA* | *cyoA* | 2.58 | 2.11 | -7.63 | 1.94e-19 | 2.95e-19 | 5.2e-202 |
| 00375 | 00370 | type II secretion system protein | *mshD* | 2.43 | 1.92 | -1.28 | 1.11e-26 | 8.03e-22 | 5.54e-9 |
| 03415 | 03365 | acetate kinase | *ackA1* | 2.40 | 0.30 | -2.29 | 2.11e-43 | 0.16 | 1.72e-31 |
| 05910 | 05815 | ammonium transporter | - | 2.40 | 2.31 | -2.68 | 5.41e-18 | 3.09e-18 | 2.64e-14 |
| 00370 | 00365 | prepilin-type N-terminal cleavage/methylation domain-containing protein | *mshC* | 2.37 | 2.00 | -1.01 | 7.23e-24 | 1.88e-31 | 6.64e-7 |
| 10080 | 09945 | *lipB* | *lipB* | 2.34 | 1.28 | -0.95 | 2.72e-31 | 5.26e-14 | 7.85e-6 |
| 00385 | 00380 | hypothetical protein | - | 2.32 | 1.97 | -0.65 | 5.96e-18 | 8.4e-35 | 0.01 |
| 21720 | 21425 | *cyoB* | *cyoB* | 2.30 | 1.99 | -7.32 | 5.16e-10 | 1.46e-9 | 2e-104 |
| 03410 | 03360 | *pta1* | *pta1* | 2.24 | 0.35 | -1.93 | 2.18e-18 | 0.27 | 1.56e-12 |
| 00360 | 00355 | type II secretion system protein | - | 2.21 | 1.68 | -2.33 | 1.64e-16 | 2.35e-15 | 1.20e-39 |
| 05295 | 05215 | *ectA* | *ectA* | 2.14 | 2.36 | 2.35 | 5.54e-9 | 1.58e-18 | 6.13e-10 |
| 00380 | 00375 | prepilin-type N-terminal cleavage/methylation domain-containing protein | - | 2.06 | 1.79 | -1.10 | 6.78e-16 | 7.97e-14 | 4.3e-5 |
| 00600 | 00590 | hypothetical protein | - | 2.03 | 0.94 | -1.13 | 1.48e-11 | 4.42e-5 | 3.29e-5 |
| 05310 | 05230 | aspartate kinase | - | 2.02 | 1.83 | 2.56 | 3.19e-6 | 6.66e-10 | 1.52e-14 |

**Table S7. Differentially expressed genes and their locus tags (LT) that are most downregulated in AE2 with a $log_2$ fold change (log2FC) of <-2 along with its adjusted p-value (padJ).** Locus tags from the Refseq and Genbank databases are annotated as PN96_RSXXXXX and PN96_XXXXX, respectively, with XXXXX indicating the locus listed in the table. The full dataset is shown in Table S3.

| Locus tag | | Description | Gene | log2FC | | | padJ | | |
|---|---|---|---|---|---|---|---|---|---|
| Refseq [PN96_RS] | Genbank [PN96_] | | | AE2(0.5% NaOAc)/ WT(0.5% NaOAc) | AE3(0.5% NaOAc)/ WT(0.5% NaOAc) | WT(0.5% NaOAc)/ WT(0.5% Glucose) | AE2(0.5% NaOAc)/ WT(0.5% NaOAc) | AE3(0.5% NaOAc)/ WT(0.5% NaOAc) | WT(0.5% NaOAc)/ WT(0.5% Glucose) |
| 15355 | 15145 | calcium-binding protein | - | -5.90 | -2.34 | 7.53 | 1.12e-99 | 4.02e-13 | 1.8e-253 |
| 14420 | 14220 | cyclic nucleotide-binding/CBS domain-containing protein | - | -5.63 | -0.95 | 6.45 | 6.96e-80 | 0.02 | 3.0e-61 |
| 03995 | 03950 | calcium-binding protein | - | -5.46 | -1.77 | 7.03 | 6.5e-75 | 1.28e-32 | 2.27e-70 |
| 19210 | 18950 | molybdopterin-dependent oxidoreductase | - | -5.43 | -1.10 | 4.71 | 3.98e-63 | 4.9e-4 | 9.04e-49 |
| 15215 | 15005 | glycosyltransferase | - | -5.25 | -1.60 | 7.73 | 2.97e-77 | 1.11e-6 | 7.29e-112 |
| 14410 | 14210 | *acsA1* | *acsA1* | -5.23 | -1.15 | 7.09 | 2.28e-74 | 0.00 | 1.62e-118 |
| 23045 | 22730 | hypothetical protein | - | -5.22 | -1.55 | 1.27 | 7.52e-45 | 6.06e-4 | 4.22e-4 |
| 14450 | 14250 | DUF4212 domain-containing protein | - | -4.79 | 0.75 | 6.72 | NA | 0.27 | NA |
| 15205 | 14995 | polysaccharide biosynthesis tyrosine autokinase | - | -4.79 | -1.36 | 8.07 | 2.21e-38 | 1.65979e-4 | 1.03e-119 |
| 15230 | 15020 | glycosyltransferase | - | -4.78 | -1.38 | 6.68 | 5.44e-37 | 0.00 | 7.75e-59 |
| 23895 | NA | hypothetical protein | - | -4.75 | -0.86 | 5.12 | 3.13e-56 | 0.02 | 9.75e-102 |
| 15225 | 15015 | putative capsular | - | -4.66 | -1.30 | 7.51 | 8.03e-33 | 0.01 | 1.84e-58 |

| | | | | | | | | | |
|---|---|---|---|---|---|---|---|---|---|
| | | polysaccharide synthesis family protein | | | | | | | |
| 15210 | 15000 | putative capsular polysaccharide synthesis family protein | - | -4.66 | -1.26 | 6.99 | 6.24e-39 | 0.00 | 1.42e-55 |
| 15195 | 14985 | outer membrane beta-barrel protein | - | -4.61 | -1.52 | 7.85 | 1.07e-44 | 7.43e-8 | 7.37e-94 |
| 15780 | 15565 | DUF1232 domain-containing protein | - | -4.58 | -1.96 | 2.22 | 2.15e-44 | 1.03e-6 | 8.89e-4 |
| 14415 | 14215 | 3-5 exonuclease | - | -4.58 | -0.74 | 5.52 | 3.62e-31 | 0.12 | 4.68e-30 |
| 15220 | 15010 | O-antigen ligase family protein | NA | -4.56 | -1.25 | 7.17 | 3.41e-37 | 0.00 | 5.88e-72 |
| 21620 | 21325 | FAD-binding oxidoreductase | - | -4.47 | -1.35 | 4.30 | 1.34e-70 | 1.38e-7 | 1.1e-60 |
| 15190 | 14980 | undecaprenyl-phosphate glucose phosphotransferase | NA | -4.42 | -1.46 | 8.57 | 1.81e-36 | 6.98e-5 | 1.24e-101 |
| 15240 | 15030 | *vanZ* | - | -4.38 | -1.17 | 5.08 | 2.95e-40 | 0.01 | 1.82e-35 |
| 16035 | 15815 | type III PLP-dependent enzyme | - | -4.34 | -0.69 | 6.31 | 1.42e-7 | 0.33 | 5.8e-10 |
| 15350 | 15140 | type I secretion system permease/ATPase | - | -4.30 | -1.39 | 5.02 | 3.06e-29 | 3.06e-4 | 2.59e-32 |
| 21295 | 21005 | DUF2235 domain-containing protein | - | -4.27 | -3.00 | 3.75 | 3.64e-22 | 8.62e-24 | 1.01e-15 |
| 15200 | 14990 | polysaccharide export protein | - | -4.27 | -1.26 | 7.02 | 2.25e-27 | 8.5e-5 | 3.83e-47 |

| | | | | | | | | | |
|---|---|---|---|---|---|---|---|---|---|
| 15235 | 15025 | oligosaccharide flippase family protein | - | -4.22 | -1.09 | 5.82 | 1.05e-30 | 0.02 | 2.01e-47 |
| 07220 | 07120 | phosphoribosyl glycinamide formyltransfera se | - | -4.07 | -2.71 | 4.38 | 3.81e-25 | 2.25e-11 | 3.41e-25 |
| 19205 | 18945 | c-type cytochrome | - | -4.07 | -0.95 | 5.32 | 4.03e-22 | 0.02 | 9.58e-22 |
| 15665 | 15450 | serine protease | - | -4.05 | -0.59 | 3.34 | 9.55e-120 | 0.03 | 2.22e-58 |
| 21405 | 21115 | phage tail protein | - | -3.97 | -0.99 | 3.40 | 1.5e-38 | 1.89e-4 | 1.45e-19 |
| 15345 | 15135 | HlyD family type I secretion periplasmic adaptorsubunit | - | -3.95 | -1.33 | 4.09 | 5.53e-35 | 1.7e-5 | 1.42e-22 |
| 23420 | 23100 | DUF3360 domain-containing protein | - | -3.91 | -0.86 | 4.70 | 8.04e-51 | 0.02 | 2.09e-71 |
| 06570 | 06480 | *nlpI* | - | -3.91 | -0.93 | 3.63 | 7.87e-48 | 0.00 | 1.44e-38 |
| 17675 | 17445 | spermidine/putrescine ABC transporter substrate-binding protein | - | -3.91 | -0.42 | 5.69 | 1.27e-6 | 0.59 | 4.39e-9 |
| 20645 | 20370 | MFS transporter | - | -3.90 | -0.67 | 4.72 | 5.02e-7 | 0.35 | 2.08e-7 |
| 20665 | 20390 | DUF1800 domain-containing protein | - | -3.82 | -1.58 | 4.38 | 1.39e-17 | 6.27e-4 | 1.89e-15 |
| 20565 | 20290 | thioredoxin domain-containing protein | - | -3.74 | -1.21 | 4.69 | 5.04e-31 | 2.62e-4 | 1.57e-24 |
| 14445 | 14245 | cation acetate symporter | *actP* | -3.70 | 0.43 | 5.43 | NA | 0.59 | NA |
| 17075 | 16860 | *emrD* | *emrD* | -3.66 | -0.58 | 4.74 | NA | 0.43 | 2.36e-6 |
| 20660 | 20385 | DUF1501 domain- | - | -3.63 | -1.56 | 4.37 | 7.56e-27 | 1.94e-6 | 8.77e-41 |

| | | | | | | | | | |
|---|---|---|---|---|---|---|---|---|---|
| | | containing protein | | | | | | | |
| 17085 | 16870 | *speA* | *speA* | -3.52 | -0.88 | 4.13 | 1.35e-5 | 0.20 | 1.82e-5 |
| 16510 | 16295 | hypothetical protein | *VVA0005* | -3.49 | -2.09 | 0.00 | 8.9e-13 | 4.24e-6 | 1.00 |
| 13185 | 13015 | *uspA* | *uspA* | -3.35 | -2.26 | 3.58 | 9.72e-13 | 6.72e-7 | 4.31e-13 |
| 07635 | 07530 | hypothetical protein | - | -3.32 | -1.89 | 6.11 | 3.74e-36 | 7.3e-11 | 4.08e-64 |
| 15150 | 14940 | *pspG* | *pspG* | -3.29 | -0.75 | 3.50 | 3.16e-5 | 0.29 | 2.05e-4 |
| 11055 | 10905 | Na/Pi cotransporter family protein | *nptA* | -3.26 | -0.12 | 0.02 | 5.9e-9 | 0.89 | 0.98 |
| 17600 | 17375 | hypothetical protein | - | -3.13 | -1.10 | 4.72 | 1.59e-4 | 0.10 | 1.62e-6 |
| 04080 | 04030 | transcriptional regulator | - | -3.11 | -1.57 | 3.60 | 5.47e-8 | NA | NA |
| 19935 | 19670 | murein L, D-transpeptidase catalytic domain family protein | - | -3.09 | -0.89 | 2.18 | 4.83e-24 | 0.00 | 3.44e-12 |
| 07910 | 07805 | sigma-54-dependent Fis family transcriptional regulator | - | -3.08 | 0.46 | 1.34 | 9.08e-28 | 0.19 | 0.00 |
| 18170 | 17935 | hypothetical protein | - | -3.07 | -1.51 | 3.55 | 2.45e-68 | 5.09e-16 | 8.61e-73 |
| 07745 | 07640 | *pspA* | *pspA* | -3.06 | -1.70 | 3.15 | 2.67e-10 | 4.82e-4 | 1.33e-9 |
| 15595 | 15385 | glycine C-acetyltransferase | *kbl* | -3.04 | -1.44 | 3.11 | 5.34e-12 | 0.00 | 2.81e-11 |
| 17520 | 17300 | copper resistance protein NlpE N-terminal domain-containing protein | - | -3.03 | -1.91 | 3.41 | 4.06e-48 | 8.82e-10 | 1.11e-31 |

| | | | | | | | | | |
|---|---|---|---|---|---|---|---|---|---|
| 16600 | 16385 | DUF3081 domain-containing protein | - | -2.99 | -0.81 | 4.24 | 4.83e-7 | 0.24 | 2.37e-11 |
| 17505 | 17285 | glycine zipper 2TM domain-containing protein | - | -2.95 | -1.18 | -0.54 | 3.09e-18 | 0.00 | 0.17 |
| 07740 | 07635 | *pspB* | *pspB* | -2.92 | -1.76 | 2.33 | 2.74e-10 | 1.33e-4 | 6.49e-5 |
| 21030 | 20740 | hypothetical protein | *VPA0857* | -2.89 | -0.66 | 2.91 | 4.7e-4 | 0.36 | 0.00 |
| 17080 | 16865 | *speB* | *speB* | -2.86 | -0.94 | 3.57 | 7.54e-4 | 0.16 | 6.25e-4 |
| 21460 | 21170 | S8 family serine peptidase | - | -2.85 | -1.03 | 3.98 | 3.77e-27 | 0.00 | 5.05e-43 |
| 13475 | 13300 | Hsp20 family protein | *ibpA* | -2.84 | -2.32 | 0.42 | 4.2e-16 | 1.35e-15 | 0.22 |
| 07610 | 07510 | response regulator transcription factor | - | -2.80 | -2.38 | 3.43 | 1.92e-59 | 2.30e-58 | 1.01e-16 |
| 20400 | 20125 | DUF2589 domain-containing protein | - | -2.79 | -1.96 | 5.79 | 1.94e-11 | 6.99e-7 | 2.3e-19 |
| 13625 | 13445 | Bcr/CflA family multidrug efflux MFS transporter | *ydhC* | -2.76 | -0.77 | 2.66 | 3.70e-4 | 0.27 | 0.00 |
| 17500 | 17280 | hypothetical protein | - | -2.76 | -1.46 | 1.94 | 1.35e-6 | 0.01 | 0.00 |
| 00895 | 00875 | DUF2189 domain-containing protein | - | -2.76 | 0.37 | 4.34 | 1.69e-11 | 0.54 | 1.58e-24 |
| 16095 | 15875 | hypothetical protein | - | -2.75 | -1.42 | 1.98 | 5.75e-20 | 9.24e-7 | 4.52e-8 |
| 06555 | 06465 | homocysteine S-methyltransfera | - | -2.75 | -1.04 | 2.87 | 4.69e-29 | 1.62e-4 | 6.27e-27 |

| | | | | | | | | | |
|---|---|---|---|---|---|---|---|---|---|
| | | se family protein | | | | | | | |
| 08935 | 08820 | PLP-dependent cysteine synthase family protein | - | -2.74 | -1.39 | 2.72 | 0.00 | 0.03 | 0.01 |
| 04255 | 04205 | hypothetical protein | - | -2.73 | -1.42 | 3.07 | 7.41e-8 | 0.00 | 3.15e-9 |
| 06220 | 06130 | 2OG-Fe (II) oxygenase | - | -2.72 | -0.56 | 4.18 | 1.89e-6 | 0.45 | 5.43e-11 |
| 17995 | 17760 | beta-ketoacyl-ACP reductase | - | -2.71 | -1.90 | 4.33 | 9.71e-63 | 2.43e-25 | 1.21e-115 |
| 07735 | 07630 | *pspC* | *pspC* | -2.71 | -1.66 | 2.21 | 4.54e-10 | 1.67e-4 | 4.95e-5 |
| 01820 | 01790 | DUF1127 domain-containing protein | - | -2.70 | -1.13 | 5.71 | 8.84e-10 | 0.01 | 1.27e-20 |
| 14440 | 14240 | hypothetical protein | - | -2.69 | 0.16 | 3.25 | NA | NA | NA |
| 15590 | 15380 | *tdh* | *tdh* | -2.67 | -1.62 | 3.09 | 3.89e-13 | 1.52e-5 | 9.42e-16 |
| 16040 | 15820 | LysR family transcriptional regulator | - | -2.66 | -0.92 | 0.20 | 1.93e-7 | 0.08 | 0.75 |
| 21735 | 21440 | *mmsB* | *mmsB* | -2.66 | -1.11 | 4.46 | 1.99e-13 | 0.01 | 3.84e-27 |
| 21290 | 21000 | carboxypeptidase | - | -2.63 | -0.96 | 4.05 | 1.15e-8 | 0.02 | 1.43e-13 |
| 17100 | 16885 | *rimK* | *rimK* | -2.61 | -0.82 | 3.05 | 3.76e-9 | 0.09 | 3.66e-11 |
| 20240 | 19965 | *katG* | *katG* | -2.60 | -1.48 | 5.88 | 4.84e-29 | 3.79e-11 | 2.37e-141 |
| 04830 | 04745 | Gfo/Idh/MocA family oxidoreductase | *iolG* | -2.60 | -1.31 | 2.99 | 6.68e-27 | 3.58e-8 | 5.43e-30 |
| 23145 | 22825 | blaCARB | - | -2.60 | -1.29 | 0.48 | 4.73e-16 | 1.57e-4 | 0.17 |
| 08720 | 08615 | CBS domain-containing protein | - | -2.59 | -1.19 | 3.21 | 5.67e-12 | 0.00 | 1.58e-10 |

| | | | | | | | | | |
|---|---|---|---|---|---|---|---|---|---|
| 06425 | 06335 | hypothetical protein | *NA* | -2.56 | -1.53 | 2.89 | 3.96e-11 | 1.63e-4 | 1.37e-12 |
| 22730 | 22415 | IclR family transcriptional regulator | - | -2.54 | -1.31 | 1.85 | 1.22e-5 | 0.02 | 0.00 |
| 21770 | 21475 | isovaleryl-CoA dehydrogenase | *liuA* | -2.54 | -0.77 | 3.29 | 2.19e-17 | 0.07 | 1.57e-24 |
| 22580 | 22270 | hypothetical protein | - | -2.53 | -0.74 | 3.90 | 3.8e-36 | 8.03e-4 | 4.49e-60 |
| 21775 | 21480 | methylcrotonoyl-CoA carboxylase | *liuB* | -2.53 | -0.62 | 3.75 | 2.69e-12 | 0.22 | 2.67e-24 |
| 24195 | 12810 | hypothetical protein | *NA* | -2.52 | -2.57 | -0.82 | 2.52e-8 | 6.17e-10 | 0.10 |
| 17895 | 17660 | *doeB* | - | -2.51 | -0.75 | 4.51 | 2.75e-15 | 0.06 | 5.5e-39 |
| 21400 | 21110 | sel1 repeat family protein | - | -2.50 | -0.80 | 3.00 | 1.33e-14 | 0.02 | 1.92e-19 |
| 16605 | 16390 | DUF413 domain-containing protein | - | -2.49 | -0.68 | 2.19 | 0.00 | 0.35 | 0.02 |
| 10455 | 10310 | helix-turn-helix domain-containing protein | - | -2.49 | -1.30 | 3.80 | 2.73e-12 | 4.84e-4 | 2.19e-17 |
| 21780 | 21485 | enoyl-CoA hydratase/isomerase family protein | *liuC* | -2.48 | -0.75 | 3.13 | 2.51e-14 | 0.05 | 4.08e-11 |
| 05940 | 05845 | 4Fe-4S dicluster domain-containing protein | *fdhB* | -2.48 | -0.84 | 4.42 | 1.42e-39 | 1.05e-4 | 2.62e-73 |
| 22255 | 21950 | ATP-binding cassette domain-containing protein | - | -2.48 | -1.10 | 4.66 | 9.37e-20 | 3.27e-5 | 1.35e-30 |
| 19150 | 18890 | electron transfer flavoprotein subunit | - | -2.47 | -0.59 | 0.71 | 2.14e-23 | 0.02 | 0.01 |

| | | | | | | | | | |
|---|---|---|---|---|---|---|---|---|---|
| | | beta/FixAfamily protein | | | | | | | |
| 17845 | 17610 | hypothetical protein | *NA* | -2.47 | -1.47 | 2.11 | 1.55e-15 | 6.45e-6 | 6.93e-7 |
| 04285 | 04235 | septum formation initiator | *sdaC* | -2.47 | -1.34 | 1.73 | 5.32e-9 | 0.00 | 1.04e-4 |
| 21760 | 21465 | thiolase family protein | *atoB* | -2.46 | -1.22 | 3.32 | 6.41e-28 | 3.33e-5 | 1.57e-25 |
| 21135 | 20845 | *glgC* | *glgC2* | -2.46 | -0.53 | 3.29 | 5.41e-22 | 0.07 | 7.86e-28 |
| 22945 | 22630 | cytochrome c oxidase assembly protein | NA | -2.45 | 0.20 | 5.17 | 2.88e-5 | 0.78 | 2.75e-12 |
| 22245 | 21940 | hypothetical protein | - | -2.44 | -1.33 | 3.92 | 8.72e-12 | 2.46e-4 | 2.7e-21 |
| 15845 | 15630 | PTS glucose/sucrose transporter subunit IIB | - | -2.43 | -1.72 | 2.99 | 2.73e-12 | 2.16e-6 | 5.68e-16 |
| 07560 | 07460 | NapC/NirT family cytochrome c | - | -2.43 | -1.25 | 5.25 | 2.13e-14 | 7.67e-4 | 6.57e-57 |
| 20230 | 19955 | YbaK/EbsC family protein | - | -2.42 | -1.01 | 5.67 | 3.97e-9 | 0.03 | 1.11e-35 |
| 23110 | 22795 | *fdhF* | - | -2.42 | -1.68 | 3.34 | 2.14e-9 | 2.32e-5 | 1.67e-16 |
| 07570 | 07470 | OmcA/MtrC family decaheme c-type cytochrome | - | -2.42 | -1.35 | 4.44 | 3.74e-17 | 4.29e-5 | 3.76e-18 |
| 03240 | 03190 | SDR family oxidoreductase | *VP2120* | -2.41 | -2.06 | 1.89 | 1.73e-10 | 3.15e-15 | 0.00 |
| 21755 | 21460 | CoA-acylating methylmalonate-semialdehyde dehydrogenase | *mmsA* | -2.41 | -1.24 | 3.58 | 1.8e-10 | 0.00 | 7.29e-18 |
| 17095 | 16880 | ATP-dependent zinc protease | - | -2.41 | -0.72 | 1.11 | 4.82e-7 | 0.17 | 0.29 |

| | | | | | | | | | |
|---|---|---|---|---|---|---|---|---|---|
| 20855 | 20565 | universal stress protein | - | -2.41 | -0.99 | 3.82 | 8.74e-10 | 0.02 | 3.46e-17 |
| 19215 | 18955 | mechanosensitive ion channel | - | -2.39 | -0.81 | 3.54 | 5.26e-23 | 0.01 | 1.28e-16 |
| 05555 | 05480 | *yccX* | *acyP* | -2.39 | -1.03 | 3.36 | 7.99e-9 | 0.04 | 5.79e-14 |
| 20640 | 20365 | nitrous oxide-stimulated promoter family protein | - | -2.37 | -1.07 | 2.74 | 9.81e-6 | 0.06 | 1.96e-6 |
| 05945 | 05850 | formate dehydrogenase subunit alpha | *fdhA* | -2.37 | -0.79 | 4.06 | 8.28e-33 | 0.00 | 2.61e-64 |
| 07825 | 07720 | TIGR02647 family protein | - | -2.37 | -1.25 | 1.50 | 1.17e-12 | 4.61e-4 | 0.00 |
| 02845 | 02800 | *fadI* | *fadI* | -2.35 | -1.93 | 2.63 | 1.86e-28 | 4.3e-19 | 4.13e-25 |
| 21595 | 21300 | alanine:cation symporter family protein | NA | -2.35 | -1.23 | 3.73 | 1.79e-18 | 8.98e-5 | 9.27e-32 |
| 18830 | 18580 | *pobA* | *pobA* | -2.32 | 0.21 | 2.319 | 3.36e-11 | 0.73 | 6.16e-10 |
| 19145 | 18885 | electron transfer flavoprotein subunit alpha | - | -2.32 | -0.63 | 1.37 | 4.19e-13 | 0.08 | 1.03e-4 |
| 23475 | 23155 | Hsp20/alpha crystallin family protein | - | -2.32 | -1.42 | 4.12 | 0.00 | 0.026 | 9.2e-6 |
| 18060 | 17825 | YccF domain-containing protein | *yccF* | -2.31 | -1.48 | 2.43 | 2.49e-45 | 1.89e-17 | 2.67e-12 |
| 16100 | 15880 | hypothetical protein | NA | -2.31 | -1.40 | 1.08 | 1.08e-14 | 5.88e-7 | 0.03 |
| 22940 | 22625 | cytochrome c oxidase subunit 3 | *coxC* | -2.31 | 0.02 | 5.16 | 4.34e-5 | 0.98 | 7.51e-14 |
| 05505 | 05425 | c-type cytochrome | - | -2.31 | -0.93 | 1.99 | 6.24e-8 | 0.00 | 2.6e-8 |
| 21750 | 21455 | acyl-CoA dehydrogenase family protein | - | -2.27 | -1.14 | 3.61 | 8.16e-14 | 8.34e-4 | 1.86e-26 |

| | | | | | | | | | |
|---|---|---|---|---|---|---|---|---|---|
| 22205 | 21900 | AI-2E family transporter | - | -2.27 | -0.72 | 3.78 | 3.2e-11 | 0.07 | 4.38e-14 |
| 16045 | 15825 | ectoine synthase | *ectC* | -2.27 | -0.74 | -0.47 | 2.11e-7 | 0.08 | 0.24 |
| 18320 | 18075 | *napA* | *napA* | -2.26 | -1.05 | 3.25 | 1.65e-12 | 0.00 | 4.89e-21 |
| 21765 | 21470 | MerR family DNA-binding transcriptional regulator | - | -2.25 | -0.71 | 3.77 | 4.49e-14 | 0.05 | 2.25e-23 |
| 05950 | 05855 | transcriptional initiation protein Tat | - | -2.25 | -0.54 | 4.29 | 2.02e-11 | 0.13 | 4.85e-27 |
| 21740 | 21445 | enoyl-CoA hydratase/isomerase family protein | - | -2.25 | -0.87 | 3.99 | 7.29e-18 | 0.01 | 6.8e-41 |
| 04190 | 04140 | TRAP transporter substrate-binding protein | - | -2.25 | -0.61 | 7.74 | 6.59e-17 | 0.03 | 4.27e-201 |
| 07565 | 07465 | c-type cytochrome | NA | -2.24 | -1.38 | 5.65 | 5.16e-13 | 3.41e-5 | 1.79e-71 |
| 16125 | 15905 | phosphotransferase | - | -2.24 | -0.57 | 1.09 | 2.49e-20 | 0.01 | 8.48e-5 |
| 17890 | 17655 | Lrp/AsnC family transcriptional regulator | - | -2.24 | -0.69 | 2.98 | 2.2e-10 | 0.06 | 2.65e-10 |
| 17985 | 17750 | YdcH family protein | *VPA0402* | -2.22 | -1.19 | 4.81 | 9.75e-25 | 2.23e-6 | 6.67e-90 |
| 18315 | 18070 | nitrate reductase cytochrome c-type subunit | *napB* | -2.21 | -0.98 | 3.67 | 2.07e-9 | 0.02 | 8.08e-22 |
| 22260 | 21955 | MlaE family lipid ABC transporter permease subunit | - | -2.21 | -1.32 | 4.20 | 5.2e-17 | 2.49e-6 | 1.46e-23 |
| 15505 | 15295 | *napG* | *napG* | -2.21 | -1.25 | 2.92 | 1.92e-5 | 0.01 | 8.13e-8 |

| | | | | | | | | | |
|---|---|---|---|---|---|---|---|---|---|
| 20405 | 20130 | hypothetical protein | - | -2.21 | -1.74 | 3.62 | 2.19e-10 | 1.1e-7 | 2.9e-20 |
| 00945 | 00925 | oxidative stress defense protein | - | -2.20 | -1.14 | 0.04 | 3.42e-19 | 5.45e-5 | 0.91 |
| 13655 | 13475 | YifB family Mg chelatase-like AAA ATPase | *comM* | -2.19 | -1.77 | 3.19 | 1.77e-22 | 3.31e-16 | 2.88e-40 |
| 17545 | 17325 | elongation factor G | - | -2.19 | -1.89 | 5.30 | 9.83e-10 | 3.77e-7 | 2.07e-30 |
| 17720 | 17490 | acyl-CoA synthetase | - | -2.18 | 0.31 | 2.36 | 7.14e-6 | 0.64 | 1.58711e-4 |
| 02850 | 02805 | *fadJ* | *fadJ* | -2.18 | -1.54 | 2.57 | 8.6e-20 | 1.36e-11 | 3.69e-25 |
| 19050 | 18790 | NAD(P)-dependent oxidoreductase | - | -2.17 | -1.12 | 1.70 | 4.66e-12 | 3.49e-4 | 8.77e-6 |
| 18535 | 18285 | alpha/beta hydrolase | - | -2.17 | -1.16 | 4.19 | 5.45e-20 | 3.97e-6 | 7.53e-62 |
| 17980 | 17745 | DUF4382 domain-containing protein | - | -2.17 | -1.25 | 4.05 | 7.75e-28 | 4.83e-10 | 1.75e-82 |
| 19155 | 18895 | electron transfer flavoprotein-ubiquinone oxidoreductase | - | -2.16 | -0.82 | 2.25 | 2.19e-9 | 0.03 | 3.1e-8 |
| 07615 | 07515 | GHKL domain-containing protein | *phoQ* | -2.16 | -1.64 | 3.27 | 3.97e-36 | 5.65e-14 | 1.99e-51 |
| 19310 | 19050 | thiolase family protein | - | -2.16 | -0.90 | 2.56 | 5.08e-17 | 0.00 | 3.38e-14 |
| 17510 | 17290 | DMT family transporter | - | -2.16 | -0.30 | -0.09 | 1.31e-4 | 0.67 | 0.90 |
| 22935 | 22620 | DUF2909 domain-containing protein | - | -2.16 | 0.26 | 4.91 | 9.34e-5 | 0.72 | 8.32e-14 |

| | | | | | | | | | |
|---|---|---|---|---|---|---|---|---|---|
| 08140 | 08035 | S8 family serine peptidase | - | -2.14 | -0.52 | 2.59 | 6.25e-22 | 0.08 | 9.09e-21 |
| 07605 | 07505 | membrane protein | - | -2.13 | -2.18 | 3.94 | 2.34e-12 | 5.89e-40 | 4.34e-77 |
| 15670 | 15455 | hypothetical protein | - | -2.12 | -0.14 | 3.50 | 5.63e-11 | 0.72 | 2.24e-16 |
| 17975 | 17740 | hypothetical protein | - | -2.12 | -1.01 | 2.87 | 5.49e-9 | 0.01 | 2.94e-12 |
| 19100 | 18840 | *amrB* | - | -2.12 | -1.05 | 3.36 | 9.07e-16 | 6.73e-4 | 2.03e-25 |
| 17825 | 17590 | *xdhB* | *xdhB* | -2.11 | -1.05 | 3.03 | 1.49e-7 | 0.02 | 3.26e-13 |
| 07580 | 07480 | MtrB/PioB family decaheme-associated outer membrane protein | - | -2.10 | -1.17 | 5.44 | 1.48e-9 | 0.00 | 2.3e-54 |
| 18580 | 18330 | LysR family transcriptional regulator | NA | -2.09 | -0.12 | 2.16 | 6.21e-17 | 0.76 | 1.43e-14 |
| 21745 | 21450 | enoyl-CoA hydratase | - | -2.09 | -0.89 | 3.30 | 5.34e-12 | 0.01 | 2.66e-19 |
| 17815 | 17580 | 2-oxo-4-hydroxy-4-carboxy-5-ureidoimidazoline decarboxylase | - | -2.09 | -0.44 | 3.33 | 5.88e-5 | 0.46 | 1.09e-7 |
| 12955 | 12785 | *add* | *add* | -2.08 | -0.74 | 0.98 | 1.29e-8 | 0.07 | 0.01 |
| 21325 | 21035 | hypothetical protein | - | -2.08 | -1.20 | 3.49 | 1.47e-8 | 9.35e-4 | 1.87e-17 |
| 05960 | 05865 | 4Fe-4S binding protein | - | -2.07 | -0.69 | 3.75 | 5.39e-15 | 0.01 | 4.05e-32 |
| 19095 | 18835 | *amrS* | - | -2.07 | -1.00 | 3.69 | 7.42e-6 | 0.04 | 1.23e-13 |
| 17900 | 17665 | *doeA* | - | -2.07 | -0.79 | 4.60 | 7.08e-15 | 0.02 | 3.36e-61 |
| 19105 | 18845 | *amrA* | - | -2.06 | -0.25 | 2.84 | 4.84e-15 | 0.53 | 7.17e-14 |

| | | | | | | | | | |
|---|---|---|---|---|---|---|---|---|---|
| 15340 | 15130 | mechanosensitive ion channel family protein | - | -2.06 | -0.86 | 1.78 | 9.54e-20 | 0.00 | 4.42e-10 |
| 17810 | 17575 | *uraH* | - | -2.05 | -0.90 | 3.44 | 1.22e-4 | 0.10 | 4.42e-8 |
| 06085 | 05995 | CPXCG motif-containing cysteine-rich protein | - | -2.03 | -0.89 | 1.28 | 1.78e-5 | 0.07 | 0.06 |
| 15580 | 15370 | cold-shock protein | *cspA* | -2.03 | -0.77 | 2.47 | 6.22e-19 | 1.65e-4 | 1.53e-16 |
| 19165 | 18905 | long-chain fatty acid--CoA ligase | - | -2.03 | -0.37 | 3.06 | 2.55e-5 | 0.52 | 6e-9 |
| 21205 | 20915 | *fruA* | *fruA* | -2.02 | -1.04 | 2.72 | 3.31e-14 | 5.62e-4 | 3.3e-22 |
| 05935 | 05840 | formate dehydrogenase subunit gamma | *fdnI* | -2.02 | -0.84 | 4.11 | 6.2e-14 | 1.21e-4 | 1.6e-79 |
| 21035 | 20745 | TerC family protein | *alx* | -2.02 | -0.95 | 3.31 | 4.6e-4 | 0.11 | 7.1e-8 |
| 23870 | NA | hypothetical protein | - | -2.01 | -1.52 | 1.94 | NA | NA | NA |
| 07575 | 07475 | DmsE family decaheme c-type cytochrome | - | -2.01 | -1.22 | 5.45 | 6.34e-12 | 1.67e-4 | 3.17e-66 |
| 01835 | 01810 | prepilin peptidase | - | -2.01 | 0.18 | 3.84 | 3.14e-31 | 0.50 | 3.63e-61 |
| 19540 | 19285 | ExeM/NucH family extracellular endonuclease | - | -2.00 | -1.61 | 3.02 | 7.21e-9 | 2.57e-6 | 1.2e-14 |
| 23025 | 22710 | GFA family protein | NA | -2.00 | -0.95 | 3.22 | 1.42e-8 | 0.01 | 7.72e-14 |
| 08480 | 08375 | porin | - | -2.00 | -0.45 | -1.78 | 2.89e-18 | 0.04 | 4.79e-16 |

**Table S8. List of strains used in this study.** Detailed description of the plasmids is listed in Table S5.

| **Internal strain name** | **Plasmid** | **Genotype** | **Remarks** | **Reference or source** |
| --- | --- | --- | --- | --- |
| GFN22 | | *V. natriegens* DSMZ 759 ΔVNP1+2 Δ*dns* | Prophage knockout strain published by Pfeifer et al. (2019) with additional exonuclease deletion (Δ*dns*), used as ancestral "WT" strain in this study | Pfeifer et al. 2019, this study |
| GFN135 | | GFN22 Δ*acsA1* | Δ*acsA1* deletion phenotype verification strain | This study |
| GFN133 | | GFN22 Δ*acsA2* | Δ*acsA2* deletion phenotype verification strain | This study |
| GFN163 | | GFN22 Δ*acsA1* Δ*acsA2* | Δ*acsA1* Δ*acsA2* double deletion phenotype verification strain | This study |
| GFN36 | | GFN22 Δ*pta1* | Δ*pta1* deletion phenotype verification strain | This study |
| GFN38 | | GFN22 Δ*pta2* | Δ*pta2* deletion phenotype verification strain | This study |
| GFN175 | | GFN22 Δ*pta1* Δ*pta2* | Δ*pta1* Δ*pta2* double deletion phenotype verification strain | This study |
| GFN40 | | GFN22 Δ*ackA1* | Δ*ackA1* deletion phenotype verification strain | This study |
| GFN137 | | GFN22 Δ*ackA2* | Δ*ackA2* deletion phenotype verification strain | This study |
| GFN172 | | GFN22 Δ*ackA1* Δ*ackA2* | Δ*ackA1* Δ*ackA2* double deletion phenotype verification strain | This study |
| GFN52 | | GFN22 AE1 rep 1 | GFN22 adapted to 122 mM NaOAc for 1000 generations | This study |
| GFN53 | | GFN22 AE1 rep 2 | GFN22 adapted to 122 mM NaOAc for 1000 generations | This study |
| GFN48 | | GFN22 AE2 rep 1 | GFN22 adapted to 122 mM NaOAc for 1000 generations | This study |
| GFN49 | | GFN22 AE2 rep 2 | GFN22 adapted to 122 mM NaOAc for 1000 generations | This study |
| GFN50 | | GFN22 AE3 rep 1 | GFN22 adapted to 122 mM NaOAc for 1000 generations | This study |
| GFN51 | | GFN22 AE3 rep 2 | GFN22 adapted to 122 mM NaOAc for 1000 generations | This study |
| GFN97 | | GFN22 Δ*bccT1* | Δ*bccT1* deletion phenotype verification strain | This study |
| GFN239 | | GFN22 Δ*ect* | Δ*ect* deletion phenotype verification strain | This study |

| | | | | |
|---|---|---|---|---|
| GFN241 | | GFN22 Δ*bccT1* Δ*ect* | Δ*bccT1* Δ*ect* deletion phenotype verification strain | This study |
| GFN143 | | GFN22 Δ*luxU* | Δ*luxU* deletion phenotype verification strain | This study |
| GFN44 | | GFN22 Δ*vps* | Δ*cps* deletion phenotype verification strain | This study |
| GFN159 | | AE2 Δ*acsA1* | Δ*acsA1* deletion phenotype verification strain after ALE | This study |
| GFN157 | | AE2 Δ*acsA2* | Δ*acsA2* deletion phenotype verification strain after ALE | This study |
| GFN165 | | AE2 Δ*acsA1* Δ*acsA2* | Δ*acsA1* Δ*acsA2* double deletion phenotype verification strain after ALE | This study |
| GFN155 | | AE2 Δ*ackA1* | Δ*pta1* deletion phenotype verification strain after ALE | This study |
| GFN147 | | AE2 Δ*ackA2* | Δ*pta2* deletion phenotype verification strain after ALE | This study |
| GFN173 | | AE2 Δ*ackA1* Δ*ackA2* | Δ*pta1* Δ*pta2* double deletion phenotype verification strain after ALE | This study |
| GFN153 | | AE2 Δ*pta1* | Δ*ackA1* deletion phenotype verification strain after ALE | This study |
| GFN167 | | AE2 Δ*pta2* | Δ*ackA2* deletion phenotype verification strain after ALE | This study |
| GFN169 | | AE2 Δ*pta1* Δ*pta2* | Δ*ackA1* Δ*ackA2* double deletion phenotype verification strain after ALE | This study |
| GFN1-103 | pRP1_008 | GFN22 | Inducible PHB production strain | This study |
| GFN1-107 | pRP1_008 | AE2 | Inducible PHB production strain after ALE | This study |
| GFN1-108 | pRP1_008 | AE3 | Inducible PHB production strain after ALE | This study |
| GFN1-126 | pRP1_037 | GFN22 | Empty plasmid control strain | This study |
| GFN1-158 | pRP1_048 | GFN22 | $P_{vps}$-*lux* (*cps*) biosensor strain | This study |
| GFN1-160 | pRP1_048 | GFN22 Δ*luxU* | $P_{vps}$-*lux* (*cps*) biosensor strain with Δ*luxU* deletion | This study |
| GFN1-109 | pRP1_008 | AE2 Δ*phaY* | Inducible PHB production strain for Δ*phaY* deletion verification (Supplementary Fig. S6) | This study |
| GFN1-148 | pRP1_008 | AE2 Δ*phaY2* | Inducible PHB production strain for Δ*phaY2* deletion verification (Supplementary Fig. S6) | This study |

| | | | | |
|---|---|---|---|---|
| GFN1-150 | pRP1_008 | AE2 Δ*phaY* Δ*phaY2* | Inducible PHB production strain for Δ*phaY*Δ*phaY2* double deletion verification (Supplementary Fig. S6) | This study |

**Table S9. Primers used in this study for the generation of the tDNA used in the genome engineering process.** All sequences written in the 5’ -> 3’ direction.

| Part | Forward oligonucleotide | | Reverse oligonucleotide | | Synthesis method | Homology flank size |
|---|---|---|---|---|---|---|
| *ackA1* tDNA 5' homology | GF1321 | CGACAAGTCAGAAAGTCCAGTC | GF1314 | CATTAAAGACCTGCTAGAACTAGCTTAGACATGTATGACTACC | fusion PCR | 1619 |
| *ackA1* tDNA 3' homology | GF1315 | CTAAGCTAGTTCTAGCAGGTCTTTAATGCATAAACTG | GF1316 | CTACAGTTTTCCCATCTGTCTCG | fusion PCR | 3049 |
| *ackA2* tDNA 5' homology | GF1888 | CGAATAACTGCTTCAAAACCCACC | GF1889 | GACTGTTGAGCGTGACGAGCTTCCTGAGTTGATC | fusion PCR | 2393 |
| *ackA2* tDNA 3' homology | GF1890 | CAGGAAGCTCGTCACGCTCAACAGTCGGTCGAG | GF1891 | CAAAACCTCGTCCATGCTCTC | fusion PCR | 2547 |
| *acsA1* tDNA 5' homology | GF1882 | CGAGAATTACTCACTCCCGTC | GF1883 | CGGCAAGCGTTGAGTAAGTTTCATTATCCGCGTGTG | fusion PCR | 1502 |
| *acsA1* tDNA 3' homology | GF1884 | CGGATAATGAAACTTACTCAACGCTTGCCGACCCTAG | GF1885 | ACTAGAGGTTGGTCGAATTCAACTG | fusion PCR | 1323 |
| *acsA2* tDNA 5' homology | GF1874 | CGCAGACACCATTCTATTGCTTG | GF1875 | CGAAATATCACCTTCATTGTCCGCTAGCGTTTGGTG | fusion PCR | 1516 |
| *acsA2* tDNA 3' homology | GF1877 | CGCTAGCGGACAATGAAGGTGATATTTCGACTTTGGAG | GF1878 | GACCTTCTTCATTGAGCAGCATG | fusion PCR | 1593 |
| *pta1* tDNA 5' homology | GF1354 | GGTCACTATCTGCAAGGAACG | GF1322 | GCTTGCGCATACCCATAATAGTACGAGACATTCGTAGAG | fusion PCR | 1531 |
| *pta1* tDNA 3' homology | GF1323 | CGTACTATTATGGGTATGCGCAAGCCTGTAAAC | GF1324 | GACCGTGTTCTGGTTATGTATTTGG | fusion PCR | 2547 |
| *pta2* tDNA 5' homology | GF1287 | GATTCGTGCTCTCGGTTGCG | GF1292 | GCCTTTCACTTGTTCCATCGCCTTCATACTATAAATTCC | fusion PCR | 1149 |

| | | | | | | |
|---|---|---|---|---|---|---|
| *pta2* tDNA 3' homology | GF1293 | GAAGGCGATGGAACAAGTGAAAGGCCAGTCTAACG | GF1346 | CGAGTTATGCAGACACCTTCAAC | fusion PCR | 3097 |
| *luxU* tDNA 5' homology | GF1911 | GCCAAATTCTTAACTCATTGATCTCTG | GF1912 | GTATCGCGCAGAACCAATCTCCATGGACAGATCGTC | fusion PCR | 1588 |
| *luxU* tDNA 3' homology | GF1913 | CCATGGAGATTGGTTCTGCGCGATACACAAGAG | GF1914 | CTCGATACCATTCATCACTTCGG | fusion PCR | 1772 |
| *bccT1* tDNA plasmid 5' homology | GF1559 | AAAAGGTCTCGCTCGAACAGATACATCATACTCAGTGCACGTAG | GF1560 | AAAAGGTCTCCCTCACTCCCGCAACGAATATTAACATCAGTCAC | Golden gate | 1359 |
| *bccT1* tDNA plasmid 3' homology | GF1561 | AAAAGGTCTCACTCGCGCTGTTTAAGAACCGAACTACCTGCATAC | GF1562 | AAAAGGTCTCCCTCAAGCTATAGTCACCCGCAGAGTAAGG | Golden gate | 1087 |
| *ect* tDNA plasmid 5' homology | GF2756 | CCTCAAATGGGTGCATGATATCG | GF2759 | CGTCTTGTTGTGTAATTGACGTGAAAATTCATATCCTTTGAGGTCTAATTA | fusion PCR | 2085 |
| *ect* tDNA plasmid 3' homology | GF2758 | AAAGGATATGAATTTTCACGTCAATTACACAACAAGACGTGATAC | GF2757 | CGACACAAATTACTGCGCTTTTTAATC | fusion PCR | 1979 |
| *phaY* tDNA 5' homology | GF1413 | AAAAGGTCTCACTCGCAGACACATTAAACGGCGTG | GF1414 | AAAAGGTCTCTAACCAATCCAACGTAACTCACTAATC | Golden gate | 2124 |
| *phaY* tDNA 3' homology | GF1415 | AAAAGGTCTCTGGTTGTCCGCTGGCAGGATAATACG | GF1416 | AAAACGTCTCTCTCAATGCCTCTAAAAAATAATAAGCTGATTATGAC | Golden gate | 1717 |
| *phaY2* tDNA 5' homology | GF1901 | CAAAGCCCGGTGACTTTTATGC | GF1902 | AAATCTTTAGCGCAACGGCTCTGAAGGAGTATTACGGTG | fusion PCR | 1584 |
| *phaY2* tDNA 3' homology | GF1903 | TAATACTCCTTCAGAGCCGTTGCGCTAAAGATTTGATGTC | GF1904 | CCTAGTTTTGAATGCCTAGCTTC | fusion PCR | 1533 |
| *vps* tDNA 5' homology | GF1548 | AAAAGGTCTCGCTCGAACAGGAACAATCTTAGAAACTGCACC | GF1549 | AAAAGGTCTCCCTCACTCCCTGCTTCATGAGAACGAATAAGTCC | fusion PCR | 1892 |
| *vps* tDNA 3' homology | GF1550 | AAAAGGTCTCACTCGCGCTGCATACGAAATCACTAAGCTGTCG | GF1551 | AAAAGGTCTCCCTCAAGCTCACTTATAGGGTTGTTCTCTTACTGC | fusion PCR | 1978 |

**Table S10. Oligonucleotides used to assemble the gRNA sequences for the NT-CRISPR plasmid.** Forward and reverse oligonucleotides were annealed and cloned into pST_116 plasmid according to Stukenberg et al. (2022)[21]. All sequences written in the 5’ -> 3’ direction.

| **Target** | **LT (PN96_RS)** | **LT (PN96_)** | **gRNA sequence** | **Forward oligonucleotide** | | **Reverse oligonucleotide** | | **Final Plasmid name** |
|---|---|---|---|---|---|---|---|---|
| *ackA1* | 03415 | 03365 | GGTCACCGTGTAGTACACGGCGG | GF1317 | GTCCGGTCACCGTGTAGTACACGGCGG | GF1318 | AAACCCGCCGTGTACTACACGGTGACC | pST_116_ackA1 |
| *ackA2* | 21805 | 21510 | CGGTGAAAAATTCACAACGA | GF1456 | GTCCCGGTGAAAAATTCACAACGA | GF1457 | AAACTCGTTGTGAATTTTTCACCG | pST_116_ackA2 |
| *acsA1* | 14410 | 14210 | GCAAACGTCAGAAACACTTG | GF1880 | GTCCGCAAACGTCAGAAACACTTG | GF1881 | AAACCAAGTGTTTCTGACGTTTGC | pST_116_acsA1 |
| *acsA2* | 21410 | 21120 | CTGAACACTCTGCTACCACG | GF1872 | GTCCCTGAACACTCTGCTACCACG | GF1873 | AAACCGTGGTAGCAGAGTGTTCAG | pST_116_acsA2 |
| *pta1* | 03410 | 03360 | CGGTCCTCTACAGTACGACGCGG | GF1327 | GTCCCGGTCCTCTACAGTACGACGCGG | GF1328 | AAACCCGCGTCGTACTGTAGAGGACCG | pST_116_pta1 |
| *pta2* | 22305 | 22000 | CCGATGTGGTTCGTACGGCTTTA | GF1303 | GTCCCCGATGTGGTTCGTACGGCTTTA | GF1304 | AAACTAAAGCCGTACGAACCACATCGG | pST_116_pta2 |
| *luxU* | 03340 | 03290 | TATTGATAAGAAAGCGAAAG | GF1909 | GTCCTATTGATAAGAAAGCGAAAG | GF1910 | AAACCTTTCGCTTTCTTATCAATA | pST_116_luxU |
| *bccT1* | 06115 | 06025 | GTCGTTCGTGGCATCGATGG | GF1557 | GTCCGTCGTTCGTGGCATCGATGG | GF1558 | AAACCCATCGATGCCACGAACGAC | pST_116_bccT1 |
| *ectB* | 05300 | 05220 | GTACTTCTTGAAACAGTACA | GF275 | GTCCGTACTTCTTGAAACAGTACA | GF2753 | AAACTGTACTGTTTCAAGAAGTAC | PST_116_ectB |
| *phaY* | 21480 | 21190 | CAGAATCCTAGTAAAGACTGGGG | GF1341 | GTCCCAGAATCCTAGTAAAGACTGGGG | GF1342 | AAACCCCCAGTCTTTACTAGGATTCTG | pST_116_phaY |
| *phaY2* | 22040 | 21745 | TCTCCCCACATGATTAACGT | GF1899 | GTCCTCTCCCCACATGATTAACGT | GF1900 | AAACACGTTAATCATGTGGGGAGA | pST_116_phaY2 |
| *vps* | 15195 | 14985 | CCAAGAAGATGACTACACCG | GF1374 | GTCCCCAAGAAGATGACTACACCG | GF1375 | AAACCGGTGTAGTCATCTTCTTGG | pST_116_vps |

**Table S11. Oligonucleotides used for the verification of correct deletion through colony PCR and sanger sequencing.** Locus tags were derived from Refseq (PN96_RS) and Genbank (PN96_). All sequences written in the 5' -> 3' direction.

| **LT (PN96 _RS)** | **LT (PN96_)** | | **Primer outside homology flank** | | **Sequencing primer** | |
|---|---|---|---|---|---|---|
| | | **Loci** | **Name** | **Sequence** | **Name** | **Sequence** |
| 03415 | 03365 | *ackA1* | GF1313 | CAGCTAAGCCGACCACAATG | GF1354 | GGTCACTATCTGCAAGGAACG |
| 21805 | 21510 | *ackA2* | GF1892 | GTTGACATGAATTGGAAGCTTGATG | GF1893 | GGGTGCTATTCACTTATTTGTTGG |
| 14410 | 14210 | *acsA1* | GF1886 | AGTGATGGTCAACAATCCTAAGTG | GF1887 | CAGCAATTTACTGGTCTAATCTCC |
| 21410 | 21120 | *acsA2* | GF1878 | CCAAGTTTATAAGCAGCCTCCC | GF1879 | CTTTATTAAGTGCAGACATGACACC |
| 03410 | 03360 | *pta1* | GF1321 | CGACAAGTCAGAAAGTCCAGTC | GF1355 | GTAACCAGCTCAGTAAACTCGC |
| 22305 | 22000 | *pta2* | GF1345 | GATTTACTATGGCTGCTGTTTAACG | GF1291 | ATCCTTGTAGAGATATGAACTCGTC |
| 03340 | 03290 | *luxU* | GF1915 | CAGTCGTTTCTATTGGCTTTATTTG | GF1916 | CGTCATGACCGTAATAAACTCAAG |
| 06115 | 06025 | *bccT1* | GF1595 | GCCTCCCTTGTATTTGCAGTG | GF1596 | CCACTGGTGTCGCATCTTCTC |
| 05300 | 05220 | *ect* | GF2754 | GATTGGAACTCGAATCGCCTG | GF2755 | GTTTGGGTTTTTGGCTCGGTTAC |
| 21480 | 21190 | *phaY* | GF1337 | CGTCAAGCTGGTTGAGATGC | GF1358 | CAACACTCCACCACAACCAC |
| 22040 | 21745 | *phaY2* | GF1905 | GCCTTTATTCTTACCTCAGTGCTAC | GF1906 | CCAAAAACGTTTATGAAGTTATTAGAGC |
| 15190-15240 | 14980-15030 | *vps* | GF1594 | GGTGTGATAATGTGATTTCCGCC | GF1377 | CTTTCTGACTCATCTCGTATCGTG |

**Table S12. Oligonucleotides used to create Marburg Collection-compatible genetic parts in this study.** All sequences written in the 5' -> 3' direction. The amplicons were purified and cloned into pMC_V_01 from the Marburg Collection[20].

| **Internal cryostock name** | **Internal plasmid name** | **Description** | **Forward oligonucleotide** | | **Reverse oligonucleotide** | |
|---|---|---|---|---|---|---|
| GFMC0-362 | pRP0_001 | Lvl 0 *phaBAPC* CDS | GF1403 | AACGTCTCGCTCGAATGAATAAAGTCGCTTTGATCACC | GF1404 | AACGTCTCCCTCAAAGCAGGTCATTTCACAGGCAATACC |
| GFMC0-351 | pRP0_002 | Lvl 0 $P_{vps}$ Vn | GF1586 | AAAACGTCTCGCTCGGGAGTTCATCAAAGCCACTCATAGCAATAAG | GF1587 | AAAACGTCTCCCTCAAGTAAGCTACCGCCAATGGAGTTATTG |

**Table S13. Plasmids used in this study with their parts.** The parts used in this study were mainly from the Marburg Collection[20].

| **Internal cryostock name** | **Internal plasmid name** | **Description** | **Parts** |
|---|---|---|---|
| GFMC1-009 | pRP1_008 | ATc-inducible PHB production plasmid | pMC0_8_18_Akan(Vn) (sfgfp)(Vn), pMC0_7_04_OpMB1-M, pMC0_1_02_5C1CLN, pMC0_2_23_Ptet, pMC0_3_03_RB0030, pRP0_001, pMC0_5_03_TB0015, pMC0_6_16_3C5CSN |
| GFMC1-085 | pRP1_037 | Empty plasmid control | pMC0_8_18_Akan(Vn) (sfgfp)(Vn), pMC0_7_04_OpMB1-M, pMC0_1-6_02_1-6 Connector |
| GFMC1-104 | pRP1_048 | $P_{vps}$-*lux* biosensor plasmid | pMC0_8_18_Akan(Vn) (sfgfp)(Vn), pMC0_7_04_OpMB1-M, pMC0_1_02_5C1CLN, pRP0_002, pMC0_3_09_RB0064, pMC0_4_02_CDSlux Operon, pMC0_5_03_TB0015, pMC0_6_16_3C5CSN |
| GFMC2-012 | pST_116 | NT-CRISPR single gRNA entry vector | Described in Stukenberg et al. (2022)[21] |
